# The role of maternal pioneer factors in predefining first zygotic responses to inductive signals

**DOI:** 10.1101/306803

**Authors:** George E. Gentsch, Thomas Spruce, Nick D. L. Owens, James C. Smith

**Author notes:** Correspondence should be addressed to G.E.G or J.C.S.

## Abstract

Embryonic development yields many different cell types in response to just a few families of inductive signals. The property of a signal-receiving cell that determines how it responds to such signals, including the activation of cell type-specific genes, is known as its competence. Here, we show how maternal factors modify chromatin to specify initial competence in the frog *Xenopus tropicalis*. We identified the earliest engaged regulatory DNA sequences, and inferred from them critical activators of the zygotic genome. Of these, we showed that the pioneering activity of the maternal pluripotency factors Pou5f3 and Sox3 predefines competence for germ layer formation by extensively remodeling compacted chromatin before the onset of signaling. The remodeling includes the opening and marking of thousands of regulatory elements, extensive chromatin looping, and the co-recruitment of signal-mediating transcription factors. Our work identifies significant developmental principles that inform our understanding of how pluripotent stem cells interpret inductive signals.

## INTRODUCTION

The specification of different cell types during embryonic development is achieved through the repeated use of a small number of highly conserved intercellular signals. The property of a cell that defines the way it responds to such signals (if it responds at all) is known as its competence^1^. Classical experiments with amphibian embryos show that competence is regulated in both space and time. For example, head ectoderm of the tailbud embryo responds to the underlying optic vesicle by forming a lens, while other surface ectoderm is unable to do so^2^. Similarly, on a temporal scale, naïve pluripotent (animal cap) tissue of the early *Xenopus* embryo loses the ability to form muscle in response to Nodal signaling by the mid-gastrula stage^3^.

How is competence regulated at the molecular level? A simple mechanism would involve the acquisition or loss of components of a signal transduction pathway, but this fails to explain differences in context-dependent responses to the same signals. In addition, although there are some tissue-specific signal receptor isoforms, transduction to the nucleus is limited to just a few signal mediators with transcriptional activity such as β-catenin (canonical Wnt signaling), Smad2 (Nodal signaling) and Smad1 (BMP signaling). Thus, a more plausible and common way in which tissue-specific competence might arise is through the recruitment of these signal mediators to different gene regulatory sites which in turn would drive the specification of different cell types.

A series of experiments indicates that cell lineage determinants such as sequence-specific transcription factors (TFs) may play a role in determining competence. For lens induction, the responding cells of the head ectoderm depend on the presence of the eye-regulating TF Pax6^4^. However, although we know that certain TFs can act as competence factors, these TFs have not been identified in a systematic fashion, and their modes of action remain largely unknown. Our understanding of how chromatin interprets inductive signals is especially important because the generation of therapeutically relevant cells like insulin-producing pancreatic β-cells frequently relies on the deployment of signal modulators at different stages of cell differentiation *in vitro*^5^. Moreover, the initiation and progression of tumors is often associated with a change in their competence which, in some instances, is correlated with the erroneous re-activation of embryonic TFs stimulating excessive cell proliferation^6^.

In an effort to analyze systematically the molecular basis of competence, we chose, for the following reasons, to analyze the first inductive signaling events in the embryo of *Xenopus tropicalis*. First, like most multicellular organisms, *X. tropicalis* begins development with a transcriptionally silent genome (**Fig. 1a**). After fertilization, twelve rapid cleavages convert the egg into a mid-blastula embryo with little transcription and little diversification among its 4,096 cells^7^. Both the low levels of transcription and the cellular homogeneity assist in the interpretation of whole-embryo and loss-of-function chromatin profiling. Second, TFs involved in zygotic (or embryonic) genome activation (ZGA) are likely to occupy top-level positions within gene regulatory networks, and their binding should thus correlate strongly with gene target expression^8^. This makes them more amenable to study. Third, based on the nuclear accumulation of signal mediators^9^ (**Fig. 1a**), we know a great deal about where and when inductive signaling occurs during ZGA. And finally, *in vitro* fertilization yields thousands of synchronously developing embryos, which greatly facilitates temporal chromatin profiling.

**Figure 1.**
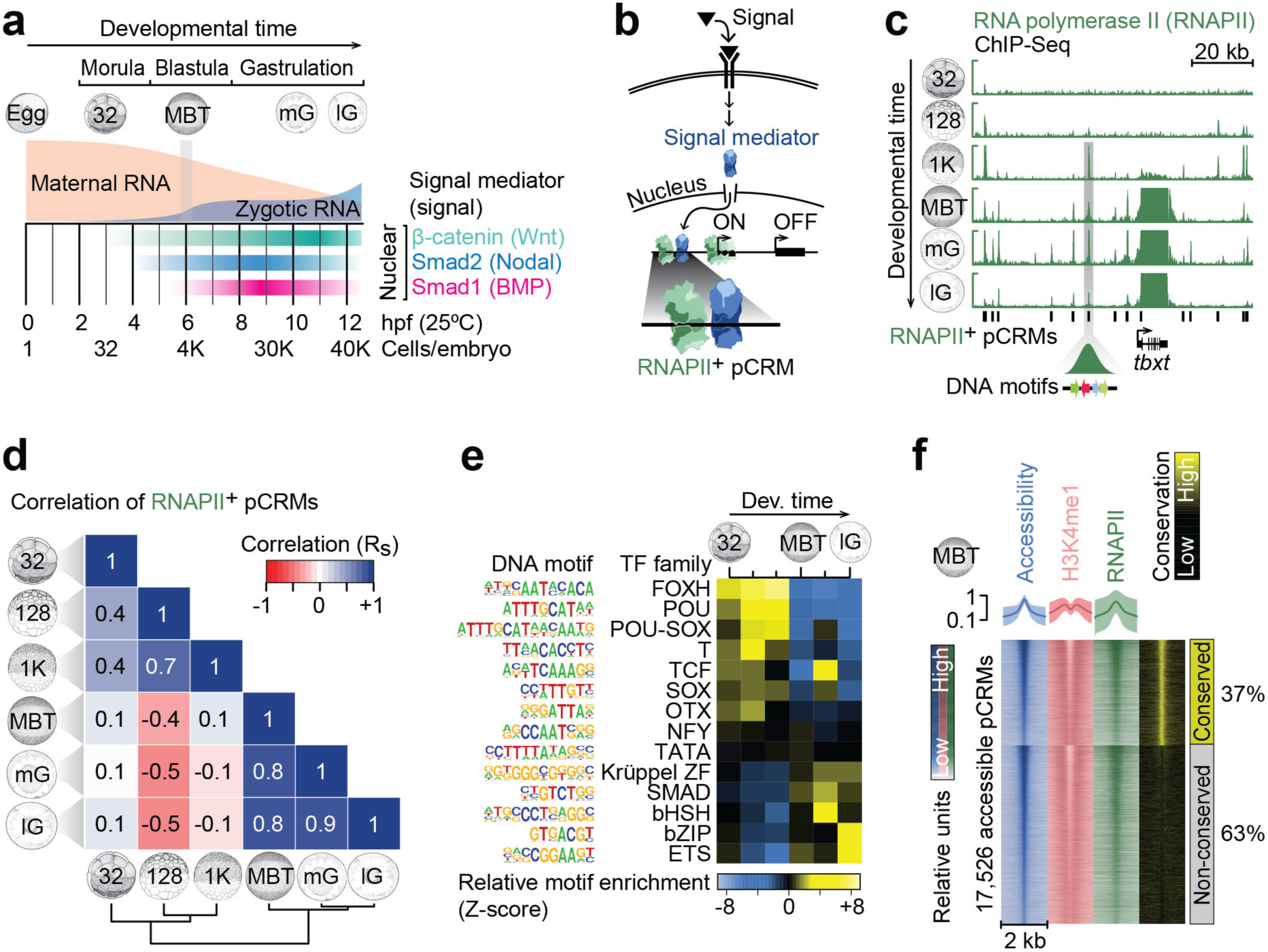
Characterization of pCRMs instructing ZGA. (a) Timeline (hpf, hours post-fertilization, at 25°C) of the maternal-to-zygotic transition and earliest signaling events (nuclear accumulation of Wnt, Nodal and Bmp signal mediators β-catenin, Smad2 and Smad1, respectively) during early *X. tropicalis* development up to the late gastrula embryo (12 hpf) with ∼40,000 (40K) cells. (b) Signal transduction pathway causing signal mediators to enter the nucleus and engage with pCRMs (e.g. marked by RNAPII). (c) Snapshot of RNAPII recruitment to pCRMs of the zygotic gene locus *tbxt* from the 32-cell to the late gastrula stage. The underlying DNA sequence of RNAPII^+^ pCRMs are used to discover enriched DNA motifs *de novo* (illustrated as colored arrows for one RNAPII^+^ pCRM). (d) Spearman correlations (R_s_) of RNAPII binding levels across ∼27,000 pCRMs (Supplementary Table 2) between the indicated developmental stages. (e) Temporal enrichment (Z-score) of consensus DNA motifs known to be recognized by indicated TF families among RNAPII^+^ pCRMs. (f) MBT-staged heat map of DNase-probed chromatin accessibility (n=2), RNAPII binding and H3K4me1 (n=2) marking across ∼17,500 pCRMs (Supplementary Table 3) grouped by sequence conservation levels (phastCons) and sorted by the statistical significance of pCRM accessibility. Abbreviations used for the developmental timeline: 32, 128 and 1K, 32-, 128- and 1,024-cell stage; MBT, mid-blastula transition; eG, mG and lG, early, mid- and late gastrula stage.

Taking advantage of these features of *X. tropicalis*, we have used transcriptional, translational and multi-level chromatin profiling to help identify the earliest regulatory DNA sequences (hereafter called putative *cis*-regulatory modules, pCRMs). From these we infer the critical maternal activators of the zygotic genome as potential competence factors. We next compared the DNA binding of several maternal/zygotic TFs and signal mediators across the mid-blastula transition (MBT) to reveal the effect of TF co-expression on chromatin recruitment *in vivo*. Finally, we have demonstrated how maternal TFs of the pluripotency network structure the chromatin landscape, which in turn determines the signal-mediated regionalization of gene activity and the specification of the three germ layers: ectoderm, mesoderm and endoderm.

## RESULTS

### DNA motifs at pCRMs correlate with the affinities of frequently translated TFs during ZGA

In an effort to understand how early chromatin dynamics influence the recruitment of signal mediators to the genome (**Fig. 1b**), we first identified ∼27,000 pCRMs from the 32-cell to the late gastrula stage by mapping focal RNA polymerase II (RNAPII) depositions on a genome-wide scale by means of ChIP-Seq (**Fig. 1c** and **Supplementary Tables 1** and **2**). RNAPII has no DNA sequence preference and its chromatin engagement is a reliable and objective indicator of pCRM usage^10^. The number of RNAPII-engaged (RNAPII^+^) pCRMs increased from ∼650 at the 32-cell stage to >10,000 at the 1,024-cell and later developmental stages (**Supplementary Fig. 1a**). The largest changes to RNAPII^+^ pCRMs, as calculated by pairwise Spearman correlations, were detected between the 1,024-cell stage and the MBT (**Fig. 1d**), with most pCRMs being engaged only transiently before MBT (e.g., 6,145 from the 128-cell to the 1,024-cell stage) and more persistently after MBT (**Supplementary Fig. 1a**). The analysis of enriched DNA motifs among RNAPII^+^ pCRMs suggests that pre-MBT recruitment is predominantly directed by members of the FOXH, POU, SOX and T domain TF families (**Fig. 1e** and **Supplementary Fig. 1b**).

The discovery of RNAPII^+^ pCRMs is difficult in promoters and gene bodies where extended RNAPII depositions associated with transcript elongation might hamper the correct detection of RNAPII^+^ pCRMs. MBT-staged pCRMs were therefore further characterized for chromatin accessibility and for the enhancer-associated histone mark H3K4me1. In our hands, the high yolk content in early *X. tropicalis* embryos made it impossible to use transposition^11^ to probe chromatin accessibility. Instead, we used an approach involving DNase I mediated digestion followed by deep sequencing (DNase-Seq), in which we selected small accessible fragments of DNA (see **Methods** and **Supplementary Fig. 1c** for exemplar comparison with other chromatin features). We detected ∼17,500 accessible (DNase hypersensitive) pCRMs, ∼85% of which showed both RNAPII and flanking H3K4me1 above background (**Fig. 1f**, **Supplementary Fig. 1d** and **Supplementary Tables 1** and **3**). About 29% and 31% of accessible pCRMs (compared with ∼16% and ∼38% of pCRMs detected by RNAPII peak calling) were found in promoters and gene bodies, respectively (**Supplementary Fig. 1e**).

Enriched DNA motifs among accessible/RNAPII^+^/H3K4me^+^ pCRMs were then correlated with maternally inherited and translated sequence-specific factors identified by egg-staged mass spectrometry^12^ and pre-MBT ribosome footprinting^13^ to identify members of the TF families that may play a role in the ZGA (**Fig. 2a**, **Supplementary Fig. 2a** and **Supplementary Table 4**). It proved that the binding preferences of the most frequently translated maternal TFs and signal mediators matched the most significantly enriched DNA motifs (**Fig. 2b**). These were POU-SOX (Pou5f3-Sox3 heterodimer), Krüppel-like zinc finger (ZF; Sp1 and several Klf), POU (Pou5f3), SOX (Sox3), bZIP (Max), FOXH (Foxh1), ETS (Ets2), NFY (NFYa/b/c), SMAD (Smad1/2), T (mVegT, a maternal VegT isoform), TCF (Tcf/β-catenin), basic helix-span-helix domain (bHSH; Tfap2), and OTX (Otx1). Interestingly, the enrichment level of these motifs was independent of pCRM sequence conservation (based on phastCons scoring) detected among vertebrates (**Figs. 1f** and **2b**).

**Figure 2.**
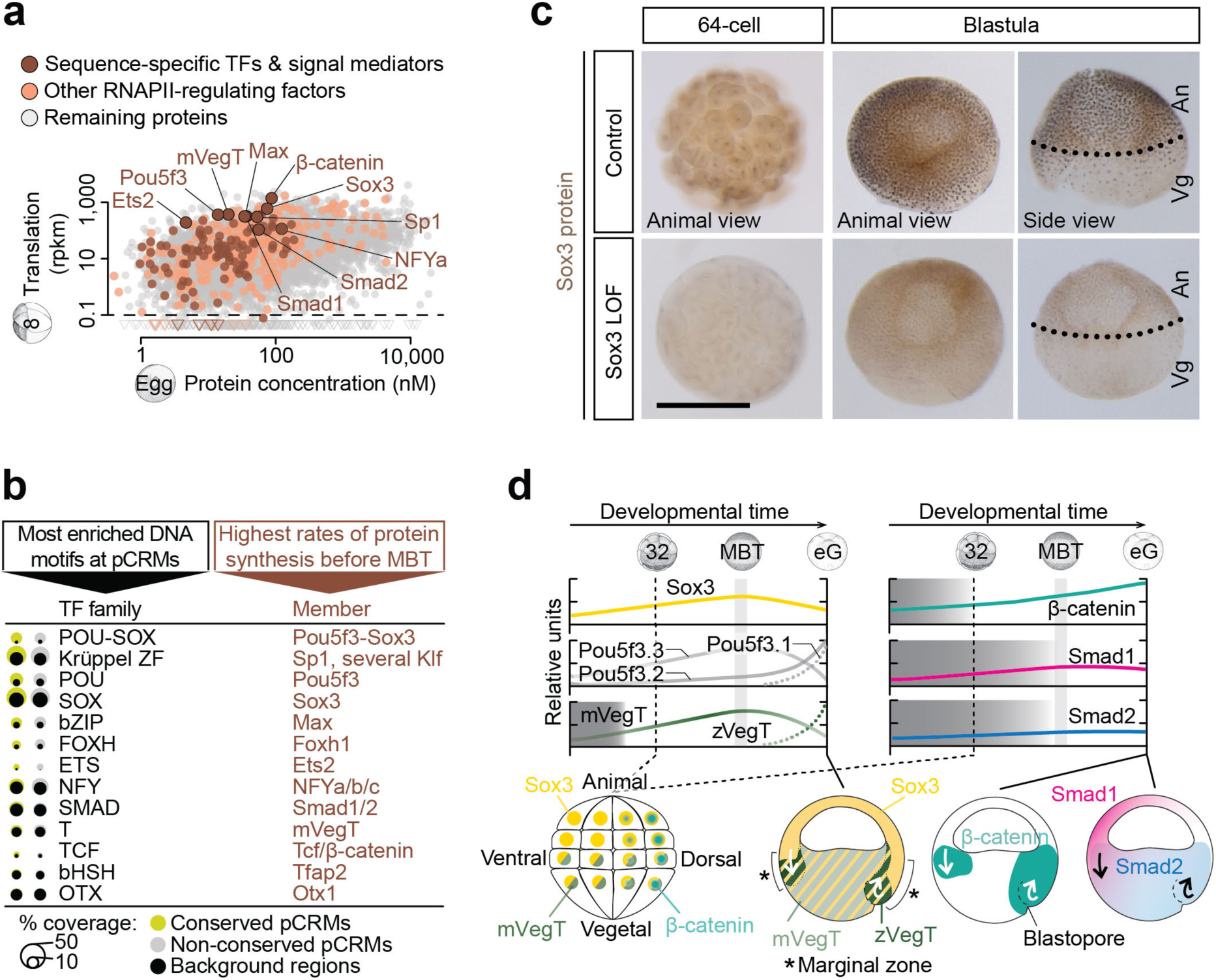
Search for ZGA-critical proteins based on their significantly enriched DNA recognition motifs at accessible and engaged (RNAPII^+^/H3K4me1^+^) pCRMs and their high translation frequency before MBT. (a) Maternal protein concentrations in the egg^12^ versus ribosome footprint (translation) levels at the 8-cell stage^13^. Most frequently translated representatives of various TF families are labeled. (b) Matching canonical pCRM-enriched DNA motifs (sorted by statistical significance) with frequently translated TFs and signal mediators. (c) WMIHC of Sox3 protein in control and Sox3 loss-of-function (LOF) embryos at the 64-cell and blastula stage. Nuclear accumulation of Sox3 protein was detected in both the animal (An) and vegetal (Vg) hemisphere of control embryos. Scale bar, 0.5 mm. (d) Graphical illustration of protein levels (derived from mass spectrometry data^40^) and nuclear localizations (mainly derived from WMIHC, see references below) of selected TFs and signal mediators based on our and previously published results: Sox3 (this study and ref. ^48^), mPouV (Pou5f3.2 and Pou5f3.3) and (zygotic) Pou5f3.1 (deduced from transcript data^14^), mVegT and zVegT^15^, β-catenin^9, 16^, Smad1 (this study and ref. ^9, 17^) and Smad2 (this study and ref. ^9, 17^). Shaded boxes indicate periods of non-nuclear protein localization. Arrows indicate tissue movements of gastrulation. Abbreviations used for the developmental timeline: 8 and 32, 8-cell and 32-cell stage; MBT, mid-blastula transition; and eG, early gastrula stage.

Of the most frequently translated TFs with cognate pCRM-enriched DNA recognition motifs, we selected Pou5f3, Sox3 and mVegT as potential competence factors of canonical Wnt, Nodal and BMP signaling. Sox3 and presumably also Pou5f3 (based on the spatial distribution of its maternal transcripts *Pou5f3.2* and *Pou5f3.3*^14^) are detected ubiquitously (**Fig. 2c**), while mVegT is restricted to the vegetal hemisphere^15^. The zygotic isoform of VegT (zVegT) is expressed within the marginal zone (**Fig. 2d** and **Supplementary Fig. 2e**). With respect to signal mediators, nuclear β-catenin begins to accumulate weakly on the dorsal side of the embryo (first detected at the 32-cell stage^16^) before spreading more prominently around the upper lip of the forming blastopore after the MBT^9^ (**Fig. 2d**). Signal-induced nuclear accumulation of Smad1 and Smad2 is first detected around the MBT, with Smad1 being preferentially ventral and Smad2 being vegetal and within the marginal zone^9, 17^ (**Fig. 2d** and **Supplementary Fig. 2h**).

### pCRM-enriched DNA motifs reflect patterns of chromatin recruitment

Informed by the time of onset of their nuclear accumulation (**Fig. 1a** and **2d**), we generated genome-wide chromatin profiles across the MBT (**Fig. 3a** and **Supplementary Table 1**) to ask whether the recruitment of β-catenin, Smad1 and Smad2 might be influenced by co-expressed maternal TFs like Sox3, mVegT and Foxh1^18^ or vice versa (see **Fig. 2c** and **Supplementary Fig. 2b-n** for ChIP antibody verification). As an outgroup control, we selected the binding profiles of several zygotic T-box TFs including Eomes (Eomesodermin), zVegT, Tbxt (Brachyury)^19^ and Tbx6, which collectively regulate the neuro-mesodermal cell lineage during and beyond gastrulation^20^. The profiled developmental stages and chromatin factors are color-coded as illustrated in **Fig. 3a** and ChIP-Seq peak call coordinates are listed in **Supplementary Table 5**.

**Figure 3.**
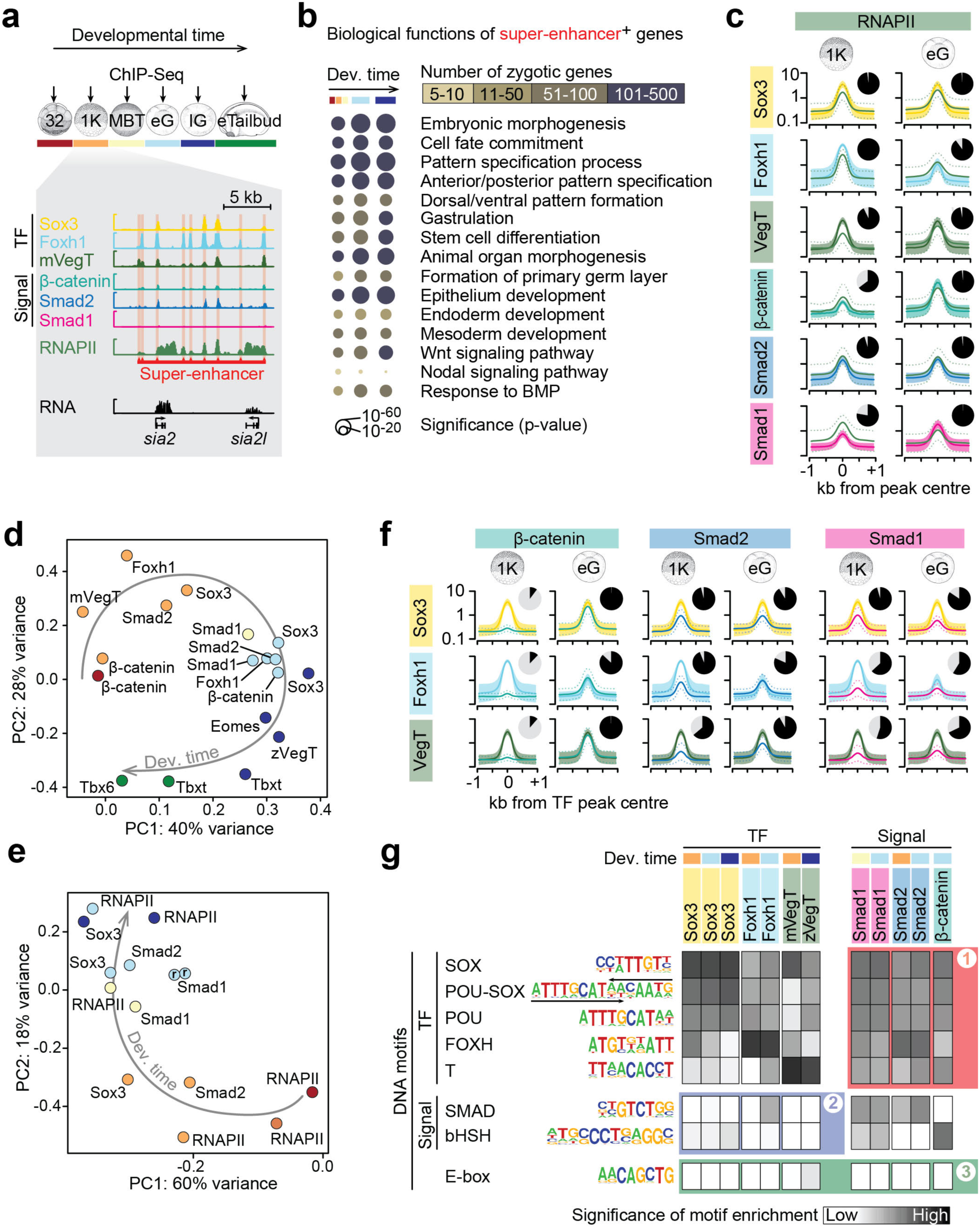
Chromatin engagement of TFs and signal mediators during the maternal-to-zygotic transition. (**a**) Chromatin profiling (ChIP-Seq, n=1-3) of selected TFs and signal mediators from the 32-cell to the early tailbud stage. In all subsequent figure panels, the chromatin factors and developmental stages profiled are consistently color-coded as illustrated here. The excerpt of multiple chromatin tracks shows the binding of maternal TFs (Sox3, Foxh1 and VegT) and signal mediators (β-catenin, Smad2 and Smad1) to the *siamois2* (*sia*2 and *sia2l*) super-enhancer at the 1,024-cell stage (see **Supplementary Fig. 3a** for the temporal progression of chromatin engagement to the *siamois2* and *ventx* super-enhancers). (**b**) Bubble plot shows significantly enriched biological processes associated with zygotic super-enhancer^+^ genes (i.e. genes possessing engaged super-enhancers ≤5 kb from their active TSS at indicated developmental stages). (**c**) Meta-plots (mean [solid line] ± SD [dotted line or polygon]) summarize the level of RNAPII engagement across 2,000 pCRMs most frequently occupied by the indicated TFs or signal mediators at the 1,024-cell and early gastrula stage, respectively. The pie chart next to each meta-plot shows the percentage of these TF^+^ or signal mediator^+^ pCRMs bound by RNAPII (ChIP ≥2x input tag density): 98% (1,024-cell stage) and 99% (early gastrula stage) of Sox3^+^ pCRMs, 100% and 90% of Foxh1^+^ pCRMs, 93% and 97% of VegT^+^ pCRMs, 65% and 98% of β-catenin^+^ pCRMs, 96% and 98% of Smad2^+^ pCRMs, and 78% and 99% of Smad1^+^ pCRMs. (**d**) Biplot of principal component (PC) 1 (accounting for 40% variance) and 2 (28% variance) shows the similarity (or dissimilarity) of TF (Sox3, mVegT, Foxh1^18, 25^, Eomes^19^, zVegT^19^, Tbxt^19^ and Tbx6) and signal mediator (β-catenin, Smad1, Smad2) binding levels across ∼12,500 highly engaged pCRMs (compiled from the 2,000 pCRMs with the highest DNA occupancy levels detected per protein and developmental stage) over several developmental stages. Note that developmental time (arrow) separates these profiles best. (**e**) Biplot of PC1 (accounting for 60% variance) and PC2 (18% variance) for the temporal progression of RNAPII and TF binding levels across the same set of pCRMs as in (**d**). Abbreviation: r, biological replicates. (**f**) Meta-plots (mean [solid line] ± SD [dotted line or polygon]) summarizes the level of signal mediator engagement across 2,000 pCRMs most frequently occupied by the indicated TF at the 1,024-cell and early gastrula stage, respectively. The pie chart next to each meta-plot shows the percentage of these TF^+^ pCRMs bound by signal mediators (ChIP ≥2x input tag density): β-catenin at Sox3^+^ pCRMs (10% at the 1,024-cell stage and 100% at the early gastrula stage), Smad2 at Sox3^+^ pCRMs (96% and 91%), Smad1 at Sox3^+^ pCRMs (96% and 85%), β-catenin at Foxh1^+^ pCRMs (12% and 87%), Smad2 at Foxh1^+^ pCRMs (94% and 82%), Smad1 at Foxh1^+^ pCRMs (63% and 59%), β-catenin at VegT^+^ pCRMs (11% and 100%), Smad2 at VegT^+^ pCRMs (64% and 92%) and Smad1 at VegT^+^ pCRMs (55% and 68%). (**g**) Heat map shows the statistical significance of finding TF- and signal-specific DNA consensus motifs (y-axis) across 2,000 pCRMs most frequently occupied by the indicated TFs or signal mediators (x-axis). Abbreviations used for the developmental timeline: 32 and 1K, 32-cell and 1,024-cell stage; MBT, mid-blastula transition; eG and lG, early and late gastrula stage; and eTailbud, early tailbud.

Maternal and zygotic TFs and signal mediators shared many chromatin characteristics: (1) DNA occupancy levels followed a log-normal distribution with <1,000 RNAPII-transcribed genes receiving high and super-enhancer-like^21^ input (i.e. clusters of occupied pCRMs separated by ≤25 kb). Similar distributions were also observed for chromatin accessibility and RNAPII engagement (see examples in **Fig. 3a** and **Supplementary Fig. 3a,b** and systematic analysis in **Supplementary Fig. 4a**). (2) From the 1,024-cell to the late gastrula stage, TF-bound super-enhancers were linked through the gene ontologies of nearby zygotic genes (≤5 kb) to early embryonic processes including germ layer and body axis formation (**Fig. 3b**). (3) The binding sites of TFs and signal mediators were also frequently defined by RNAPII deposition (**Fig. 3c**). (4) Among all zygotic genes, promoter-proximal regions were more consistently bound than other pCRMs (**Supplementary Fig. 4b**).

Differences in DNA occupancy levels at different developmental stages, compared by pairwise Spearman correlations and principal component (PC) analysis, suggest that the recruitment to chromatin of signal mediators and TFs is driven both by their individual properties and by the developmental stage (**Fig. 3d** and **Supplementary Fig. 5a**). For example, chromatin recruitment of mVegT at stages of pluripotency resembled that of other sequence-specific factors at the same developmental stage but differed from the binding of its zygotic isoform and of related T-box TFs at later developmental stages (highlighted in **Supplementary Fig. 5a**). The importance of developmental stage in driving patterns of chromatin recruitment was further revealed by the changing DNA binding patterns of sequence-nonspecific RNAPII to pCRMs (**Fig. 3e**).

The identification of enriched DNA recognition motifs at occupied pCRMs confirmed known TF/signal mediator properties such as the sequence-specificity of their DNA binding domains, oligomerization tendencies and protein-protein interactions (**Supplementary Fig. 5b**). For example, in contrast to other T-box TFs, Tbxt recognizes palindromic T motifs due to its propensity to form homodimers^22^; and Smad2 chromatin recruitment is frequently associated with the FOXH motif because Smad2 interacts with Foxh1^23^ (**Supplementary Fig. 5b**). Other DNA motifs such as the POU or POU-SOX motifs were consistently co-enriched from the 1,024-cell to the late gastrula stage in most binding profiles. This is indicative of the pluripotent state, a developmental context associated with co-expression of Pou5f (Oct4) and SoxB1 (e.g. Sox2 or Sox3) proteins as previously observed *in vitro*^24^ (**Fig. 3g** and **Supplementary Fig. 5b**).

The hierarchical clustering of DNA occupancy levels for the selected factors at different developmental stages revealed specific DNA motif combinations. For example, pCRMs showing ‘unique’ binding of mVegT (that is, binding that is not shared with the other profiled factors) show a high frequency of T, OTX and SOX motifs (**Supplementary Fig. 6**).

With respect to the chromatin recruitment of TFs versus signal mediators, we note that, first, Smads and/or β-catenin were frequently detected at Sox3, Foxh1 or VegT binding sites (top 2,000 peaks shown in **Fig. 3f**) and, second, Smad-and/or β-catenin-bound pCRMs (top 2,000 peaks) were significantly enriched for SOX, POU-SOX, POU, FOXH and T motifs (red field #1 in **Fig. 3g**) suggesting that corresponding TFs affect the recruitment of these signal mediators. The reverse was much less the case as shown by the low significance of SMAD and bHSH motif enrichments at TF-bound pCRMs (with the exception of Smad2-interactor Foxh1^23^) (blue field #2 in **Fig. 3g**).

### TF co-expression drives the dynamics of chromatin recruitment

To explore the suggested importance of TF co-expression on pCRM engagement, we ectopically expressed an HA-tagged version of the muscle determinant MyoD (MyoD-HA) in animal cap cells and profiled pCRMs for MyoD-HA as well as for endogenous Sox3 and RNAPII at the early gastrula stage (**Fig. 4a** and **Supplementary Tables 1** and **6**). MyoD was chosen because its canonical E-box recognition motif is normally not significantly enriched before or during gastrulation (green field #3 in **Fig. 3g** and **Supplementary Fig. 5b**) so its effect on chromatin engagement ought to be clearly discernible, while Sox3 and RNAPII were selected because they are ubiquitously expressed and represent sequence-specific and nonspecific DNA binding factors, respectively. The ectopic expression elevated the Spearman correlations of Sox3 and RNAPII with MyoD-HA (**Supplementary Fig. 7a**) and shifted the first and second PC of Sox3 and RNAPII toward MyoD-HA (**Fig. 4b**) suggesting that MyoD-HA altered the binding of both Sox3 and RNAPII. Differential binding analysis on a genome-wide (heat map in **Fig. 4c**) and locus-specific (pileup track in **Fig. 4d**) scale confirms that MyoD-HA co-recruits endogenous Sox3 and RNAPII to its gene targets like *actc1* and *myl1*. The E-box motif of MyoD-HA emerged as a significantly enriched motif of Sox3 and RNAPII binding, while MyoD-HA binding itself seemed to be influenced by endogenous TFs as judged by the developmental stage-characteristic enrichment of SOX, POU-SOX and FOX motifs at MyoD^+^ pCRMs (**Fig. 4e**). However, this opportunistic recruitment to non-canonical binding sites, such as MyoD-HA to functional Sox3 gene targets (e.g. *otx2* and *sox2* activated by Sox3-HA), did not affect transcription in animal caps (**Fig. 4f-h**).

**Figure 4.**
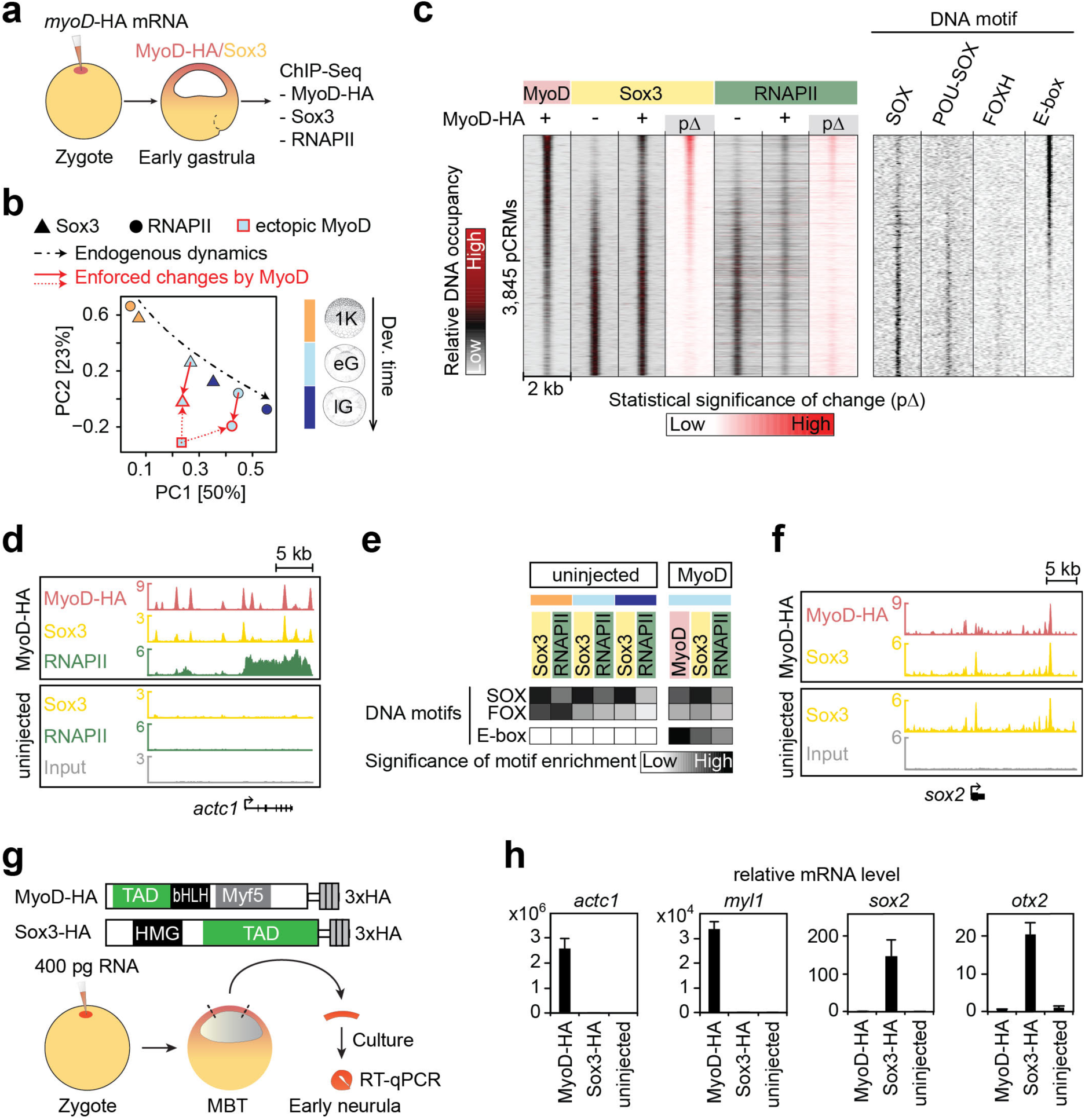
Ectopic expression of the muscle determinant MyoD reveals the effect of co-expressed TFs on chromatin engagement and gene expression. (a) Experimental design: Ectopic expression of the MyoD-HA mRNA construct injected into the animal hemisphere followed by the genome-wide chromatin profiling (ChIP-Seq; n=2) of MyoD-HA, Sox3 and RNAPII in early gastrula embryos. (b) Biplot of principal component (PC) 1 (accounting for 50% variance) and 2 (23% variance) shows the relationship of Sox3 (triangle), RNAPII (circle) and MyoD-HA (square) binding levels across MyoD^+^ and/or Sox3^+^ pCRMs. Arrows show the normal temporal dynamics of Sox3 and RNAPII binding (black dash-dotted line) and the MyoD-enforced (red dotted line) changes to them (red solid line) at early gastrula stage. Fill color of symbols represents the developmental stage as indicated, while line colors refer to whether MyoD-HA was expressed (red) or not (black). Abbreviations used for the developmental timeline: 1K, 1,024-cell stage; eG and lG, early and late gastrula stage. (c) Heat map of the DNA occupancies of 3,845 pCRMs and enriched DNA motifs sorted by the significance of MyoD-HA-enforced changes (pΔ) to Sox3 binding levels. (d,f) Snapshot of chromatin co-recruitment to the super-enhancers of canonical MyoD and Sox3 target genes *actc1* and *sox2*, respectively. (e) Heat map shows the significance of DNA motif enrichments (y-axis) at 10,000 pCRMs most frequently occupied by the indicated proteins in uninjected and MyoD-HA injected embryos (x-axis). (g) Experimental design: Animal cap assay to quantify MyoD- or Sox3-enforced transcription. (e) RT-qPCR results from the animal cap assay. Error bars, mean+SD (n=2).

The influence of TF co-expression on DNA occupancy was further substantiated by profiling chromatin for Sox3 in different anterior-posterior regions of the central nervous system (CNS) (**Fig. 5a** and **Supplementary Tables 1** and **7**). The analysis of enriched DNA motifs suggested that Sox3 binding was affected by differentially expressed homeodomain TFs such as orthodenticle homeobox (OTX) in the brain (head) and caudal homeobox (CDX) in the spinal cord (trunk, bud) (**Fig. 5c** and **Supplementary Fig. 7c,d**; see **Fig. 5d** for graphical illustration of TF co-expression at the posterior end of the embryo). This was particularly apparent in Sox3 binding to the colinear *HoxD* cluster defining anterior-posterior cell identity (**Fig. 5b**). Similar influences on chromatin engagement were observed within anterior and posterior mesoderm marked by Eomes in gastrula embryos and Tbxt/Tbx6 in gastrula and early tailbud embryos, respectively (**Fig. 5c,d** and **Supplementary Fig. 7b,c**). On a temporal scale, the influence of FOXH and POU motif-recognizing TFs in recruiting Sox3 and T-box TF was more pronounced early than late in development (**Fig. 5c** and **Supplementary Figs. 5b** and **7c**).

**Figure 5.**
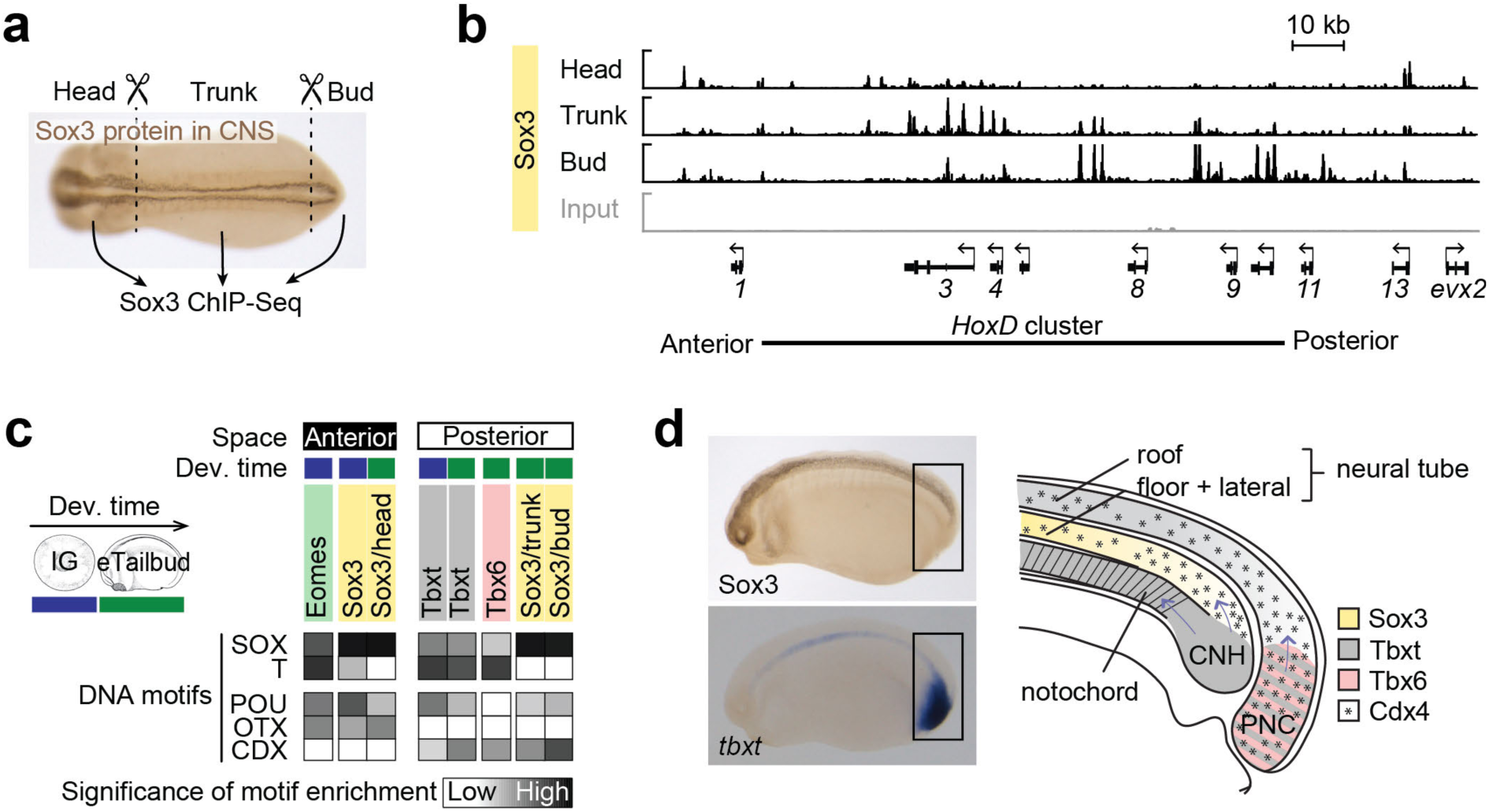
Profiling chromatin for Sox3 in different anterior-posterior compartments of the central nervous system (CNS). (a) Experimental design: Genome-wide Sox3 profiling of head, trunk and bud dissected from early tailbud embryos after tissue fixation. (b) Snapshot of Sox3 binding to *HoxD* cluster in the head, trunk and bud. Note the differential binding of Sox3 with ‘anterior’, ‘middle’ and ‘posterior’ Hox genes being preferentially occupied in head, trunk and bud, respectively. (c) Heat map shows the significance of DNA motif enrichments (y-axis) across 10,000 pCRMs most frequently occupied in each chromatin profile (x-axis). Profiles are grouped depending on whether the cells expressing the TF of interest preferentially contribute to the anterior or posterior compartment. Abbreviations used for the developmental timeline: lG, late gastrula; and eTailbud, early tailbud. (d) Anatomical map of TF expression at the posterior end of an early tailbud embryo to explain the co-enrichment of ‘posterior’ TF-specific DNA recognition motifs in (c). Sox3-expressing cells of the posterior neural tube originate from the chordoneural hinge (CNH) and posterior wall of the neurenteric canal (PNC) expressing Brachyury and Brachyury/Tbx6, respectively^20^. Both neural tube and PNC are also exposed to the expression of Cdx such as Cdx4^49^.

### Signal-induced regionalization of ZGA depends on maternal TFs

To ask whether signal-mediated ZGA requires maternal TFs, as suggested by the observed chromatin dynamics (**Fig. 3**), we next compared the effects of loss of maternal Sox3/PouV (Pou5f3.2 and Pou5f3.3; mPouV) or VegT (mVegT), and canonical Wnt, Nodal or BMP signal transduction, on zygotic transcription from MBT to late blastula and early gastrula stages (**Fig. 6c** and **Supplementary Table 1**). The high translation frequencies (**Fig. 2a** and **Supplementary Fig. 2a**) coupled with fairly steady or even transient (mVegT and Pou5f3.3) protein levels around MBT (**Fig. 2d**), suggest that maternal TFs and signal mediators have short half-lives. Consistent with this idea, the injection of antisense morpholino oligonucleotides (MOs) blocking the translation of maternal transcripts such as *Sox3* or *mVegT* was effective in reducing protein levels (**Fig. 2c** and **Supplementary Fig. 2c,e,f**). We used previously-validated MOs to knock down mPouV^25^ and β-catenin^26^. Although MOs can cause off-target mis-splicing and the induction of an immune response, the effects on splicing do not affect mature maternal transcripts and the immune response is only detectable beyond gastrula stages^27^. The nuclear accumulation of Smad1 and Smad2 in response to BMP and Nodal signaling was inhibited with the small molecules LDN193189^28, 29^ and SB431542^30, 31^, respectively. The morphological defects of these loss-of-function (LOF) treatments ranged from undetectable (Sox3), to weak (mVegT, BMP), to moderate (mPouV), to severe (mPouV/Sox3, Nodal, β-catenin) (**Fig. 6a** and **Supplementary Fig. 8a,c**). Moderate and severe defects affected gastrulation, while weak ones were only obvious later. All phenotypes were either in line with previous publications (β-catenin^26^, Nodal^32^ and BMP^33^) or could be rescued at the morphological (mPouV/Sox3; **Fig. 6b**) or transcriptional level (mVegT; **Supplementary Fig. 8b**) by co-injecting cognate mRNA. Interestingly, the role of maternal Sox3 could only be detected by knocking it down together with mPouV. In contrast to single Sox3 or mPouV LOF, double LOF embryos failed to close the blastopore (**Fig. 6a** and **Supplementary Movie 1**).

**Figure 6.**
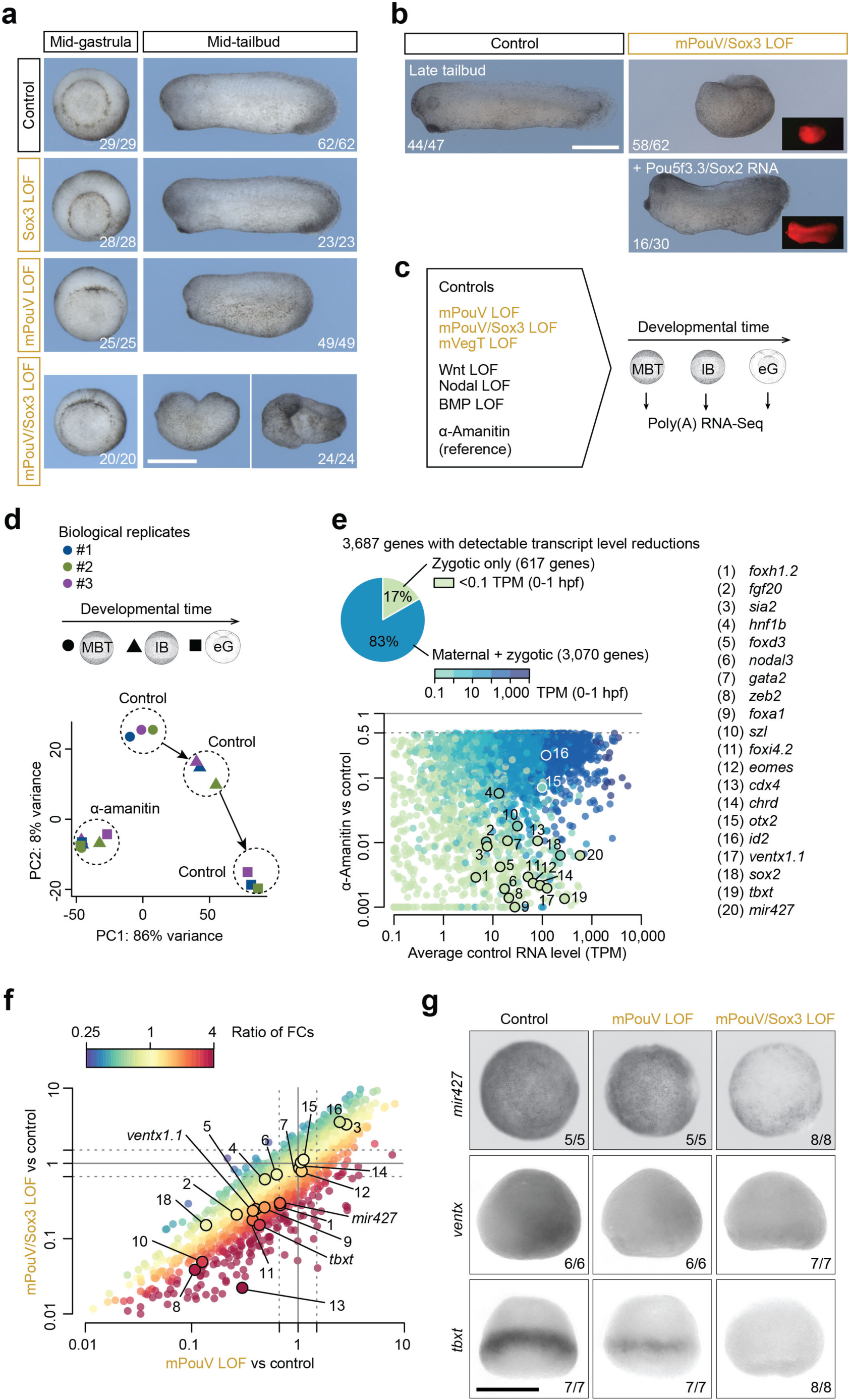
Synergistic relationship between maternal Pou5f3 (mPouV) and Sox3. (a) Morphological phenotypes caused by single and combined LOFs of Sox3 and mPouV when control embryos reached mid-gastrula and mid-tailbud stage. See also Supplementary Movie 1. (b) Phenotypical rescue of mPouV/Sox3 LOF embryos by the co-injection of both *X. laevis Pou5f3.3* and *Sox2* mRNA alongside *mCherry* mRNA as a tracer. (c) Experimental design: Profiling the poly(A) RNA transcriptome (n=3) over three consecutive developmental stages under indicated conditions. Abbreviations used for the developmental timeline: MBT, midblastula transition; lB, late blastula; and eG, early gastrula. (d) Biplot of PC1 (accounting for 86% variance) and PC2 (8% variance) shows the relationship of developmental stage-specific poly(A) RNA transcriptomes of control and α-amanitin-injected embryos in biological triplicates (#1-3). (e) Detection of 3,687 zygotic genes with reduced transcript levels (≥50%, FDR ≤10%) in α-amanitin-injected embryos. These genes were used as reference for all other LOFs. Dots in scatterplot are colored according to the maternal contribution^50^ to the transcript level of each of these zygotic genes. (f) Scatterplot of transcript fold changes (FCs) caused by mPouV and mPouV/Sox3 LOFs with dots colored according to the ratio of FCs. Numbered dots refer to some developmentally relevant genes listed in (e). (g) Early gastrula-staged WMISH: *mir427*, animal view; ventx, lateral view, ventral side facing right; and *tbxt*, dorsal view. Numbers in the right bottom corner of each image refer to the count of embryos detected with the displayed WMISH staining among all embryos analyzed per condition and *in situ* probe. Scale bars, 0.5 mm.

Transcriptome analysis of TF and signal LOF embryos was confined to the 3,687 zygotic genes for which ≥two-fold reductions in exonic and/or intronic transcript level (see **Online Methods**) could be detected following injection of the RNAPII inhibitor α-amanitin (**Supplementary Table 9**). About 83% of these genes had a maternal contribution of ≥1 per 10 million transcripts as detected within first hour post-fertilization when the genome is quiescent (**Fig. 6e**). α-Amanitin prevented any transcriptional changes and thus blocked the gastrulation movements that are normally initiated by ZGA (**Fig. 6d** and **Supplementary Fig. 8d**). The LOF-mediated reduction of ZGA ranged from ∼2% for BMP to ∼25% for mPouV/Sox3 (**Supplementary Fig. 11a**). We note that the additional loss of Sox3 in mPouV LOF embryos further reduced ZGA of developmentally relevant genes, including those that are expressed ubiquitously, like *mir427*, and those that are activated in a subset of mPouV/Sox3^+^ cells, like *ventx* and *tbxt* (**Fig. 6f,g**).

Spatial analysis of reduced gene activation confirmed that many of the 918 mPouV/Sox3-dependent genes showed enriched expression along the animal-vegetal or dorso-ventral axes (**Fig. 7a,e**). More specifically, comparison with signal LOFs revealed that 268 of the 708 genes induced by Wnt, Nodal or BMP also depended on mPouV/Sox3 (**Fig. 7b** and **Supplementary Fig. 9** and **10b**). Similarly, 175 of the 239 mVegT-dependent genes, including such as *foxa1*, *nodal5/6* and *sox17a/b*, were activated in response to signaling (**Supplementary Figs. 9** and **10a**). Loss of mVegT and mPouV/Sox3 jointly affected 71 genes, 57 of which also depended on Wnt, Nodal or BMP signaling (**Supplementary Fig. 10b**).

**Figure 7.**
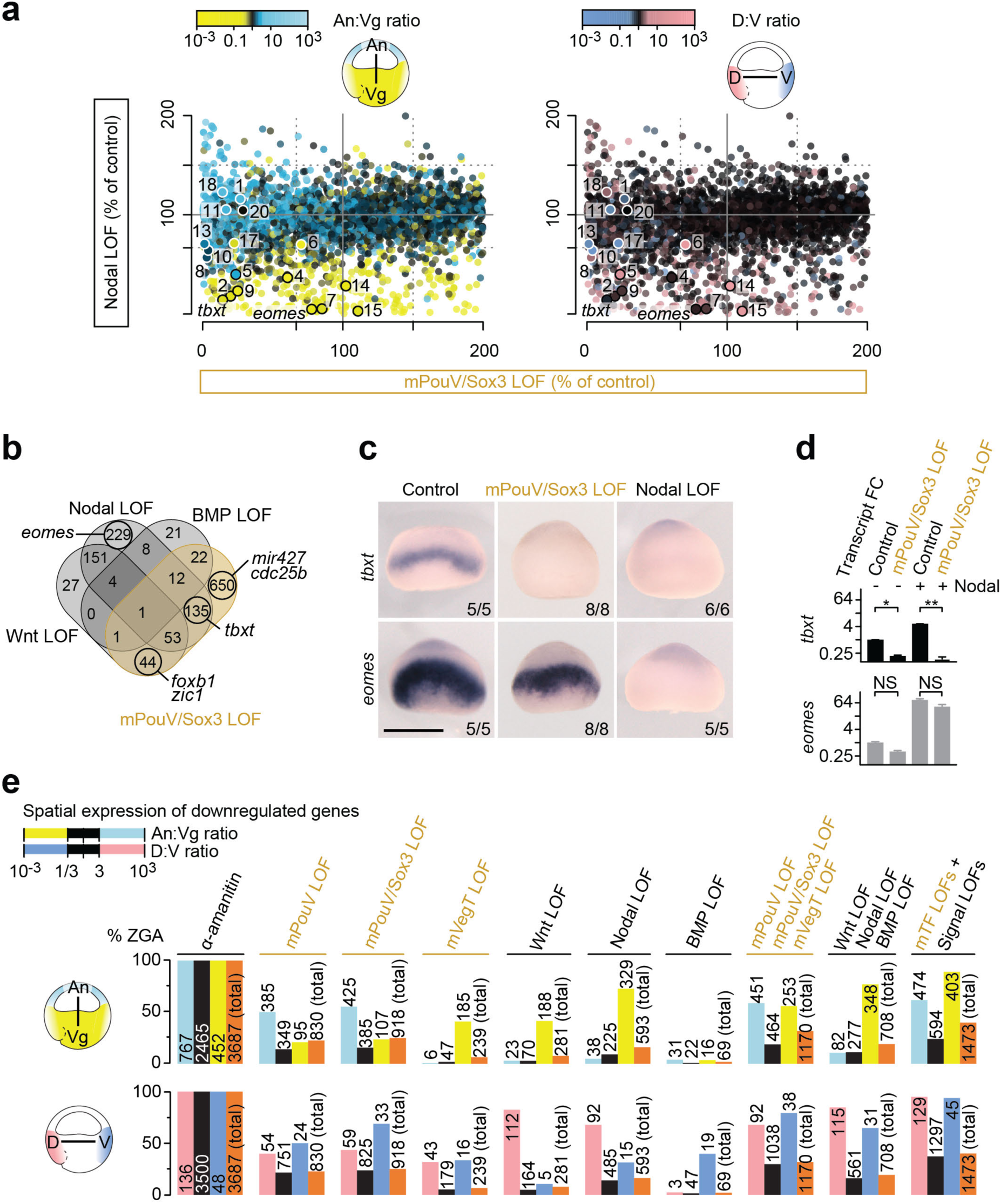
Signal-induced regionalization of ZGA depends on maternal TFs. (a) Transcriptional comparison of zygotic genes between the LOFs of maternal PouV/Sox3 and Nodal signaling. Dots are colored according to the normal ratio of transcript levels (regional expression) across the animal-vegetal (An:Vg) or dorso-ventral (D:V) axis^51^. Numbered dots refer to genes listed in Fig. 6e. (b) Venn diagram of genes downregulated by the mPouV/Sox3 LOF (orange) or the LOF of single signal transduction pathways (black). (c) Early gastrula-staged WMISH for *tbxt* and *eomes* under indicated LOFs. Scale bar, 0.5 mm. (d) Quantification of *tbxt* and *eomes* transcript levels in control and mPouV/Sox3 LOF animal caps with or without stimulated Nodal signaling. Error bars, mean+SD (n=2). Two-tailed Student’s t-test: *, p=0.014; **, p=0.005; and NS, not significant (p≥0.02). (e) Bar graphs show the percentage (and number) of downregulated zygotic genes (% ZGA under indicated LOFs) grouped by the normal ratio of transcript levels (regional expression) across the animal-vegetal (An:Vg) and dorso-ventral (D:V) axis.

Remarkably, the requirement for particular maternal TFs varied even among related genes with similar expression patterns that are activated by the same signals. For instance, *tbxt* and *eomes* are both activated by Nodal signaling in the marginal zone between the animal and vegetal hemispheres, but only *tbxt* requires mPouV/Sox3 for its signal responsiveness (**Fig. 7a-c**). Significantly, DNA occupancy levels did not explain the difference in gene regulation between *tbxt* and *eomes*, because both gene loci showed high levels of Sox3 binding (see Sox3 track in **Fig. 10a,b**). Similar dependencies were found for Wnt-responsive genes like *foxb1* and *zic1* (**Supplementary Fig. 10c**). We confirmed these mPouV/Sox3-dependent signal inductions by treating control and mPouV/Sox3-depleted animal cap tissue with Nodal (Activin) and Wnt (CHR99021) agonists (**Fig. 7d** and **Supplementary Fig. 10d**). Interestingly, mPouV/Sox3 also facilitated basal low-level expression of (especially) *tbxt* and *foxb1* without Nodal or Wnt stimulation.

Together, our selected maternal TFs and signals activated 1,473 of the 3,687 (∼40%) zygotic genes. These included substantial percentages of genes with preferential expression in the animal (∼62%), the vegetal (∼89%), the dorsal (∼95%) and the ventral (∼94%) parts of the embryo, while many ubiquitously expressed genes remained unaffected (**Fig. 7e**). mPouV/Sox3 and mVegT tend to affect biological functions for which signal-induced regionalization of ZGA is essential, such as the formation of the main body axes and the segregation of the three germ layers (**Supplementary Fig. 11b**). We also note that LOF of mPouV/Sox3 caused significant increases in transcript levels of genes encoding gamete-specific biological processes, suggesting that reprogramming towards embryonic pluripotency was compromised (**Supplementary Fig. 11a,b**).

### Pioneering activity of pluripotency TFs predefines signal-mediated gene induction

To discover how maternal TFs allow cell type-specific genes to be signal-induced, we compared various chromatin features from genome-wide accessibility (DNase-Seq; **Fig. 8a** and **Supplementary Fig. 12a**) and DNA occupancy (ChIP-Seq for H3K4me1, RNAPII, Smad2 and β-catenin; **Fig. 8a**) to the high-resolution conformation contacts (next-generation capture-C; **Fig. 9a**, **Supplementary Fig. 12b** and **Supplementary Table 10**) of 30 selected promoters (**Fig. 9b**) between control and mPouV/Sox3 LOF embryos at MBT (**Supplementary Table 1**). We selected mPouV/Sox3 rather than mVegT because of their stronger effect on the ZGA (**Supplementary Fig. 11a**). Genome-wide analysis showed that mPouV/Sox3 LOF strongly reduced chromatin accessibility as measured by the significant (FDR ≤10%) loss of DNase cleavages in 6,738 of the 16,637 (∼41%) pCRMs (**Fig. 8b,c** and **Supplementary Table 11**). Sorting of these pCRMs according to the significance of lost accessibility (**Fig. 8d**) suggests that chromatin opening depends on the pioneering activity of mPouV and Sox3 to recognize their canonical motifs in compacted chromatin (**Fig. 8e**). By comparison, unaffected pCRMs contained canonical POU/SOX motifs less frequently, and were strongly enriched for promoter-centric motifs of the Krüppel ZF, bZIP and NFY protein families (**Fig. 8e,f**). At the extreme end of affected loci (e.g. *tbxt*, *foxb1*, *cdc25b* and *zic1*) entire or large proportions of super-enhancers became inaccessible upon mPouV/Sox3 LOF (**Figs. 9d** and **10a** and **Supplementary Figs. 13** and **14**).

**Figure 8.**
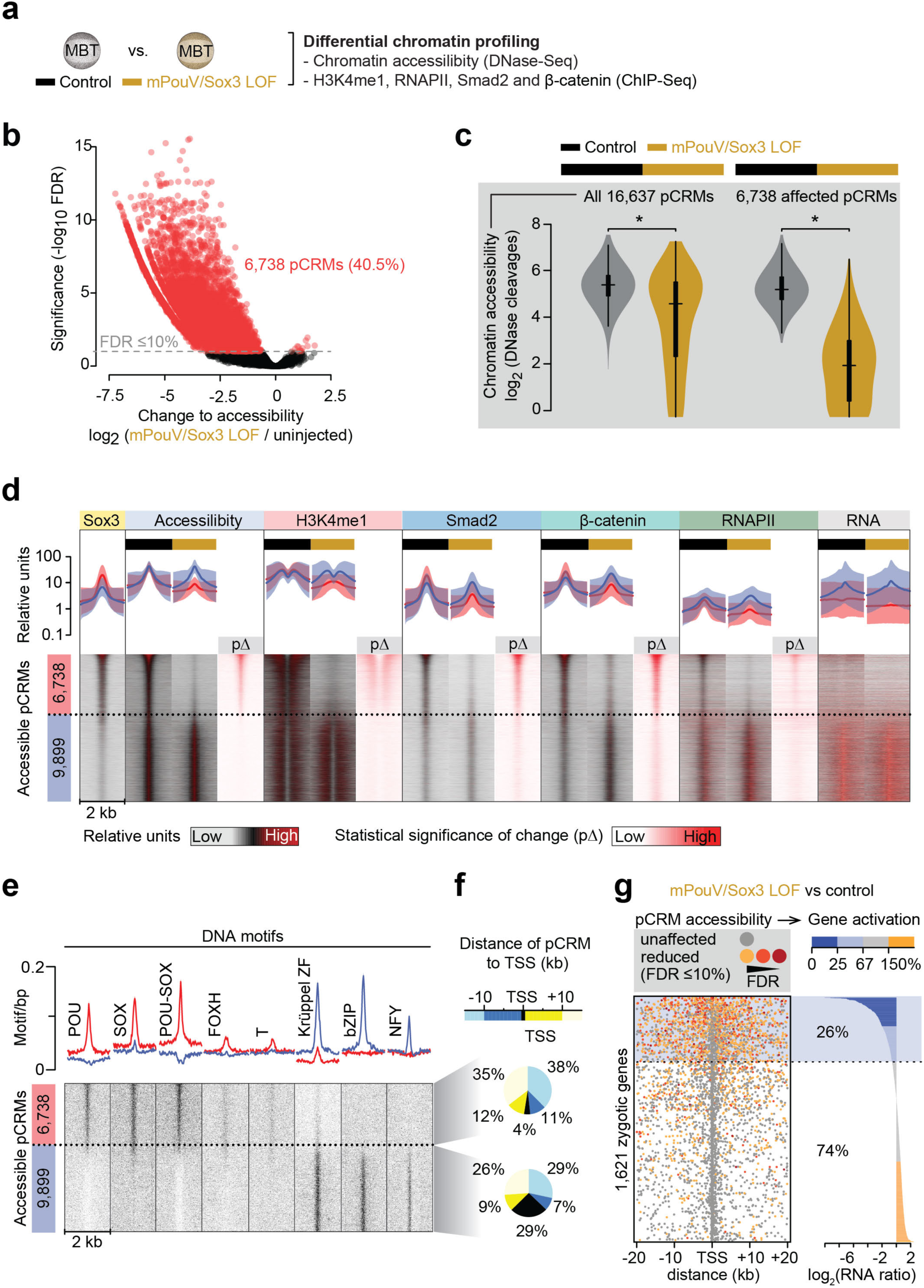
Maternal pluripotency TFs mPouV and Sox3 remodel 40% of the accessible chromatin landscape to contribute to one quarter of ZGA. (a) Used approach to reveal the effect of mPouV/Sox3 on chromatin accessibility (DNase-Seq; n=2) and chromatin composition (ChIP-Seq; n=2) at MBT. (b) Double-logarithmic volcano plot shows massive chromatin accessibility (DNase cleavage) reductions caused by mPouV/Sox3 LOF. pCRMs (dots) with significant accessibility changes (n=2, FDR ≤10%) are marked in red. (c) Violin plots show the comparison of chromatin accessibility (DNase cleavages) between uninjected and mPouV/Sox3 LOF embryos at all and affected (FDR ≤10%) pCRMs. Wilcoxon test: *, p <2.2×10^−16^. (d) Normalized meta-plots (top row, mean±SD) and heat maps (bottom row) show the level of chromatin accessibility, chromatin engagement (Sox3, H3K4me1, β-catenin, Smad2 and RNAPII) and RNA (n=3) across accessible pCRMs in uninjected and mPouV/Sox3 LOF embryos. The β-catenin binding profile was generated at late blastula stage rather than the 1,024-cell stage (Sox3) or MBT (all others). RNA was profiled at and beyond MBT as shown in Fig. 6c. In the heat map, the pCRMs are sorted and grouped by significantly reduced DNase cleavages under mPouV/Sox3 LOF: red group, affected (FDR ≤10%) and blue group, unaffected (FDR >10%). These groups are represented in the meta-plots. Each heat map under mPouV/Sox3 LOF are followed by a heat map showing the statistical significance of changes (pΔ) caused by mPouV/Sox3 LOF. (e) Heat map shows the occurrence of DNA motifs at accessible pCRMs sorted and grouped as in (d). (f) Pie charts summarize the distribution of distances (kb) to nearest zygotic TSSs of affected (top pie chart) and unaffected (bottom pie chart) pCRMs. (g) Panel compares the effect of mPouV/Sox3 LOF on chromatin accessibility and RNAPII-mediated gene expression. Plot to the left shows the localization of accessible pCRMs (affected, dot colored in orange to red with FDR decreasing from 10%; and unaffected, grey dot) relative to the zygotic TSSs that are active by the MBT^52^ and produce enough RNA transcripts to show significant ≥two-fold reductions upon α-amanitin injection (Fig. 6e). Gene loci are sorted by mPouV/Sox3 LOF-induced transcript fold changes as shown in the log-scaled bar graph to the right.

**Figure 9.**
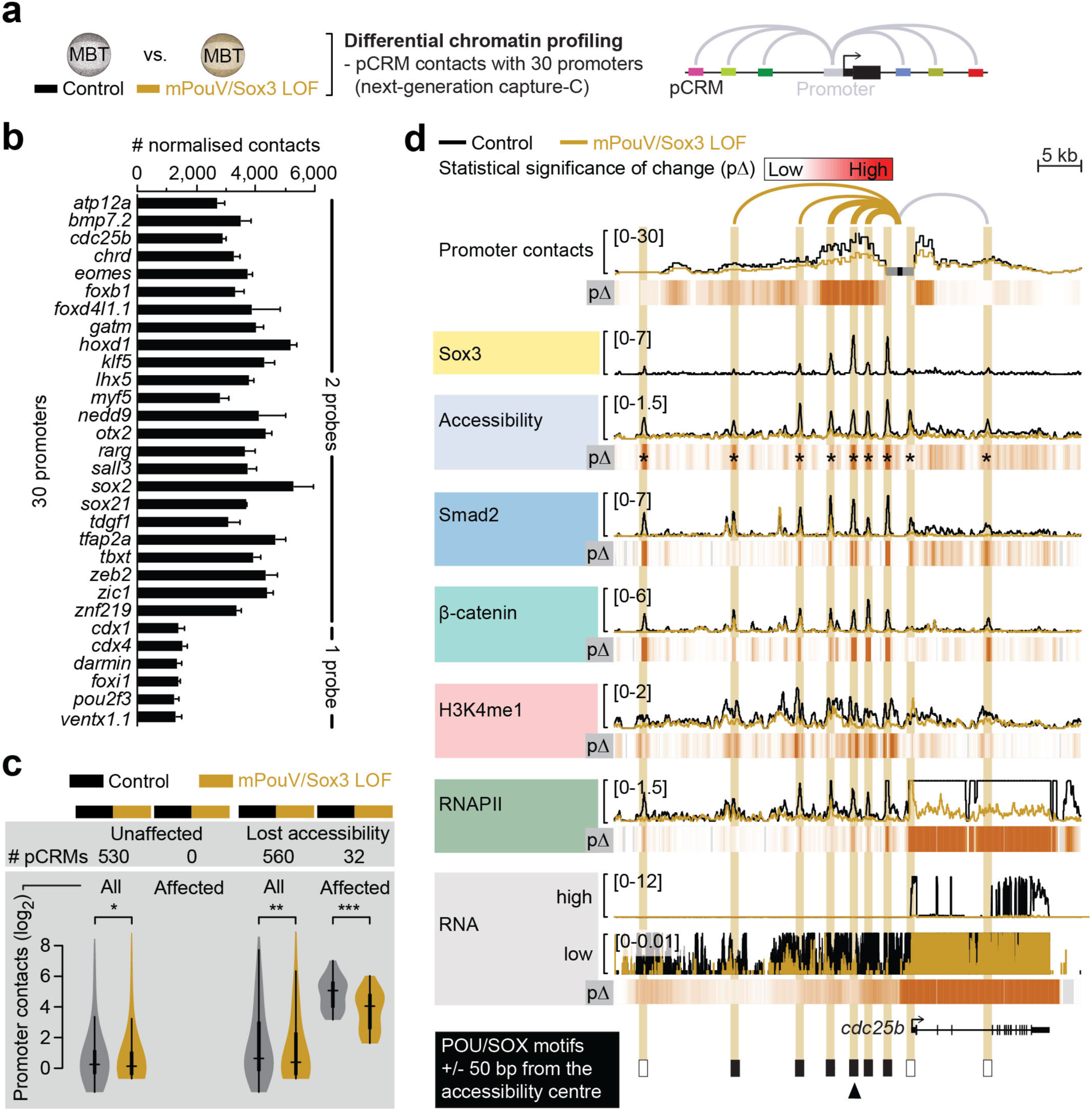
Pioneering activity of mPouV/Sox3 initiates extensive chromatin remodeling such as the chromatin looping of distal pCRMs with promoters. (a) Used approach to reveal the effect of mPouV/Sox3 on chromatin conformation between 30 promoters and distal pCRMs (next-generation capture-C; n=3) at MBT. (b) Bar graph shows the number of normalized contacts (mean+SD; n=6) derived from non-redundant capture-reporter FLASH^53^ reads (Supplementary Fig. 12b) for each promoter captured with one or two probes (Supplementary Table 10). (c) Violin plots compare the number of promoter contacts with accessible pCRMs between uninjected and mPouV/Sox3 LOF embryos. The comparison is stratified into pCRMs with stable and lost accessibility upon mPouV/Sox3 LOF and shown for both all and affected (FDR ≤10%) promoter contacts. Wilcoxon tests and effect size estimates: *, p=5×10^−5^ and r_effect_=0.12 (small effect); **, p=5×10^−30^ and r_effect_=0.34 (medium effect); and ***, p=5×10^−10^ and r_effect_=0.78 (large effect). (d) Superimposed line tracks show the level of promoter-tied chromatin conformations, chromatin accessibility and various chromatin components (Smad2, β-catenin, H3K4me1 and RNAPII) at the *cdc25b* gene locus between control (uninjected) and mPouV/Sox3 LOF embryos. The RNA track is split into a high (0-12) and low (0-0.01) expression window. Note that the low-expression window shows that locally transcribed non-coding super-enhancer RNA depend on mPouV/Sox3 as well as the gene *cdc25b*. Heat maps (pΔ) below each superimposed line plot show the statistical significance of changes caused by mPouV/Sox3 LOF. The footer highlights the occurrences of canonical POU/SOX motifs (black filled rectangles) at accessible pCRMs (±50 bp from the accessibility centre) and one strongly affected pCRM with an arrowhead. Asterisks on the pΔ heat map mark significant (FDR ≤10%) reductions to pCRM accessibility. pCRMs are boxed in and their frequency of contacts with the *cdc25b* promoter are illustrated with an arc of varying strength. Boxes of affected pCRM and arcs of promoter contacts are colored orange.

The loss of accessibility triggered further profound changes to chromatin *in situ* (see arrowheads at bottom of **Figs. 9d** and **10a** and **Supplementary Figs. 13** and **14** for strongly affected pCRMs). First, the deposition of H3K4me1, RNAPII and signal mediators Smad2 and β-catenin was substantially reduced (**Fig. 8d** and **9d** and **Supplementary Fig. 13** and **14**). Second, in contrast to unaffected pCRMs, 32 pCRMs with compromised accessibility—most of which are part of super-enhancers—also showed significantly (FDR ≤10%) reduced promoter contacts (e.g., *tbxt*, *foxb1*, *cdc25b* and *zic1*) (**Figs. 9c,d** and **10a**, **Supplementary Figs. 13** and **14** and **Supplementary Table 12**). Third, at the transcriptional level, lower usage of (clustered) pCRMs coincided with the reduction of coding (**Fig. 8g**) as well as local non-coding RNA (last column in **Fig. 8d** and ‘low RNA’ track in **Fig. 9d** and **Supplementary Figs. 13** and **14**). Importantly, the profiling of LOF embryos revealed how chromatin predetermines signal induction and why, for instance, Nodal-induced transcription of *tbxt* was strongly affected by the loss of mPouV/Sox3 and that of *eomes* less so. In contrast to those of *eomes*, all promoter-tied pCRMs (super-enhancer) of *tbxt* contain canonical POU/SOX motifs, so were not accessible to Smad2 in mPouV/Sox3 LOF embryos (**Fig. 10a,b**). Thus, Smad2 interactions with critical Nodal responsive CRMs of *eomes* remained intact, but those of *tbxt* were impeded by compacted chromatin. Pioneer-initiated competence also applied to Wnt signaling with β-catenin failing to engage with Wnt responsive CRMs of *foxb1* in the absence of mPouV/Sox3 (**Supplementary Fig. 13**). Other transcriptional deficiencies in signal-responsive or non-responsive genes that were triggered by mPouV/Sox3 LOF also coincided with significant reductions (FDR ≤10%) in chromatin accessibility (**Fig. 10c** and **Supplementary Figs. 15**, **16** and **17**), suggesting that mPouV/Sox3 facilitates expression by making critical clusters of CRMs accessible to signal and other transcriptional mediators.

**Figure 10.**
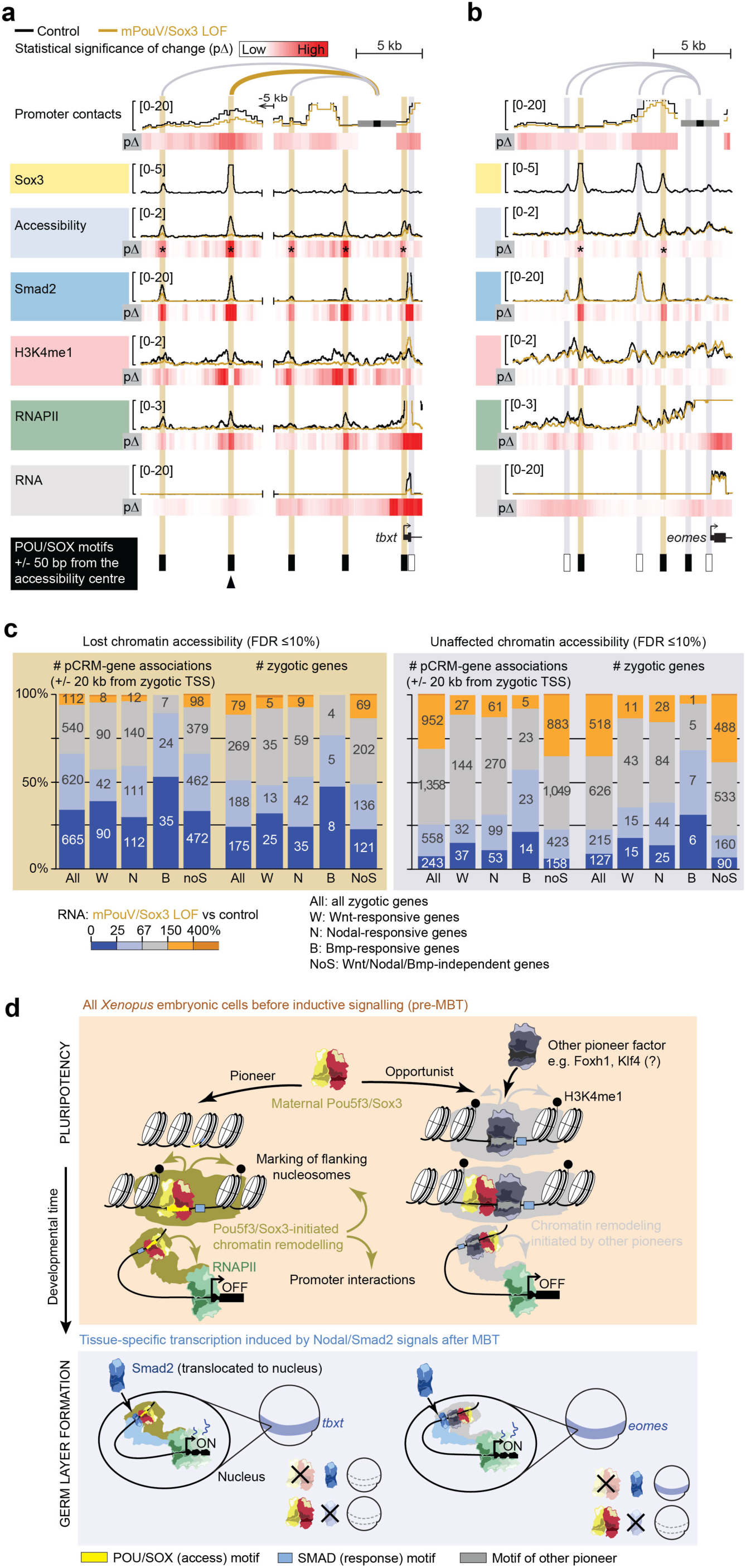
Maternal Pou5f3/Sox3-initiated chromatin remodeling to prime the first transcriptional response to inductive signals during ZGA. (a,b) Superimposed line tracks show the level of promoter-tied chromatin conformations, chromatin accessibility and various chromatin components (Smad2, H3K4me1 and RNAPII) at the Nodal-responsive mesoderm determinants *tbxt* (a) and *eomes* (b) between control (uninjected) and mPouV/Sox3 LOF embryos. Heat maps (pΔ) below each superimposed line track show the statistical significance of changes caused by mPouV/Sox3 LOF. The footer highlights the occurrences of canonical POU/SOX motifs (black filled rectangles) at accessible pCRMs (±50 bp from the accessibility centre) and one strongly affected pCRM with an arrowhead. Asterisks on the pΔ heat map mark significant (FDR ≤10%) reductions to pCRM accessibility. pCRMs are boxed in and their frequency of contacts with the promoter are illustrated with an arc of varying strength. Boxes of affected pCRM and arcs of promoter contacts are colored orange. (c) Stacked bar graphs summarize the correlation of unaffected and significantly reduced (FDR ≤10%) pCRM accessibility with RNAPII-mediated expression of all, signal responsive and non-responsive zygotic genes in mPouV/Sox3 LOF embryos at MBT. These correlations and corresponding numbers (placed on the stacked bars) are shown for pCRM-gene associations and zygotic genes. TSS-centric maps of reduced chromatin accessibility are shown for signal responsive and non-responsive genes in Supplementary Figs. 15-17. (d) Model of chromatin pioneering and opportunistic engagement to predefine first zygotic responses to inductive signals.

## DISCUSSION

Our results allow us to propose a model (**Fig. 10d**) of pioneer-initiated chromatin remodeling, or priming, that unlocks context-specific CRMs, some of which contain signal-responsive elements that enhance and regionalize transcription. Our observations are in line with recent reports of CRM priming for tissue-specific gene expression^34^ and signal interpretation^35, 36^. Thus, cell type-specific TFs pre-determine signal interpretation by shifting the binding of signal mediators and so causing the induction of different genes. These, ultimately, result in different cell fate transitions. Developmental context is imprinted on chromatin by co-expressed sequence-specific TFs recognizing their canonical DNA binding motifs in compacted chromatin. Based on *in vitro* experiments^37^, partial DNA motifs exposed on nucleosome surfaces can be enough to initiate chromatin access. Subsequent displacement of nucleosomes may be driven by cooperation among several TFs and the higher affinity of TFs for free genomic DNA. In the context of establishing pluripotency before MBT, we show that the pioneering activity of maternal Pou5f3 (mPouV) and Sox3 recognizing their POU/SOX motifs (**Fig. 10d**, left branch in top panel) triggers extensive chromatin remodeling *in situ*. This includes the H3K4me1 marking of CRM-flanking nucleosomes and chromatin looping with promoters. Newly primed CRMs can initiate further canonical as well as opportunistic (sequence-nonspecific) binding. Thus, irrespective of their preference for certain DNA sequences, chromatin factors (e.g. TFs, signal mediators, RNAPII) can appear on accessible CRMs with no canonical DNA binding motifs (**Fig. 10d**, right branch in top panel). However, the opportunistic recruitment of mPouV and Sox3, at least, seems to have little to no effect on changing the chromatin landscape and its transcriptional readout. This may be also true for other TFs as shown for ectopic MyoD in our study. Such recruitment behaviors can be difficult to detect without multi-level analysis of chromatin under LOF conditions. Overall, the canonical binding of mPouV/Sox3 strongly contributes to the pioneering of two fifths of all accessible pCRMs at MBT which permits or instructs one quarter of the ZGA (**Fig. 8b,g**). We often observe that the usage of pCRMs also coincides with the low-level activation of proximal non-coding RNA.

Although we have not performed chromatin profiling at a single-cell level, we suggest for three reasons that this priming occurs in every embryonic cell before the onset of regional Wnt, Nodal or BMP signaling. First, the nuclear accumulation of maternal TFs (e.g., mPouV, Sox3, Foxh1 and mVegT) occurs ubiquitously (or within the vegetal hemisphere in case of mVegT) and precedes that of signal mediators. Second, mPouV/Sox3 LOF affects chromatin similarly at zygotic genes with (e.g. *foxb1*) or without (e.g. *cdc25b*) tissue-specific expression. And finally, dissected animal caps replicate the *in vivo* response of the marginal zone tissue to Nodal and Wnt signaling. Importantly, and consistent with the idea of context-dependent signal interpretation, our findings suggest that signal mediators have inferior pioneering activity and thus rely on cooperating pioneer factors like mPouV and Sox3 to make their signal response elements accessible. Analysis of the related genes *eomes* and *tbxt*, both of which are induced by Nodal signaling, but with only *tbxt* depending on mPouV/Sox3, reveals that the same context-dependent competence can be differentially encoded in the genome (through the presence or absence of POU/SOX motifs in signal-responsive CRMs) (**Fig. 10d**, lower panel). Based on transcriptional dependencies shown here and in the spinal cord^38^, this regulation of competence is complemented by other TFs like mVegT and also applies to Wnt signaling and other signaling pathways such as BMP and Sonic hedgehog.

Our work has focused on the pioneering activity of mPouV and Sox3. The enrichment of DNA recognition motifs at pCRMs in use before the MBT suggests that only a few additional maternal TF families, including those of T-box, FoxH and Krüppel-like ZF families, are involved in the bulk acquisition of competence at the MBT (**Fig. 2b**). Indeed, many of these TF family members are rapidly turned over and are most abundant around the MBT^39, 40^, suggesting dynamic and strong TF activity^41^. Their activity generates a signature of accessible DNA motifs of pluripotency or early cell lineage determination that is conserved among vertebrates and is reminiscent of the chromatin footprints in embryonic stem cells *in vitro*^42^. Remarkably, as in the human genome^42^, POU/SOX motifs populate distal CRMs, while Krüppel ZF motifs are frequently found in promoter-proximal CRMs (**Fig. 8f**). This creates a functional separation among the pluripotency factors where PouV/SoxB1 and Krüppel-like factors remodel enhancers and promoters, respectively. Collectively, we show that mPouV/Sox3 predetermine cell fate decisions by initiating access to signal-responsive CRMs via POU/SOX motifs. The use of these permissive CRMs is most prominent in pluripotent cells^24, 43^ and proves to be critical *in vivo* to the induction of zygotic genes with functions in germ layer formation and primary axis determination.

The findings presented here confirm that core pluripotency TFs contribute substantially to ZGA as previously reported for SoxB1, PouV and Nanog TFs for zebrafish ZGA^44, 45^. However, in contrast to zebrafish or humans, a true *nanog* gene has not been found in the *Xenopus* genome, which has led to the suggestion that zygotic Ventx TFs with their Nanog-like DNA binding characteristics could operate as the gatekeepers of pluripotency instead^46^. This notwithstanding, we anticipate that signal competence is controlled across vertebrate embryonic stem cells in a manner resembling that in the early *Xenopus* embryo.

The approach we adopt here can be applied to any other cell type to discover the molecular basis of its competence. Ultimately, this will generate a lexicon of competence that outlines which pioneer factors unlock which (signal-responsive) CRMs and gene loci. For example, the motif compositions of CRMs later engaged in the *Xenopus* tailbud embryo points at the potential of OTX and CDX TFs in conveying competence in anterior and posterior compartments, respectively (**Fig. 5c**). Interestingly, various tumors are associated with the mis-expression of pioneer factors^47^ and thus may display different competencies from the surrounding host tissue. Conversely, knowing which pioneers are required to permit a specific signal response will increase the success of engineering patient-specific tissue for transplantation therapy. On a broader level, our profiling of chromatin states under LOF conditions discriminates functional from non-functional binding and provides a promising avenue for deciphering the non-coding part of the genome for basic and therapeutic research.

## METHODS

Methods and any associated references are available in the online version of the paper.

### Accession code

Sequencing data are deposited in the Gene Expression Omnibus (GEO) database with accession number GSE113186.

## ACKNOWLEDGMENTS

We thank Abdul Sesay, Leena Bhaw, Harsha Jani, Deborah Jackson and Meena Anissi for deep sequencing; Mike Klymkowsky, Manolis Gialitakis and John Gurdon for supplying antibodies; Stefan Hoppler and Yukio Nakamara for discussing β-catenin ChIP protocol; Jim Hughes and Damien Downes for sharing pre-publication results of an improved next-generation capture-C method; Ying Wang for providing sequence conservation tracks; Mareike Thompson, Rita Monteiro and Clara Collart for critical reading of the manuscript; and the Smith lab for discussions and advice. G.E.G and J.C.S. were supported by the Medical Research Council (program number U117597140) and are now supported by the Francis Crick Institute, which receives its core funding from Cancer Research UK (FC001-157), the UK Medical Research Council (FC001-157), and the Wellcome Trust (FC001-157).

## AUTHOR CONTRIBUTIONS

Conceptualization, G.E.G.; Methodology, G.E.G.; Software, G.E.G.; Formal Analysis, G.E.G. and N.D.L.O.; Investigation, G.E.G. and T.S.; Writing – Original Draft, G.E.G. and J.C.S.; Writing – Review & Editing, G.E.G and J.C.S.; Funding Acquisition, G.E.G. and J.C.S.; Supervision, G.E.G. and J.C.S.

## COMPETING FINANCIAL INTERESTS

The authors declare no competing financial interests.

## Supplementary information

**Supplementary Figure 1.**
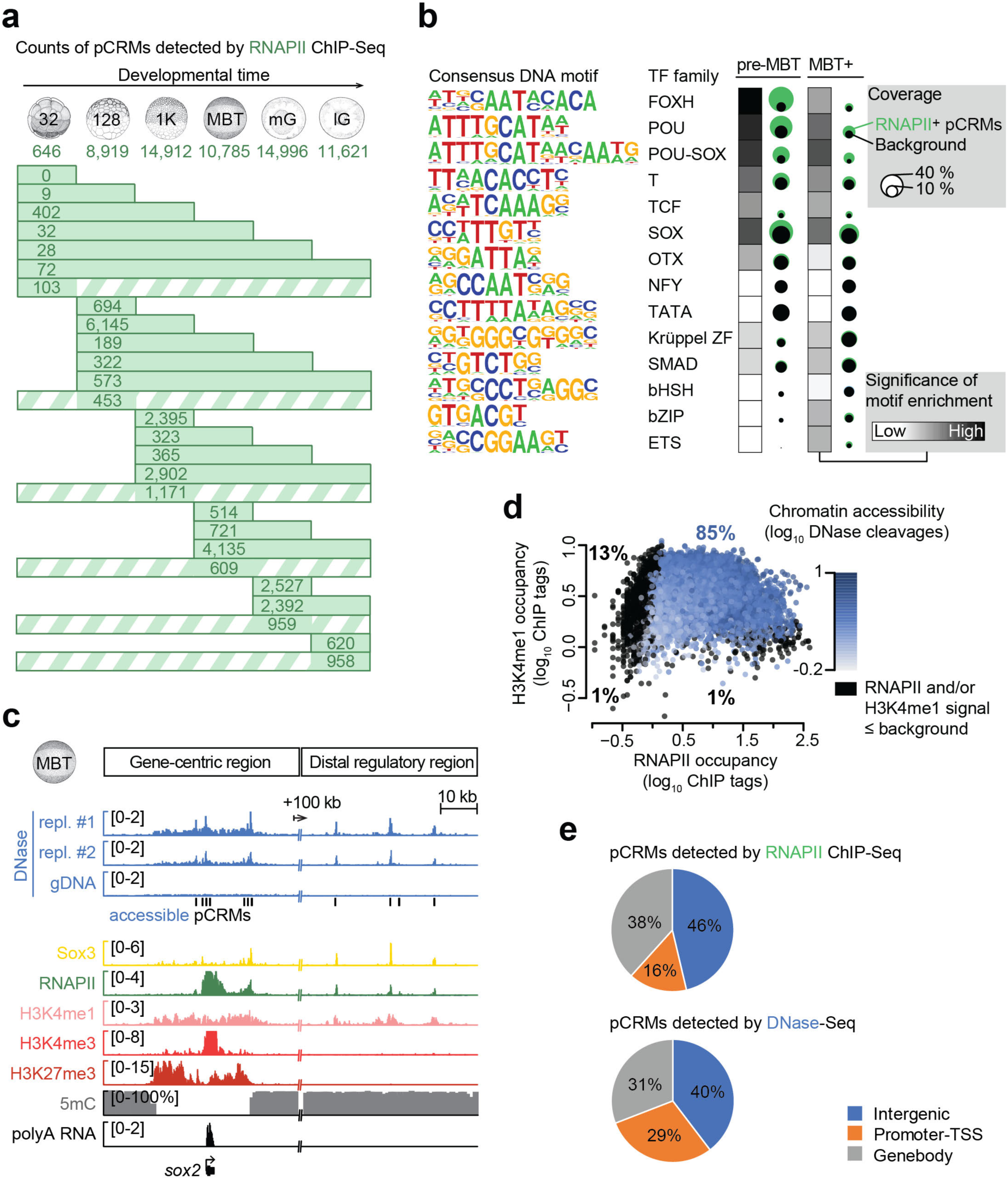
Characterization of pCRMs instructing ZGA. (**a**) Graph shows the temporal dynamics of RNAPII-engaged (RNAPII^+^) pCRMs (≥1 ChIP tag/million) from the 32-cell to the late gastrula stage. Striped bars refer to RNAPII^+^ pCRMs fragmentarily detected across consecutive developmental stages. Abbreviations used for the developmental timeline: 32, 128 and 1K, 32-, 128- and 1,024-cell stage; MBT, mid-blastula transition; mG and lG, mid- and late gastrula stage. (**b**) Heat map and bubble plot show the respective statistical significance and coverage of DNA motifs enriched among the top 5,000 RNAPII^+^ pCRMs detected from the 32-cell to the 1,024-cell stage (pre-MBT) and from the MBT to the late gastrula stage (MBT+). This analysis included 39,785 (pre-MBT) and 41,898 (MBT+) genomic ‘background’ regions matching the overall GC contents of the selected RNAPII^+^ pCRMs. (**c**) Validation of our DNase-Seq method: DNase-probed chromatin accessibility (biological replicates #1 and #2) at the *sox2* locus (gene-centric region and downstream distal regulatory region) is shown alongside with various other chromatin features (Sox3, RNAPII, H3K4me1, poly(A) RNA from this study and H3K4me3, H3K27me3, 5-methylcytosine [5mC] from ref. ^54^) detected around the MBT. DNase-treated naked genomic DNA (gDNA) was used as a negative control for DNase-probed chromatin accessibility. (**d**) Biplot shows the DNA occupancy levels of RNAPII and H3K4me1 at accessible pCRMs. pCRMs (dots) are colored according to their chromatin accessibility except for the pCRMs that showed RNAPII and/or H3K4me1 signals piled up over a distance of 1 kb below background. (**e**) Pie charts show the genomic distribution of pCRMs detected by RNAPII ChIP-Seq and DNase-Seq, respectively.

**Supplementary Figure 2.**
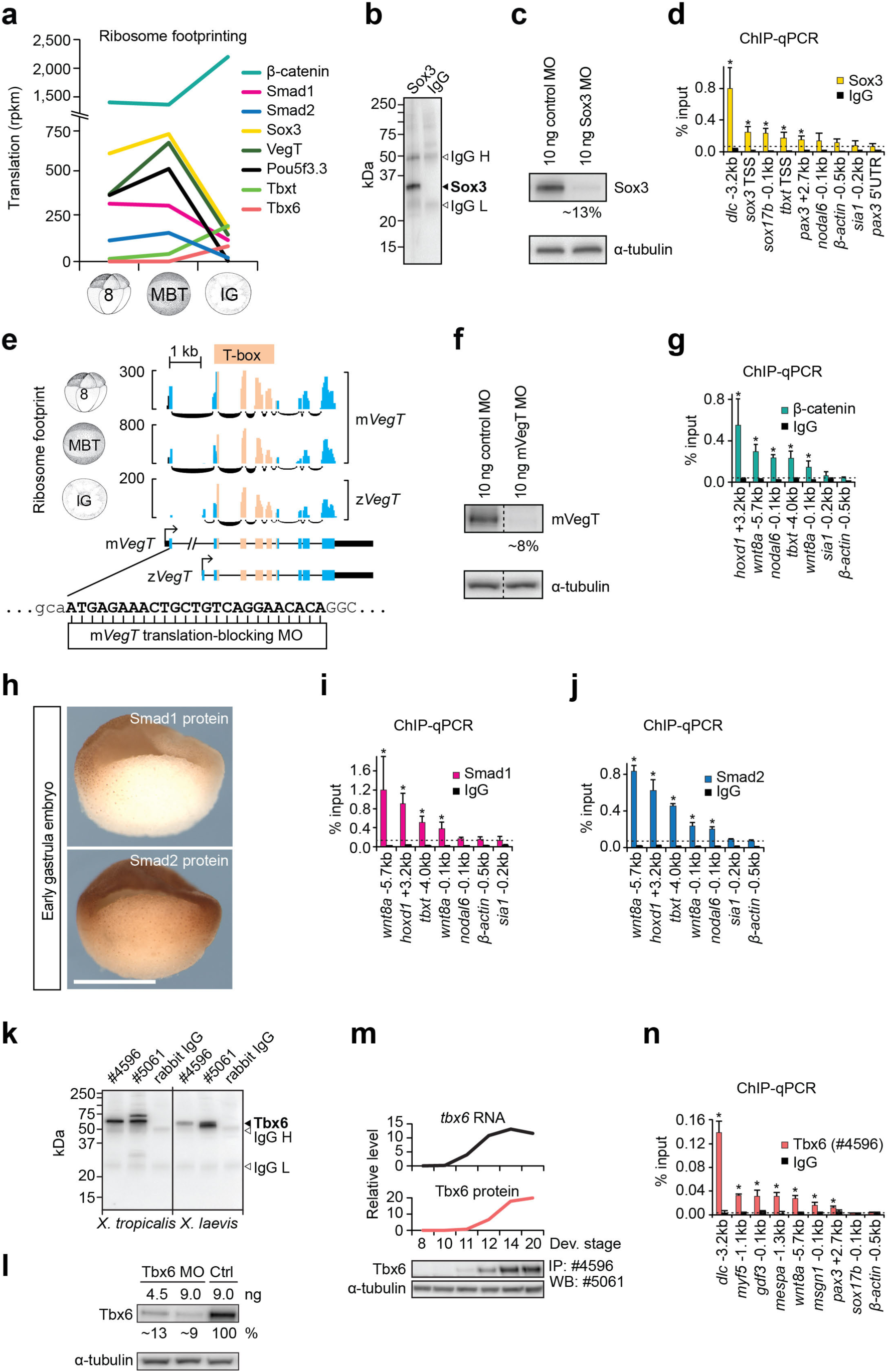
Verification of antibodies against frequently translated TFs and signal mediators. (**a**) Line chart shows the level of ribosome footprints^13^ of selected TFs and signal mediators across MBT in rpkm (reads per kilobase of transcript per million mapped reads). Abbreviations used for the developmental timeline: 8, 8-cell stage; MBT, mid-blastula transition; lG, late gastrula stage. (**b**) Western blot shows the immunoprecipitation (IP) of Sox3 protein extracted from early gastrula embryos. IgG H and L, detected IgG heavy and light chains of the IP antibody. (**c**) Western blot shows the level of Sox3 protein immunoprecipitated from control (control MO) and Sox3-depleted (Sox3 MO) blastula embryos (Sox3 LOF). α-tubulin, IP input control. (**d**,**g**,**i**,**j**,**n**) Bar graphs show the ChIP-qPCR results as a percentage of ChIP input (mean+SD; n=2-3) for Sox3, β-catenin, Smad1 and Smad2 at the early gastrula stage and for Tbx6 at the early neurula stage. One-tailed Student’s t-test (comparing to IgG control): *, p ≤0.1 and ≥2-fold enrichment relative to the lowest DNA recovery with the ChIP antibody (dotted line). (**e**) Ribosome footprinting tracks show the post-MBT switch of translation from the maternal (m) to the zygotic (z) *VegT* transcript. The VegT translation-blocking MO was designed to block translation of the maternal transcript only. (**f**) Western blot shows the level of mVegT protein immunoprecipitated from control (control MO) and mVegT-depleted (mVegT MO) blastula embryos (mVegT LOF). Dotted line indicates the elimination of irrelevant lanes from the western blot. α-tubulin, IP input control. (**h**) WMIHC shows the spatial distribution of nuclear Smad1 and Smad2 protein on bisected early gastrula embryos. Scale bar, 0.5 mm. (**k**) Western blot shows the level of Tbx6 immunoprecipitated from *X. tropicalis* and *X. laevis* late gastrula embryos. Antibody #4596 was chosen for subsequent IP and ChIP experiments, and #5061 for Western blotting. (**l**) Western blot (WB) shows the level of Tbx6 protein immunoprecipitated from standard control and Tbx6 morphants at the early tailbud stage. α-tubulin, IP input control. (**m**) Line charts show the relative level of *tbx6* transcripts (RT-qPCR) and Tbx6 protein (IP/WB) from the early blastula to the early tailbud stage. α-tubulin, IP input control.

**Supplementary Figure 3.**
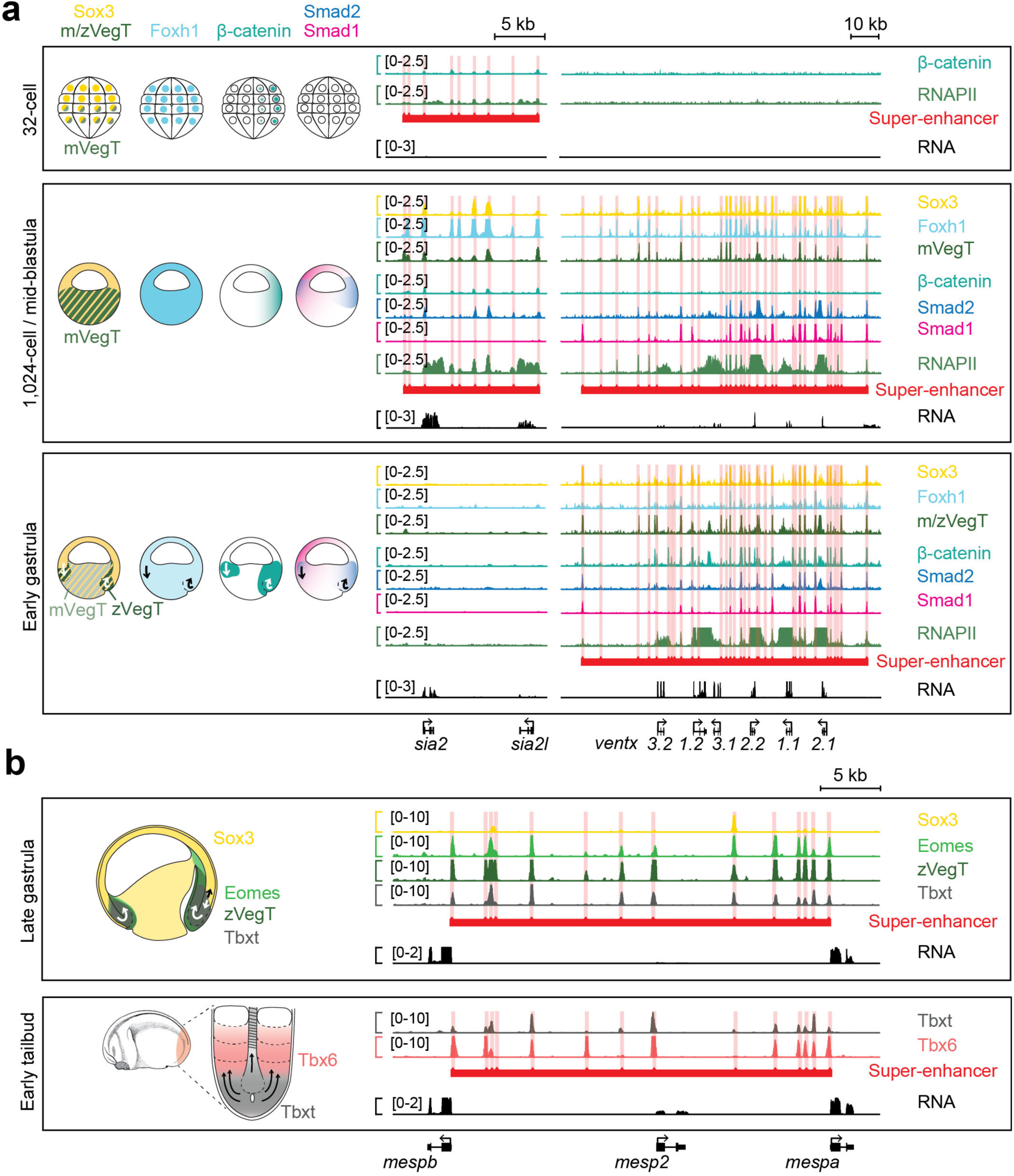
Snapshots of TF and signal mediator binding to super-enhancers during early embryogenesis. (**a**) Dynamic chromatin engagement of endogenous Sox3, Foxh1^18^, VegT, β-catenin, Smad2, Smad1 and RNAPII to putative super-enhancers of the *siamois* and *ventx* gene cluster from the 32-cell to the early gastrula stage. Illustrated sagittal sections (dorsal side is right) show the nuclear localization of the selected TFs with arrows pointing to the tissue movements of gastrulation. (**b**) Snapshot of endogenous Sox3, Eomes, zVegT, Tbxt and Tbx6 binding to the putative super-enhancer of the *mesp* gene cluster at the late gastrula and/or early tailbud stage. Illustrated sagittal sections (dorsal side is right) show the nuclear localization of selected TFs in the late gastrula embryo and at the caudal end of the early tailbud embryo with arrows pointing to the tissue movements during axial elongation^20^. All super-enhancers were formed by stitching together engaged pCRMs that are ≤25 kb apart^21^.

**Supplementary Figure 4.**
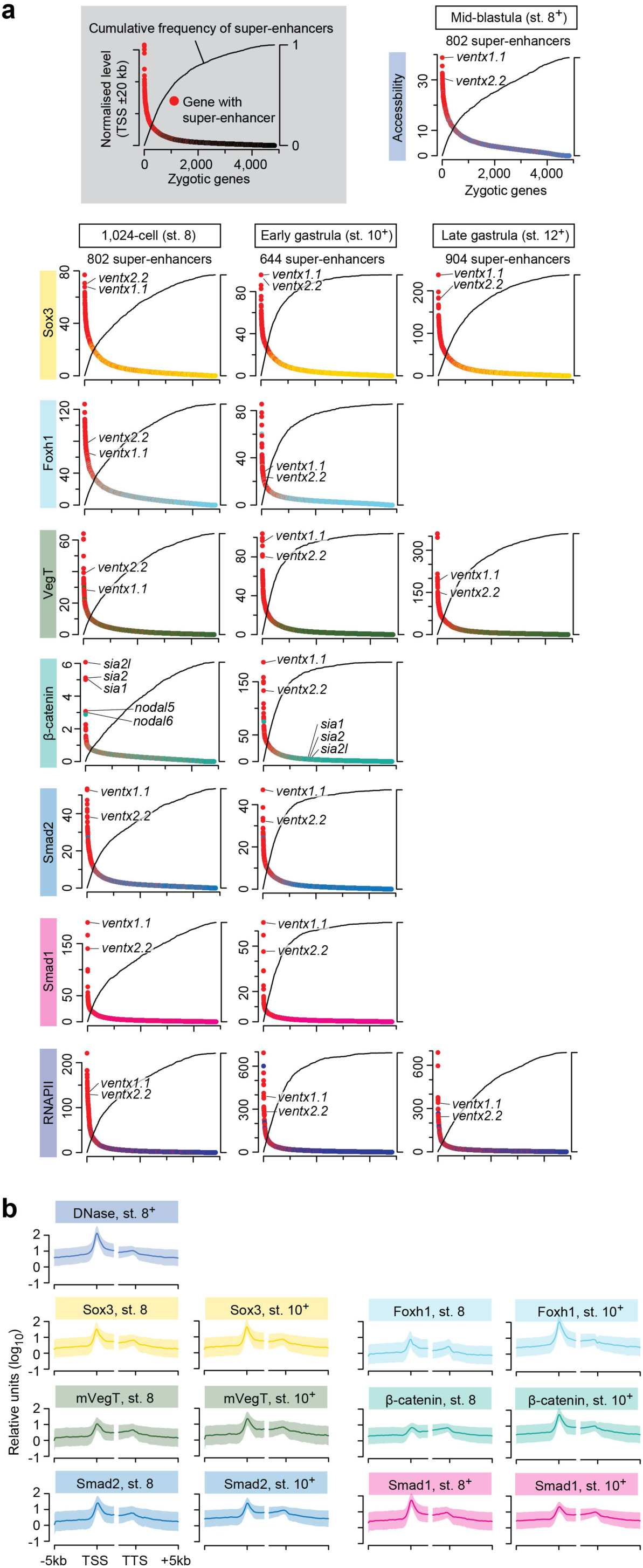
Characterization of genome-wide chromatin engagement and chromatin accessibility in early *Xenopus* embryos. (**a**) The grey box explains the composition of the following dot plots: the x-axis displays zygotic genes (detected by RNAPII profiling from the 32-cell to the late gastrula stage^52^) ranked by the total level of pCRM accessibilities or pCRM occupancies (normalized to 1 million mapped reads; primary y-axis) that are nearest (≤20 kb) to corresponding transcription start sites (TSSs). The secondary y-axis shows the cumulative frequency of gene-associated super-enhancers (≤5 kb from TSSs). Genes with super-enhancers are highlighted in red. (**b**) Meta-summary (mean±SD; log_10_ scale) of chromatin accessibility or DNA occupancy levels across zygotic TSSs and transcription termination sites (TTSs). For each developmental stage, super-enhancers were formed by stitching together pCRMs that are ≤25 kb apart and engaged by RNAPII or at least one TF or signal mediator.

**Supplementary Figure 5.**
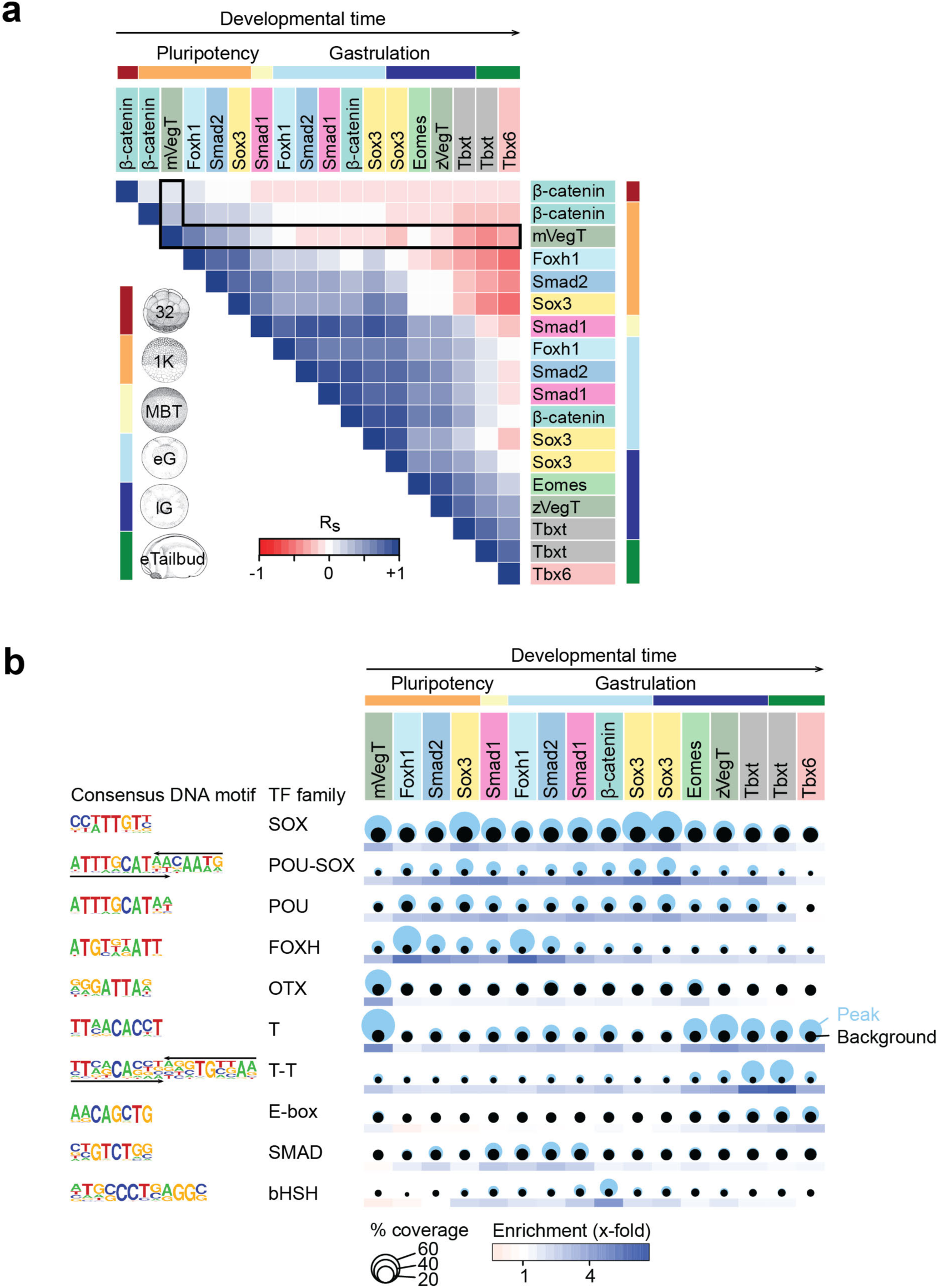
The relationship of chromatin recruitment among TFs and signal mediators and its underlying preferences for specific DNA recognition motifs in pCRMs of pluripotent and neuro-mesodermal embryonic cells. (**a**) Heat map shows pairwise Spearman correlations (R_s_) of DNA occupancy levels at ∼12,500 pCRMs among TFs (Sox3, mVegT, Foxh1^18, 25^, Eomes^19^, zVegT^19^, Tbxt^19^ and Tbx6) and signal mediators (β-catenin, Smad1, Smad2) at indicated developmental stages. The pCRM coordinates were generated by collating the strongest 2,000 peaks of each binding profile. Abbreviations used for the developmental timeline: 32 and 1K, 32- and 1,024-cell stage; MBT, mid-blastula transition; eG and lG, early and late gastrula stage; eTailbud, early tailbud stage. (**b**) Bubble plots and heat maps show the coverage and enrichment of DNA recognition motifs, respectively, among the top 2,000 pCRMs in each chromatin profile.

**Supplementary Figure 6.**
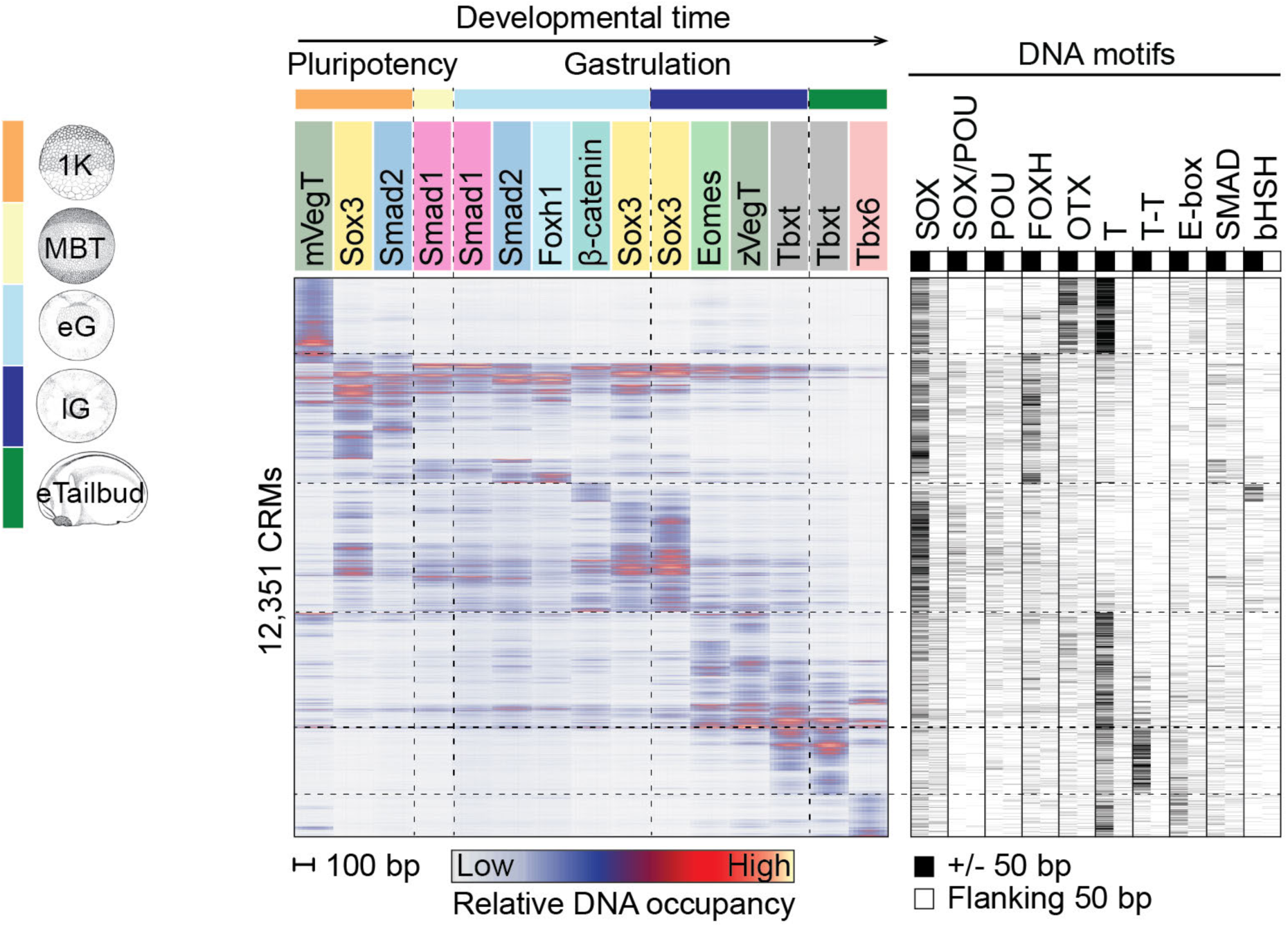
pCRM binding patterns correlate with the occurrence of specific DNA motifs. Heat map to the left shows the relative DNA occupancy levels across ∼12,500 pCRMs for a selection of TFs (mVegT, Sox3, Foxh1^25^, Eomes^19^, zVegT^19^, Tbxt^19^ and Tbx6) and signal mediators (β-catenin, Smad1, Smad2) from the 1,024-cell to the early tailbud stage. pCRM were hierarchically clustered according to the binding levels of all TFs and signal mediators. The pCRM coordinates were generated by collating the strongest 2,000 peaks of each binding profile. Heat map to the right shows the occurrence of specific DNA motifs ±50 bp from the pCRM centre as well as the flanking 50 bp. The consensus sequences of these DNA motifs are shown in **Supplementary Fig. 5b**. Abbreviations used for the developmental timeline: 1K, 1,024-cell stage; MBT, mid-blastula transition; eG and lG, early and late gastrula stage; eTailbud, early tailbud stage.

**Supplementary Figure 7.**
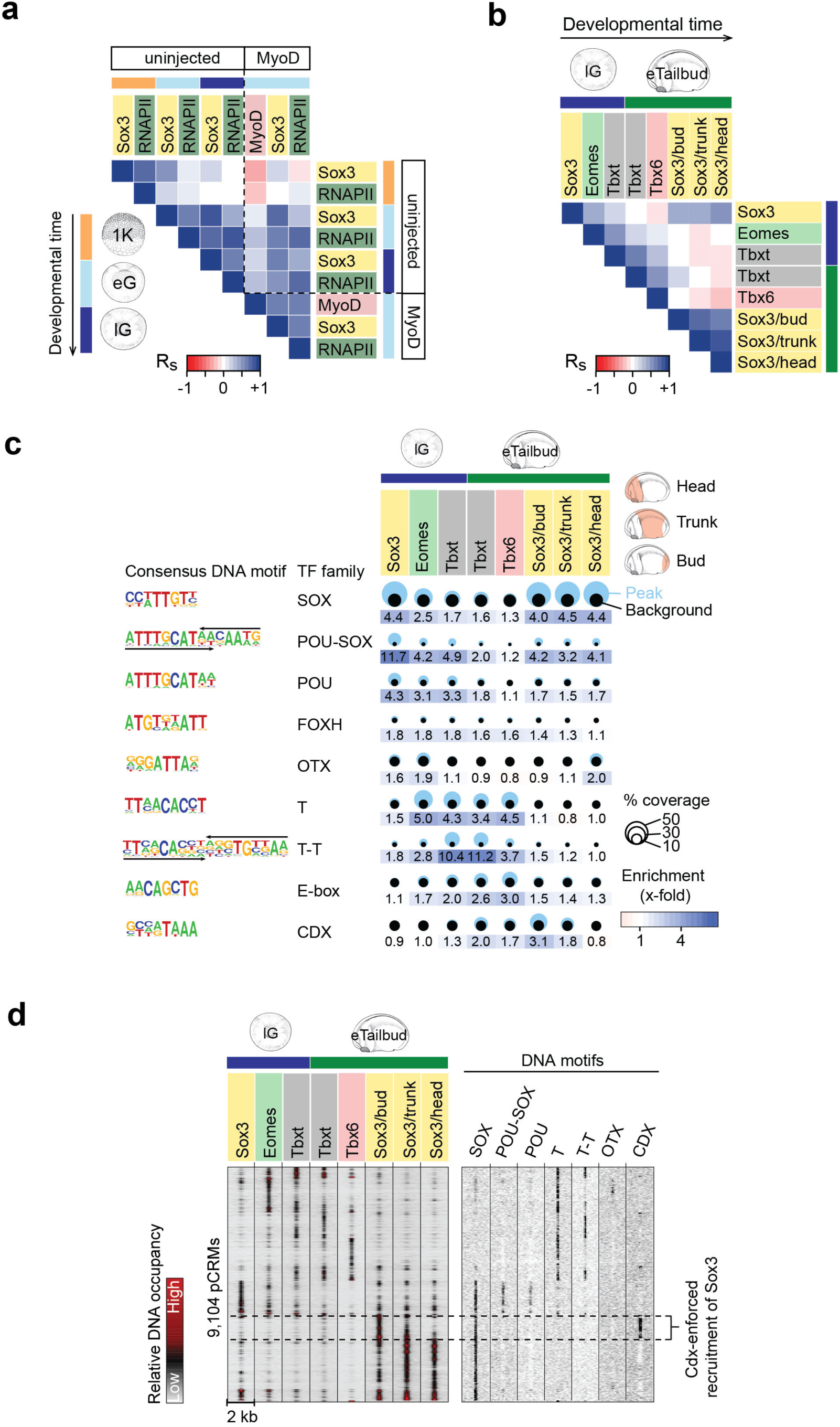
TF co-expression influences chromatin engagement. (**a**,**b**) Heat maps show Spearman correlations (R_s_) of DNA occupancy levels of indicated factors in different developmental contexts. (**c**) Bubble plot and heat map show the respective coverage and enrichment of DNA motifs in pCRMs bound by Sox3, Eomes^19^, Tbxt^19^ and Tbx6 in late gastrula and early tailbud embryos. For Sox3 profiling, the embryo was dissected into three parts: head, trunk and tailbud. (**d**) Heat map shows the hierarchical clustering of 9,104 pCRMs according to relative DNA occupancy levels. The pCRM coordinates were generated by collating the strongest 2,000 peaks of each binding profile. The pCRM coordinates were generated by collating the strongest 2,000 peaks of each binding profile. Abbreviations of the developmental timeline: 1K, 1,024-cell (early blastula); eG, early gastrula; lG, late gastrula; and eTailbud, early tailbud.

**Supplementary Figure 8.**
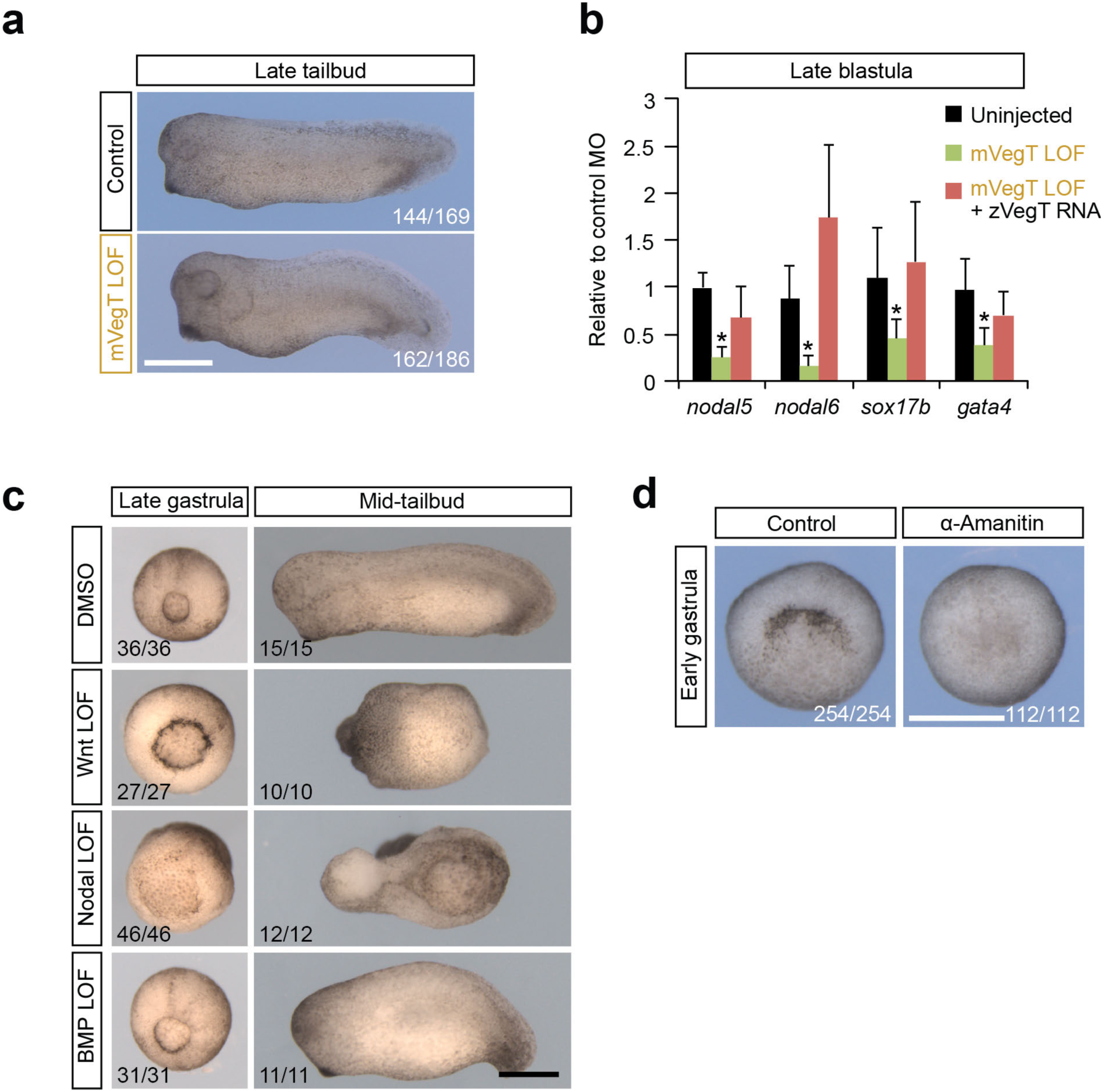
Morphological defects caused by the functional loss of maternal TFs and signal mediators (**a**) Morphological phenotype caused by the LOF of mVegT at the late tailbud stage. (**b**) Bar graph shows the RT-qPCR results of rescuing reduced *nodal5*, *nodal6*, *sox17b* and *gata4* transcript levels in mVegT LOF embryos by the injection of *X. tropicalis* z*VegT* mRNA. Error bars, mean+SD (n=5). Two-tailed Student’s t-test: *, p <0.02. (**c**) Morphological phenotypes caused by the single LOF of canonical Wnt, Nodal or BMP signaling when control embryos reached the late gastrula and mid-tailbud stage. (**d**) Morphological phenotype caused by the injection of α-amanitin when control embryos reached the early gastrula stage. Numbers in the right or left bottom corner of each image refer to the count of embryos detected with the displayed morphological phenotype. Scale bars, 0.5 mm.

**Supplementary Figure 9.**
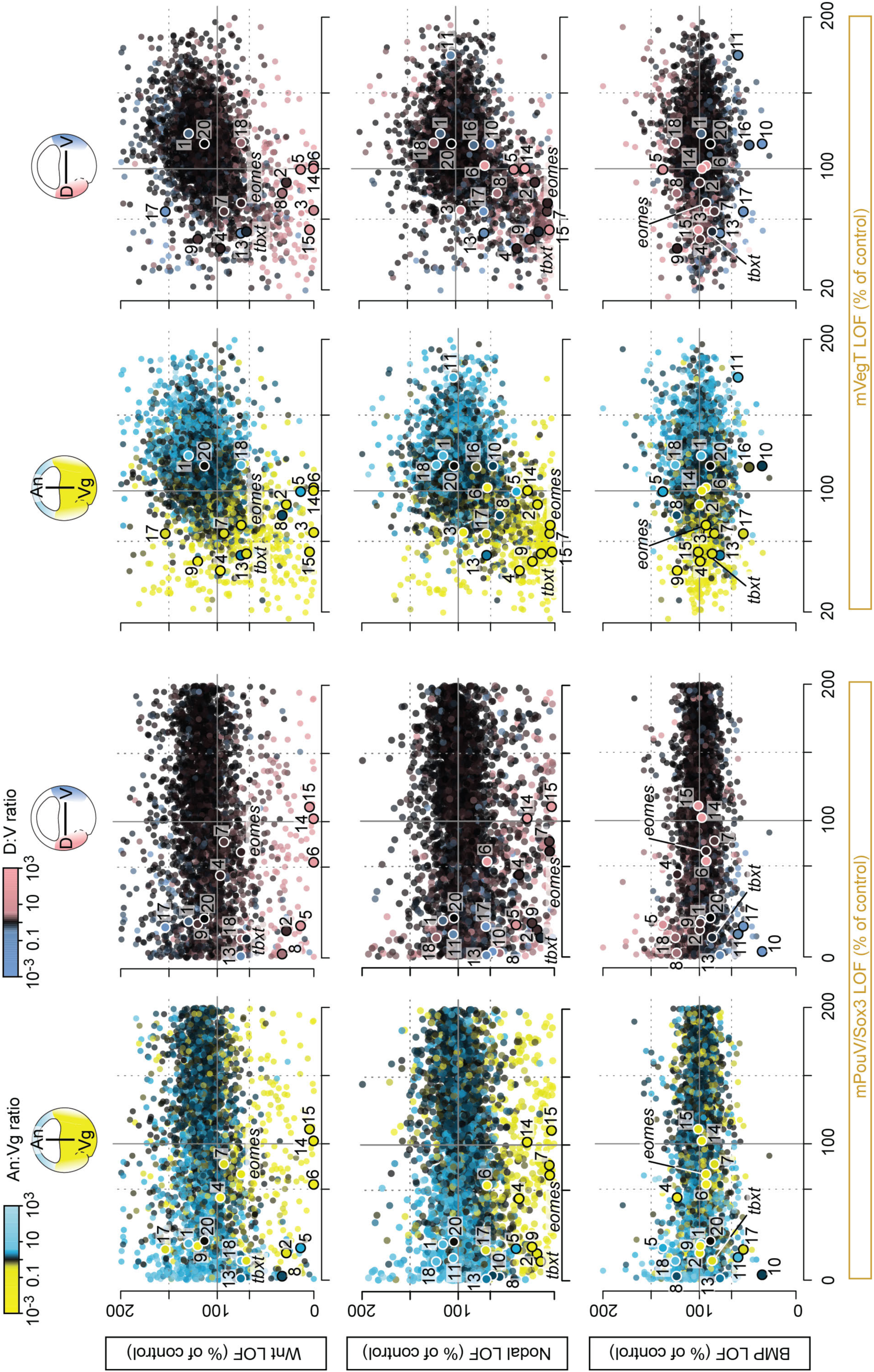
Transcriptional comparison of zygotic genes between indicated LOFs of maternal TFs or signals. Dots are colored according to the normal ratio of transcript levels (regional expression) across the animal-vegetal (An:Vg) or dorso-ventral (D:V) axis^51^. Numbered dots refer to genes listed in Fig. 6e.

**Supplementary Figure 10.**
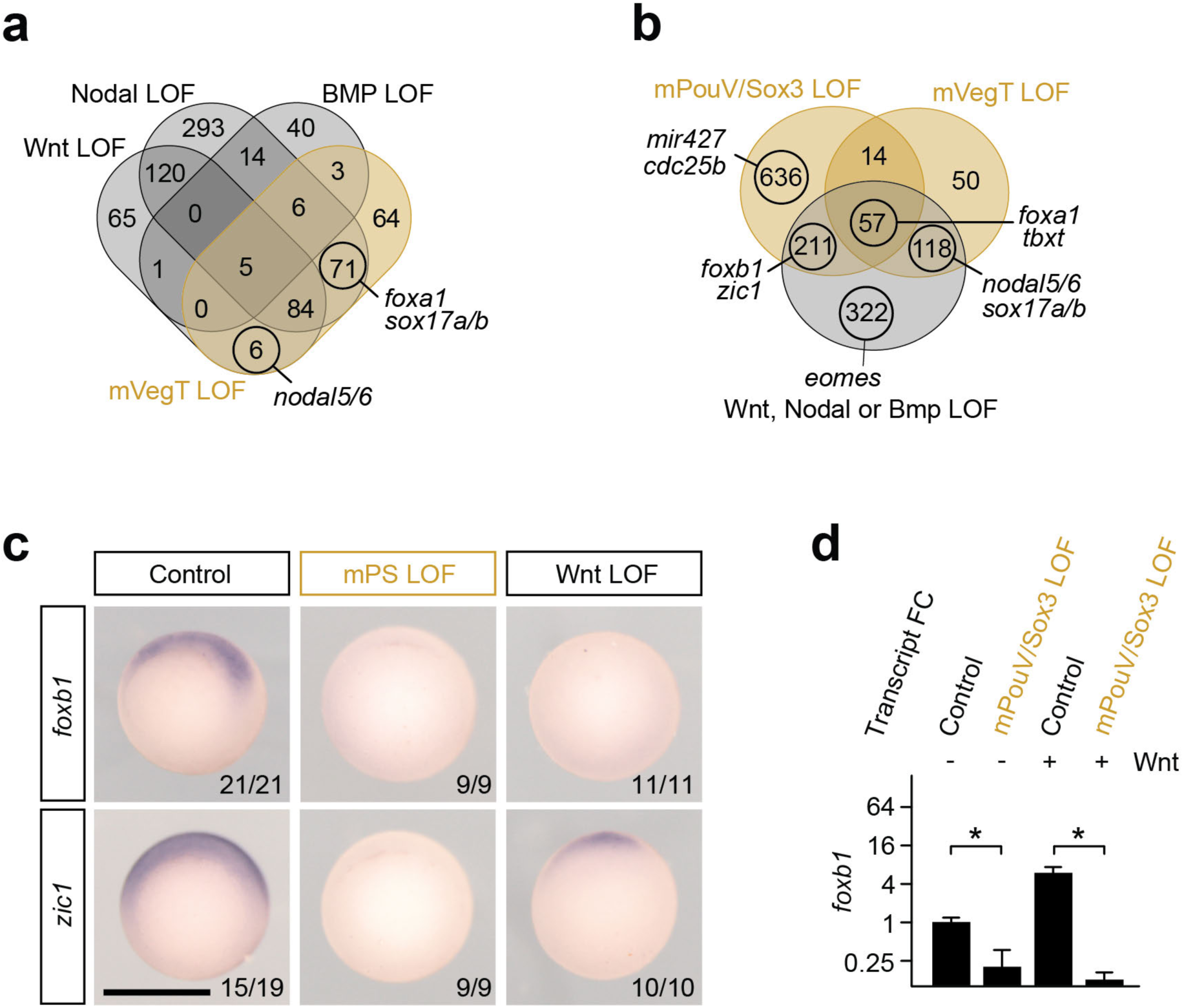
Signal-induced regionalization of ZGA depends on maternal TFs. (**a,b**) Venn diagrams show the number of genes downregulated by indicated LOFs. (**c**) Early gastrula-staged WMISHs show that Wnt-induced transcription of *foxb1* and *zic1* on the dorsal side of the embryo depends on ubiquitously expressed maternal pluripotency factors mPouV and Sox3 (mPS). Embryos are imaged from the vegetal side and orientated so that dorsal side faces top. Numbers in the right bottom corner of each image refer to the count of embryos detected with the displayed WMISH staining among all embryos analyzed per condition and *in situ* probe. Scale bars, 0.5 mm. (**d**) Bar graph shows the relative quantification of *foxb1* transcript levels (RT-qPCR) in control and mPouV/Sox3 LOF animal caps with or without canonical Wnt signaling. Error bars, mean+SD (n=2). Two-tailed Student’s t-test: *, p=0.05.

**Supplementary Figure 11.**
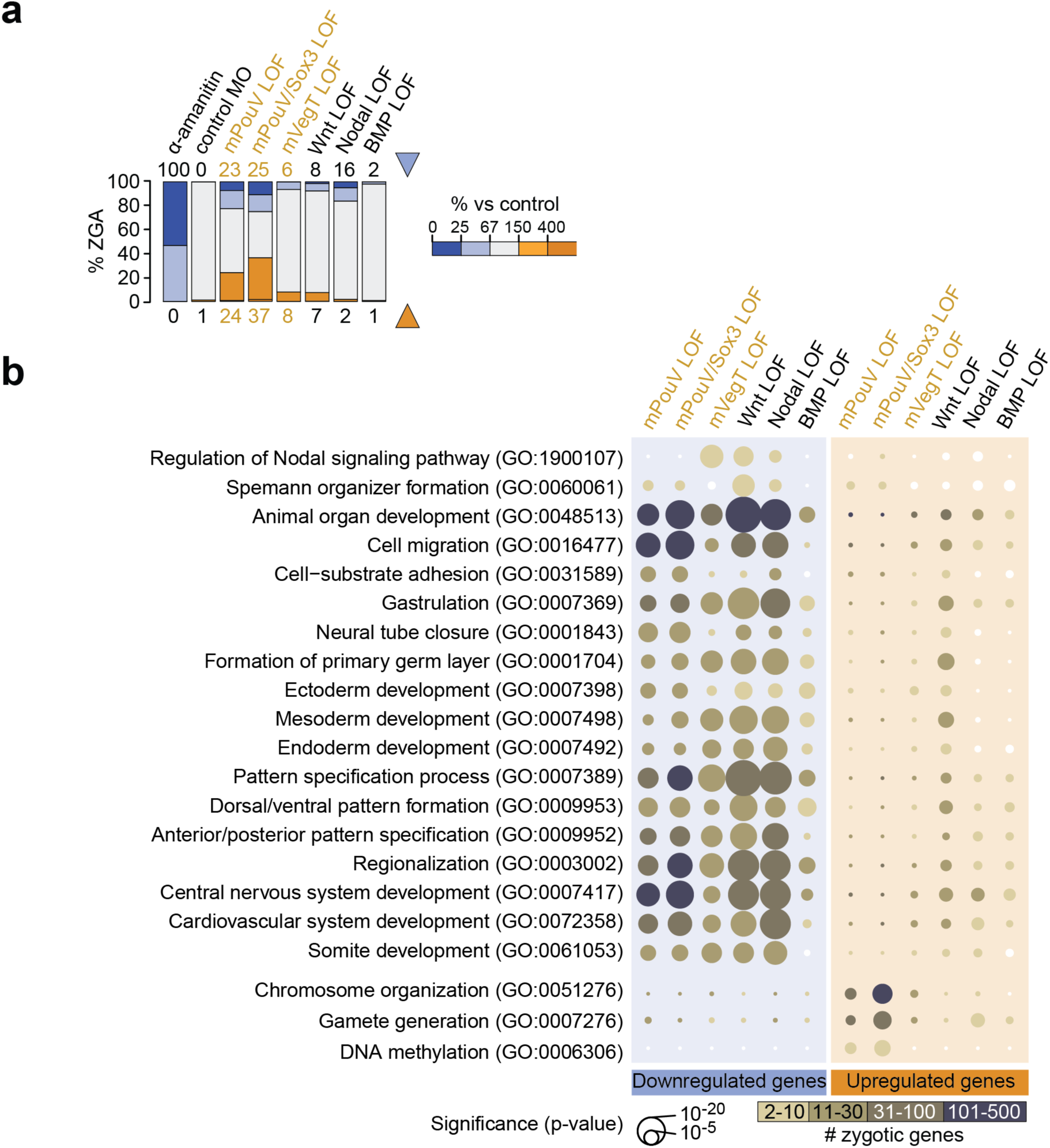
Selected maternal TFs and inductive signals regulate the regionalization of zygotic genome activation (ZGA) and the formation of primary body axes and germ layers. (**a**) Stacked bar graph shows the percentage of zygotic genes misregulated (% ZGA) by indicated LOFs. (**b**) Bubble plot shows the enrichment of key GO term-related biological processes among the zygotic genes misregulated by indicated LOFs. Size of bubble reflects the statistical significance (hypergeometric p-value) of enrichment while the color indicates the number of affected genes.

**Supplementary Figure 12.**
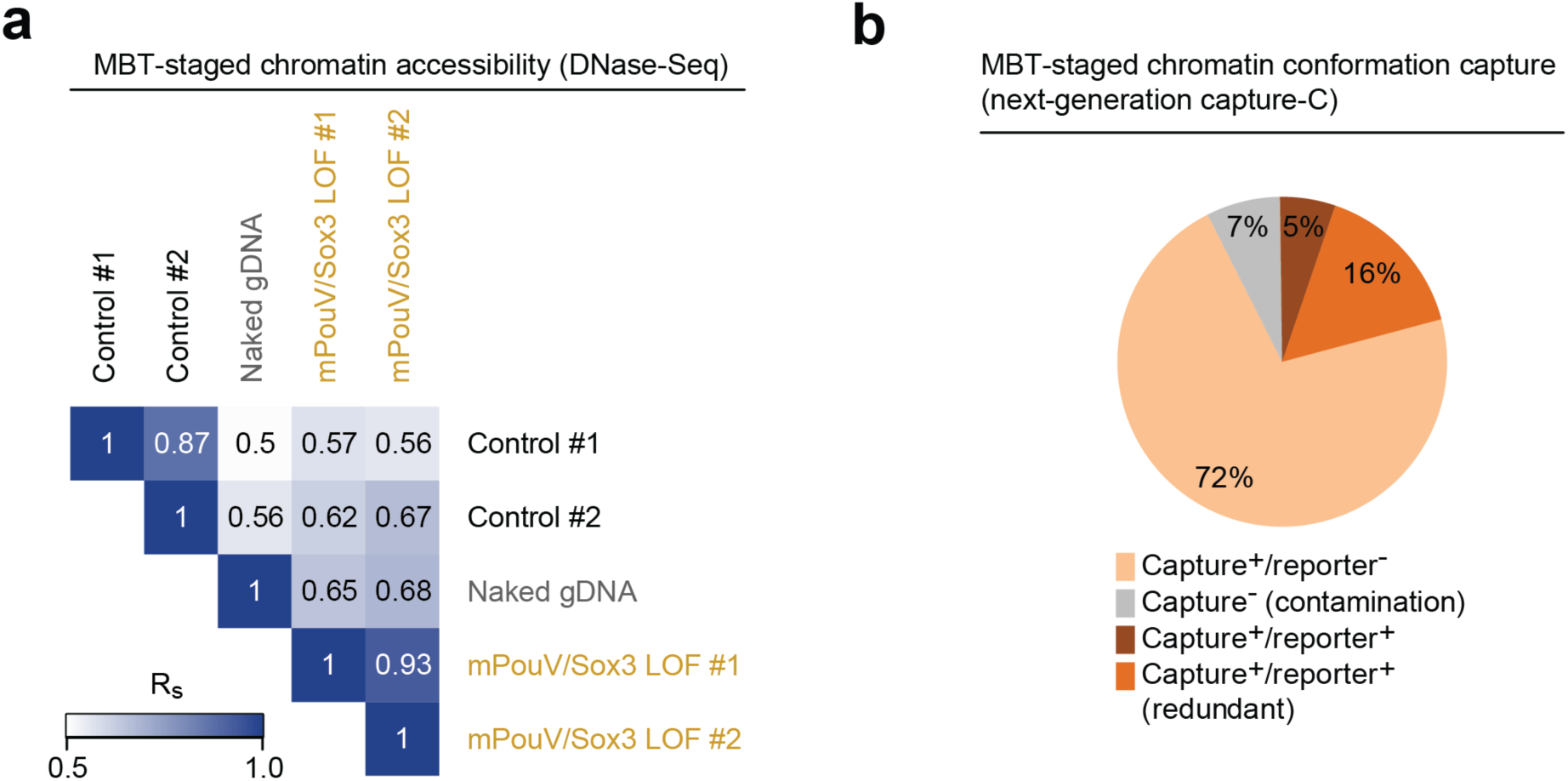
Performance of DNase-Seq and next-generation capture-C. (**a**) Heat map shows pairwise Spearman correlations (R_s_) among indicated biological replicates of MBT-staged DNase-Seq on extracted chromatin and naked genomic DNA (gDNA). (**b**) Pie chart summarizes the sequence composition (capture/reporter) of FLASH^53^ reads. Promoter contacts (capture) with distal genomic elements (reporter) were enriched by two rounds of hybridization with promoter-specific probes (see Fig. 9b and **Supplementary Table 10**).

**Supplementary Figure 13.**
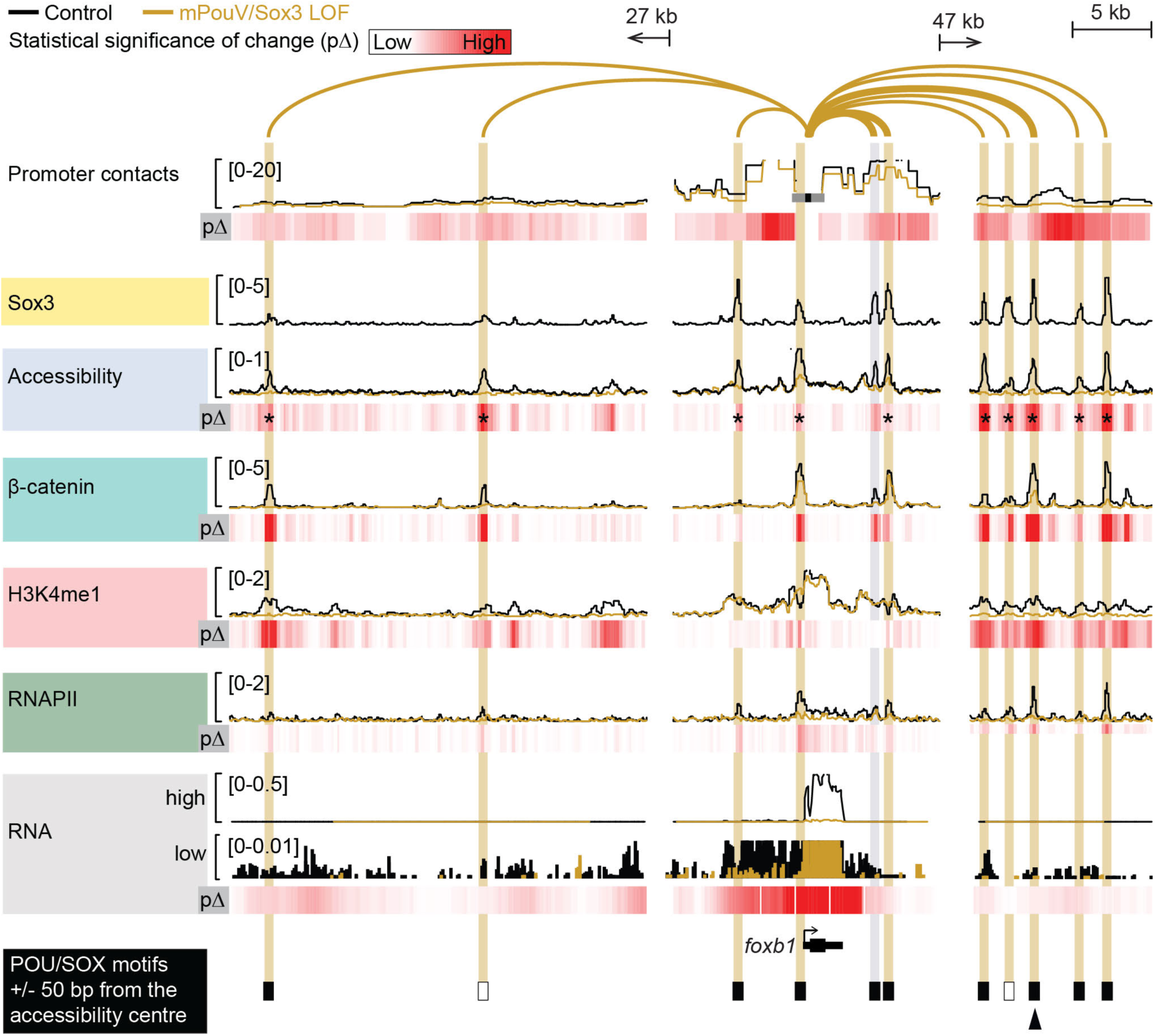
Pioneering activity of maternal PouV/Sox3 initiates extensive chromatin remodeling. Superimposed line tracks show the level of promoter-tied chromatin conformations, chromatin accessibility and various chromatin components (β-catenin, H3K4me1 and RNAPII) at the *foxb1* gene locus between control (uninjected) and mPouV/Sox3 LOF embryos. The RNA track is split into a high (0-0.5) and low (0-0.01) expression window. Note that the low-expression window shows that locally transcribed non-coding super-enhancer RNA depend on mPouV/Sox3 as well as the gene *foxb1*. Heat maps (pΔ) below each superimposed line plot show the statistical significance of changes caused by mPouV/Sox3 LOF. The footer highlights the occurrences of canonical POU/SOX motifs (black filled rectangles) at accessible pCRMs (±50 bp from the accessibility centre) and one strongly affected pCRM with an arrowhead. Asterisks on the pΔ heat map mark significant (FDR ≤10%) reductions to pCRM accessibility. pCRMs are boxed in and their frequency of contacts with the *foxb1* promoter are illustrated with an arc of varying strength. Boxes of affected pCRM and arcs of promoter contacts are colored orange.

**Supplementary Figure 14.**
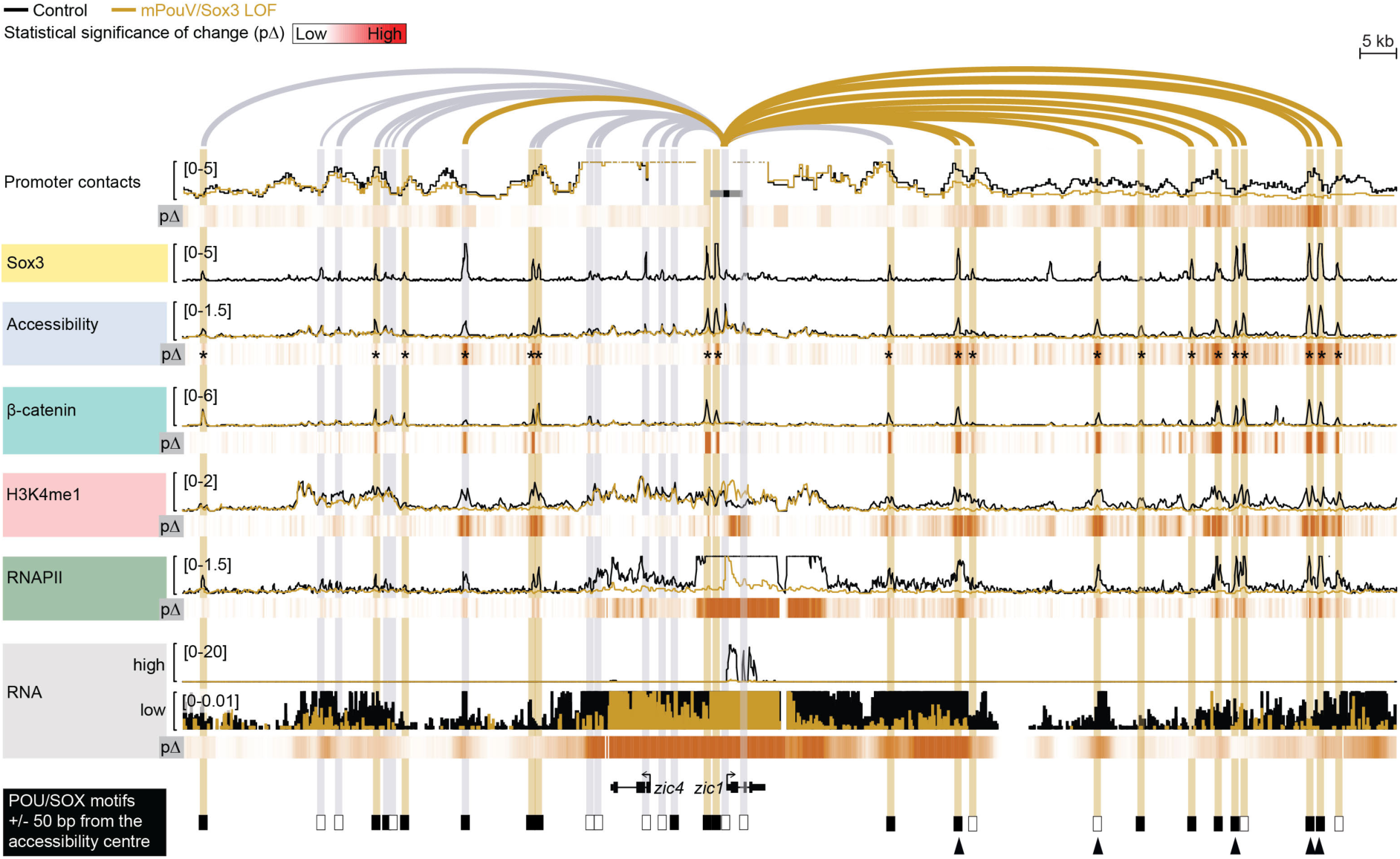
Pioneering activity of maternal PouV/Sox3 initiates extensive chromatin remodeling. Superimposed line tracks show the level of promoter-tied chromatin conformations, chromatin accessibility and various chromatin components (β-catenin, H3K4me1 and RNAPII) at the *zic1* gene locus between control (uninjected) and mPouV/Sox3 LOF embryos. The RNA track is split into a high (0-20) and low (0-0.01) expression window. Note that the low-expression window shows that locally transcribed non-coding super-enhancer RNA depend on mPouV/Sox3 as well as the gene *zic1*. Heat maps (pΔ) below each superimposed line plot show the statistical significance of changes caused by mPouV/Sox3 LOF. The footer highlights the occurrences of canonical POU/SOX motifs (black filled rectangles) at accessible pCRMs (±50 bp from the accessibility centre) and strongly affected pCRMs with arrowheads. Asterisks on the pΔ heat map mark significant (FDR ≤10%) reductions to pCRM accessibility. pCRMs are boxed in and their frequency of contacts with the *zic1* promoter are illustrated with an arc of varying strength. Boxes of affected pCRMs and arcs of promoter contacts are colored orange.

**Supplementary Figure 15.**
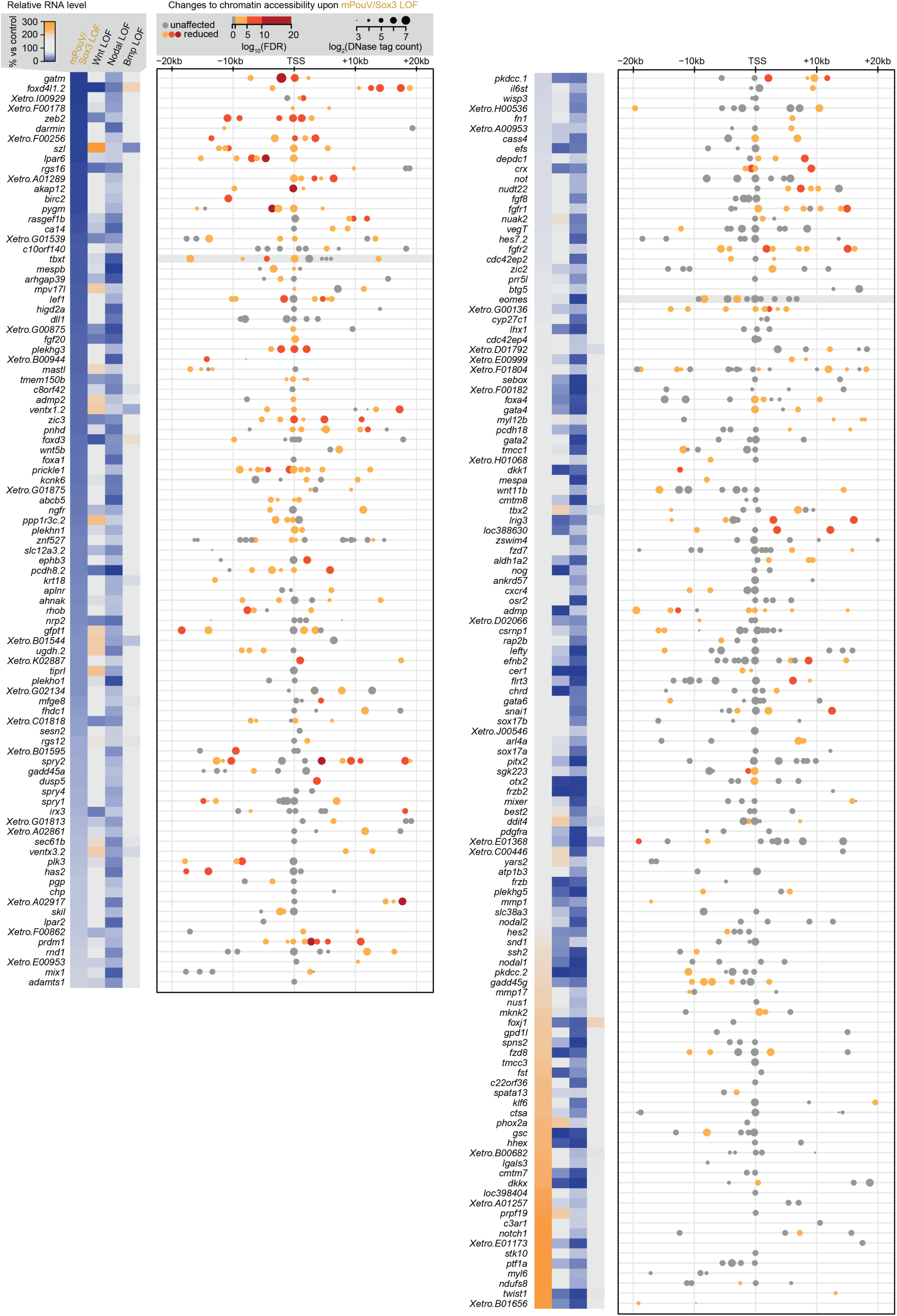
mPouV/Sox3-induced chromatin accessibility is required for the expression of Nodal-responsive genes. The heat map shows the transcript levels of Nodal-responsive genes under indicated LOFs. All genes listed here are active by the MBT^52^ and their transcript levels are significantly reduced (≥two-fold; FDR ≤10%) upon α-amanitin injection (see Fig. 6e). The plot aligned to the heat map shows the localization and DNase sensitivity (bubble size) of accessible pCRMs (affected, dot colored in orange to red with FDR decreasing from 10%; and unaffected, grey dot) relative to zygotic TSSs. Gene loci are sorted by mPouV/Sox3 LOF-induced transcript fold changes as shown in the heat map.

**Supplementary Figure 16.**
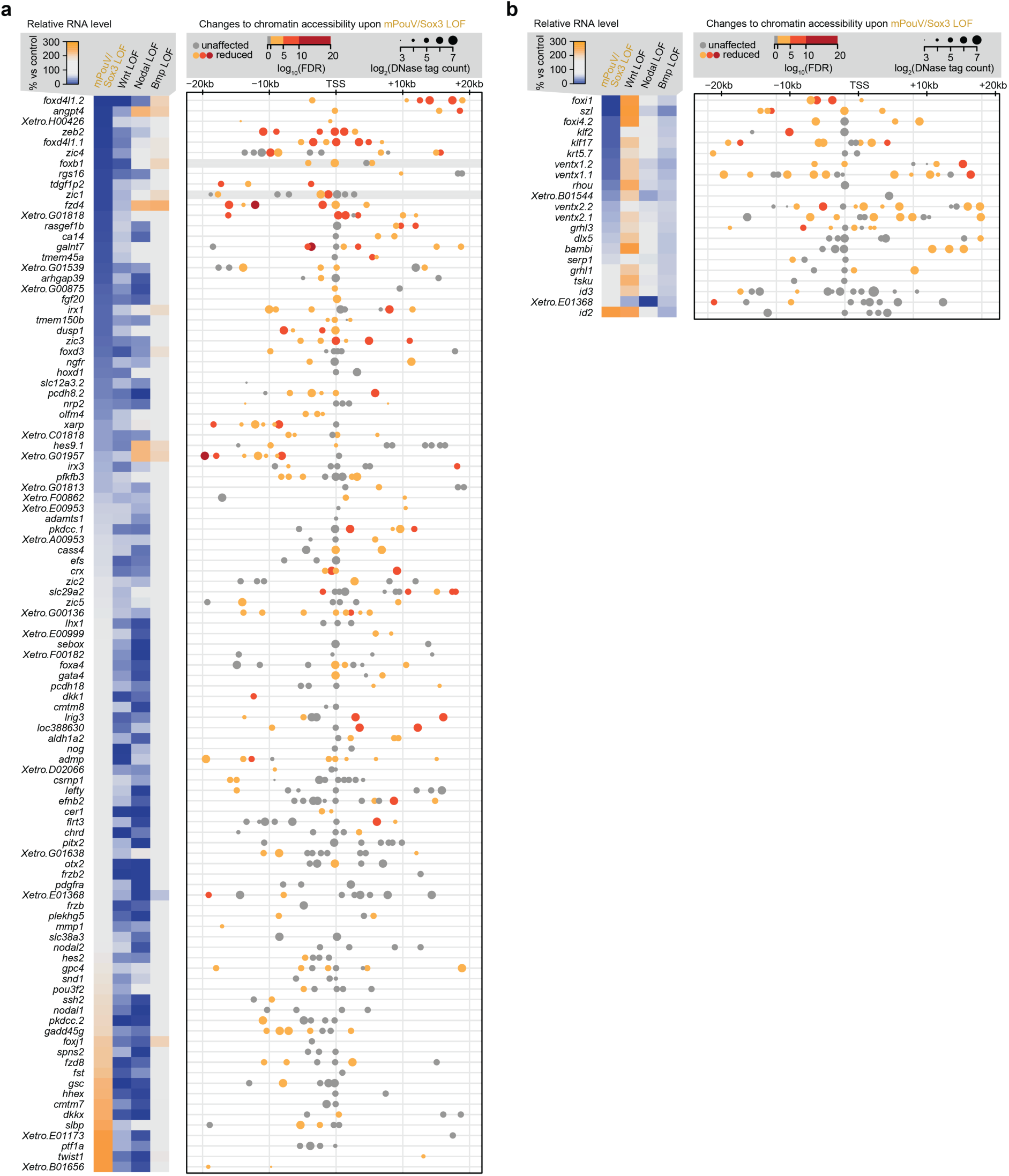
mPouV/Sox3-induced chromatin accessibility is required for the expression of Wnt- and BMP responsive genes. (**a**,**b**) Heat maps show the transcript levels of Wnt- or Bmp-responsive genes under indicated LOFs. All genes listed here are active by the MBT^52^ and their transcript levels are significantly reduced (≥two-fold; FDR ≤10%) upon α-amanitin injection (see Fig. 6e). The plot aligned to the heat map shows the localization and DNase sensitivity (bubble size) of accessible pCRMs (affected, dot colored in orange to red with FDR decreasing from 10%; and unaffected, grey dot) relative to zygotic TSSs. Gene loci are sorted by mPouV/Sox3 LOF-induced transcript fold changes as shown in the heat map.

**Supplementary Figure 17.**
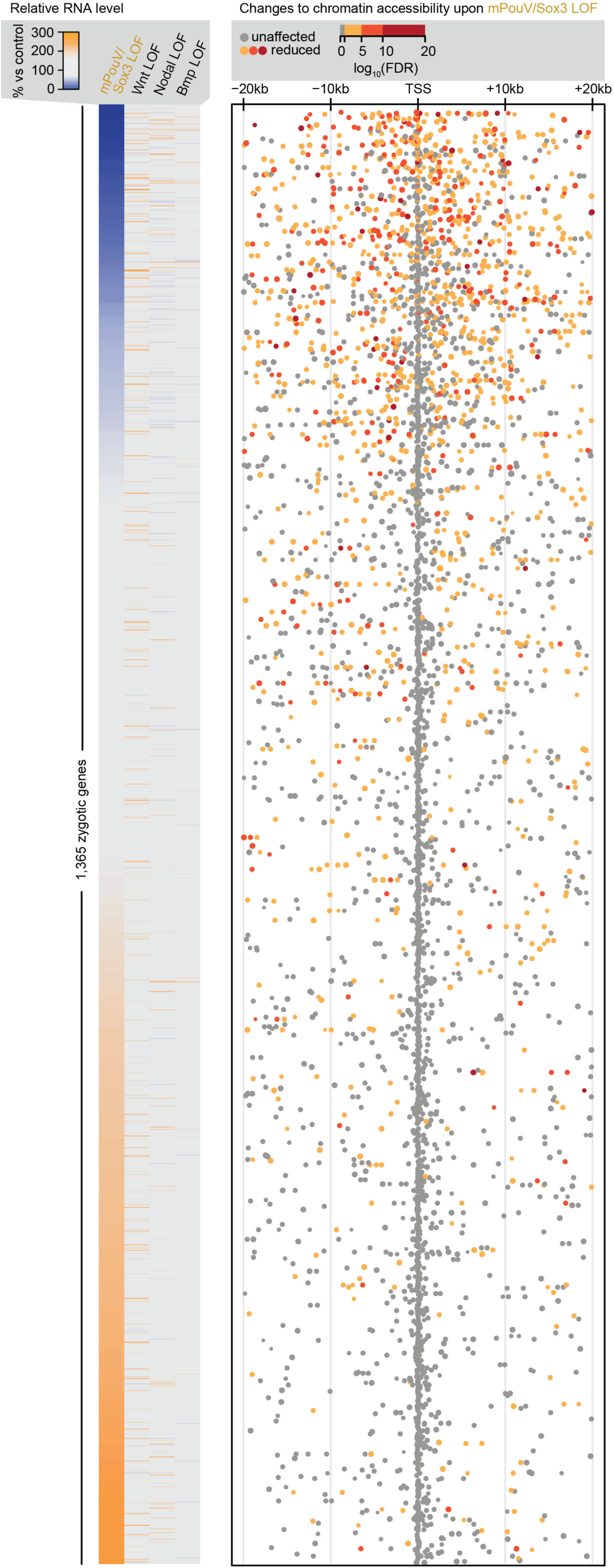
mPouV/Sox3-induced chromatin accessibility is required for the expression of signal non-responsive genes. The heat map shows the transcript levels of 1,365 signal non-responsive genes under indicated LOFs. The plot aligned to the heat map shows the localization of accessible pCRMs (affected, dot colored in orange to red with FDR decreasing from 10%; and unaffected, grey dot) relative to the zygotic TSSs. All genes listed here are active by the MBT^52^ and their transcript levels are significantly reduced (≥two-fold; FDR ≤10%) upon α-amanitin injection (see Fig. 6e). Gene loci are sorted by mPouV/Sox3 LOF-induced transcript fold changes as shown in the heat map.

**Supplementary Movie 1** Gastrulation defects caused by the loss-of-function of maternal pluripotency factors.

**Supplementary Table 1** Summary of deep sequencing and read alignments.

**Supplementary Table 2** Annotation of RNAPII^+^ pCRMs (genomic coordinates, distance to nearest TSS, DNA occupancy levels from the 32-cell to the late gastrula stage, DNA motif counts)

**Supplementary Table 3** Annotation of accessible pCRMs (genomic coordinates, distance to nearest TSS, normalized DNase cleavage count and DNA occupancy levels of RNAPII and H3K4me1)

**Supplementary Table 4** Maternal protein concentrations versus ribosome densities (translation) across the mid-blastula transition.

**Supplementary Table 5** ChIP-Seq peak calling coordinates for TFs (Eomes, Foxh1, Sox3, Tbx6, Tbxt and VegT) and signal mediators (catenin, Smad1 and Smad2) at indicated developmental stages.

**Supplementary Table 6** Genomic coordinates of super-enhancers assembled from individual TF and signal mediator ChIP-Seq profiles separated by developmental stages (up to MBT, early gastrula stage and late gastrula stage)

**Supplementary Table 7** ChIP-Seq peak calling coordinates for Sox3 and MyoD-HA in early gastrula embryos with ectopic expression of MyoD-HA.

**Supplementary Table 8** ChIP-Seq peak calling coordinates for Sox3 in dissected parts of the early tailbud embryo (head, trunk and bud).

**Supplementary Table 9** Transcriptional effect of maternal transcription factors and inductive signals on zygotic genome activation.

**Supplementary Table 10** Genomic coordinates of selected capture-C viewpoints and DNA sequences used for capturing promoter-tied regulatory elements.

**Supplementary Table 11** MBT-staged chromatin accessibility changes caused by mPouV/Sox3 LOF.

**Supplementary Table 12** Results of the next-generation capture-C performed at the MBT on control and mPouV/Sox3 LOF embryos.

## ONLINE METHODS

### *Xenopus* manipulation

Standard procedures were used for ovulation, fertilization, and manipulation and incubation of embryos^55, 56^. Briefly, frogs were obtained from Nasco (Wisconsin, USA). Ovulation was induced by injecting serum gonadotropin (Intervet) and chorionic gonadotropin (Intervet) into the dorsal lymph sac of mature female frogs. Eggs were fertilized *in vitro* with sperm solution consisting of 90% Leibovitz’s L-15 medium (Gibco) and 10% fetal calf serum (Gibco). After 10 min, fertilized eggs were de-jellied with 2.2% (w/v) L-cysteine (Sigma) equilibrated to pH 8.0. *X. tropicalis* embryos were cultured in 5% Marc’s Modified Ringer’s solution (MMR) ^56^ at 21°C-28°C. *X. laevis* embryos were cultured in 10% Normal Amphibian Medium (NAM)^56^ at 14°C-22°C. Embryos were staged as previously described^57^. All *Xenopus* work fully complied with the UK Animals (Scientific Procedures) Act 1986 as implemented by the Francis Crick Institute.

### Nucleic acid injections and treatments of embryos

Microinjections were performed using calibrated needles and embryos equilibrated in 4% (w/v) Ficoll PM-400 (Sigma) in 5% MMR or 10% NAM. Microinjection needles were generated from borosilicate glass capillaries (Harvard Apparatus, GC120-15) using the micropipette puller Sutter p97. Maximally two nanolitres of morpholino (MO) and/or mRNA were injected into de-jellied embryos at the 1-cell, 2-cell or 4-cell stage using the microinjector Narishige IM-300. Injections for the mVegT loss-of-function (LOF) and rescue were targeted to the vegetal hemisphere. All other injections were targeted to the animal hemisphere. Embryos were transferred to fresh 5% MMR or 10% NAM (without Ficoll) once they reached about blastula stage.

The following mRNA amounts were used for ectopic protein expression: 80 pg and 400 pg *MyoD*-*HA* mRNA into *X. tropicalis* and *X. laevis* embryos, respectively; and 400 pg *Sox3*-*HA* into *X. laevis* embryos.

The following MO amounts and treatments were used for LOF experiments: 5 ng *Pou5f3.2* MO, 5 ng *Pou5f3.3* MO and 5 ng standard control MO (mPouV LOF); 5 ng *Pou5f3.2* MO, 5 ng *Pou5f3.3* MO and 5 ng *Sox3* MO (mPouV/Sox3 LOF); 5 ng *β-catenin* MO (Wnt LOF); 10 ng m*VegT* MO (mVegT LOF); standard control MO (5-10 ng according to the dose used for the β-catenin or mVegT LOF experiment); and 30 pg α-amanitin (BioChemica, #A14850001). To block Nodal (Nodal LOF) or BMP (BMP LOF) signaling, embryos were treated with 100 µM SB431542 (Tocris, #1614) or 10 µM LDN193189 (Selleckchem, #S2618) from the 8-cell stage onwards. Control embryos were treated accordingly with DMSO, in which these antagonists were dissolved. MOs were designed and produced by Genetools (Oregon, USA) to block translation. β-catenin MO (5’-TTTCAAC<u>A</u>GTTTCCAAAGAACCAGG-3’, underlined base was changed from the original MO^26^), Pou5f3.2 MO (5’-GCTGTTGGCTGTACAT<u>A</u>GTGTC-3’, underlined base was changed from the original MO^25^), Pou5f3.3 MO^25^ (5’-GTACAGAACGGGTTGGTCCATGTTC-3’), Sox3 MO (5’-GTCTGTGTCCAACATGCTATACATC-3’), Tbx6 MO (5’-TACATTGGGTGCAGGGACCCTCTCA-3’) and mVegT MO^58^ (5’-TGTGTTCCTGACAGCAGTTTCTCAT-3’; see Supplementary Fig. 2e).

For the rescue of mVegT LOF, 25-50 pg z*VegT* mRNA were injected at the 2- or 4-cell stage. For the rescue of mPouV/Sox3 LOF, 75 pg *Pou5f3.3* mRNA and 75 pg *Sox2* mRNA were injected at the 2- or 4-cell stage. For both rescue experiment, 150-300 pg *mCherry* mRNA was co-injected to trace mRNA expression in embryonic cells.

### RNA constructs

All constructs are products of Gateway (Invitrogen) cloning. First, the coding sequences of interest were amplified from either plasmid using *Pfu*Turbo DNA polymerase (Agilent, #600250) or cDNA using Phusion HF DNA polymerase (NEB, #M0530), and unidirectionally cloned into the pENTR/D-TOPO vector (Thermo Fisher Scientific, #K2400). Next, these clones were LR-recombined with Gateway-compatible pCS2+ vectors containing no tag or a C-terminal 3xHA tag (see Fig. 4g). The following forward and reverse primers were used for PCR amplification: *X. laevis myoD1a* from *MyoD*2-24 pBluescript plasmid^59^, 5’-CACCATGGAGCTGTTGCCCCCAC-3’, 5’-TAAGACGTGATAGATGGTGCTG-3’; *X. laevis sox3* from the IMAGE:443920 clone (European *Xenopus* Resource Centre), 5’-CACCATGTATAGCATGTTGGACAC-3’, 5’-TATGTGAGTGAGCGGTACAGTG-3’; *X. laevis sox2* from stage 11 cDNA, 5’-CACCATGTACAGCATGATGGAGACCGA-3’, 5’-TCACATGTGCGACAGAGGCAGC-3’; *X. laevis pou5f3.3* from stage 11 cDNA, 5’-CACCATGGACCAGCCCATATTGTACA-3’, 5’-TCAGCCGGTCAGGACCCCCA-3’; *X. tropicalis* z*VegT* from stage 11 cDNA, 5’-CACCATGCACTCTCTGCCGGATGTA-3’, 5’-TTACCAACAGCTGTATGGAAAGA-3’. pCS2+8N*mCherry*^60^ (#34936) was obtained from Addgene.

### *In vitro* transcription of capped mRNA

About 10 µg plasmid was linearized by restriction digestion and purified using the QIAquick PCR purification kit (Qiagen, #28104). Capped mRNA was produced from ∼1 µg linearized plasmid using mMessage mMachine SP6 (Thermo Fisher Scientific, #AM1340) according to the manufacturer’s instructions. The following restrictions enzymes (NEB) were used for linearizing the pCS2+ constructs: *Asp*781 (*myoD1a*, *sox3*), *Pvu*II (*sox2*), *Psp*OMI (*pou5f3.3*, z*VegT*) and *Not*I (*mCherry*).

### Generation of digoxigenin-labeled RNA probes

DNA template for generating the *foxb1 in situ* hybridization probe was amplified from *X. tropicalis* stage 19-20 cDNA. KAPA HiFi HotStart polymerase (Kapa Biosystems, #KK2602) and the following PCR cycling conditions were used: 98°C for 45 sec followed by 40 cycles (98°C for 10 sec, 63°C for 10 sec, 72°C for 15 sec) and final elongation step of 20 sec at 72°C. Primers were 5’-GTCAGCGCCTATGGAGTACC-3’ and 5’-AACACTGGAGATGCCATGC-3’. Fresh PCR product was zero-blunt cloned into the pCRII-TOPO vector (Thermo Fisher Scientific, #K2800). The identity and direction of insert was verified by restriction digest and Sanger sequencing. Plasmids were linearized by restriction digestion and purified using the QIAquick PCR purification kit (Qiagen, #28104). All *in situ* hybridization probes were transcribed from ∼1 µg linearized plasmid using 1x digoxigenin-11-UTP (Roche, #11277065910), 40 U RiboLock RNase inhibitor (Thermo Fisher Scientific, #EO0381), 1x transcription buffer (Roche) and T7 RNA polymerase (Roche, #10881767001) at 37°C for 2 h. The probe was treated with 2 U Turbo DNase (Thermo Fisher Scientific, #AM2238) to remove the DNA template and purified by spin-column chromatography (Clontech). RNA was diluted to 10 ng/µl (10x stock) with hybridization buffer (50% formamide, 5x SSC, 1x Denhardt’s, 10 mM EDTA, 1 mg/ml torula RNA, 100 µg/ml heparin, 0.1% Tween-20 and 0.1% CHAPS) and stored at −20°C. The following plasmids and restriction enzymes (NEB) were used for plasmid linearization: *X. laevis eomes* pCRII-TOPO^19^, *Bam*HI; *X. tropicalis foxb1* pCRII-TOPO, *Bam*HI; *X. tropicalis mir427* pCS108 (IMAGE:7545411)^50^, *Sal*I; *X. laevis ventx2.1* pBluescript SK-^61^, *Eco*RI; *X. laevis tbxt (Xbra)* pSP73^62^, *Bgl*II; and *X. tropicalis zic1* pCMV-SPORT6 (IMAGE:7668846), *Sal*I.

### Animal cap assays

These assays were carried out for the ectopic expression of mRNA constructs in *X. laevis* and for the MO-mediated LOF in *X. tropicalis*. All animal caps were dissected at the blastula stage (stage 8 to 9) and cultured in 75% NAM containing 0.1% bovine serum albumin (Sigma). Dissections were carried out with 13 µm loop electrodes (Protech International, 13-Y1) connected to a MC-2010 micro cautery instrument (Protech International) operating at power level 2. As illustrated in Fig. 4g, control (uninjected) and MyoD-HA or Sox3-HA expressing animal caps were cultured at 20°C until sibling embryos reached early neurula stage (stage 13). For the experiment shown in Fig. 7d and Supplementary Fig. 10d, animal caps of control and mPouV/Sox3 LOF embryos were cultured with or without 10 ng/ml recombinant human activin A (Nodal agonist; R&D Systems, #338-AC) or 50 µM CHIR99021 (canonical Wnt agonist; Tocris, #4423) at 22°C for ∼2 h until sibling embryos reached early gastrula stage (stage 10^+^).

### Extraction of total RNA

10-15 embryos (or 15-20 animal caps) were homogenized in 800 (400) µl TRIzol reagent (Thermo Fisher Scientific, #15596018) by vortexing. The homogenate was either snap-frozen in liquid nitrogen and stored at −80°C or processed immediately. For phase separation, the homogenate together with 0.2x volume of chloroform was transferred to pre-spun 1.5-ml Phase Lock Gel Heavy microcentrifuge tubes (VWR), shaken vigorously for 15 sec, left on the bench for 2 min and spun at ∼16,000 g (4°C) for 5 min. The upper phase was mixed well with one volume of 95-100% ethanol and spun through the columns of Zymo RNA Clean and Concentrator 5 or 25 (Zymo Research) at ∼12,000 g for 30 sec. Next, the manufacturer’s instructions were followed for the recovery of total RNA (>17 nt) with minor modifications. First, the flow-through of the first spin was re-applied to the column. Second, the RNA was treated in-column with 3 U Turbo DNase (Thermo Fisher Scientific, #AM2238). Third, the RNA was eluted twice with 25 µl molecular-grade water. The concentration was determined on the NanoDrop 1000 spectrophotometer.

### Reverse transcription (RT)

About 0.2-1 µg total RNA was denatured at 75°C for 5 min before setting up the RT reaction including 40 U M-MLV (RNase H minus, point mutant) RT (Promega, #M3681), 500 µM of each dNTP (NEB), and 10 µM random hexamers (Sigma) in a final volume of 10 µl. The following incubation settings were used: 25°C for 15 min, 37°C for 15 min, 55°C for 45 min, 85°C for 15 min and 4°C for indefinite. The reaction was diluted with 90-190 µl molecular-grade water before using 1-2.5 µl for each qPCR reaction.

### qPCR primers

The PCR primers were designed to hybridize at ∼60°C (T_m_) and to generate 75-125 bp amplicons. Where possible, primers were checked for their specificity *in silico* using Jim Kent’s PCR tool (http://genomes.crick.ac.uk/cgi-bin/hgPcr). All primer pairs were tested to produce one amplicon of the correct size through diagnostic PCR using RedTaq DNA polymerase (Sigma, #D4309). Exceptions were previously designed primer pairs for *X. laevis actc1*, *odc1* and *otx2*, all of which generate amplicons of 200-250 bp.

Primers (forward, reverse) for *X. laevis* (L) or *X. tropicalis* (T) cDNA: *actc1* (L, 5’-TCCTGTACGCTTCTGGTCGTA-3’, 5’- TCTCAAAGTCCAAAGCCACATA-3’); *eomes* (T, 5’-ATTGACCACAACCCATTTGC-3’, 5’- TCTAGGAGAATCCGCAGGAG-3’); *foxb1* (T, 5’-CCGGATGCAGAGTGACAAC-3’, 5’-GAGGCTTCTGGTCGCTGTAG-3’); *gata4* (T, 5’-TCCTCTGATCAAGCCTCAAAG-3’, 5’-ACGGCGCCAGAGTGTAGTAG-3’); *myl1* (L, 5’- GCACCTGCATTTGATCTGTC-3’, 5’-TTGCTGTCTCCTGTCCTGTC-3’); *nodal5* (T, 5’-TTCAAGCCAACAAACCACGC-3’, 5’-CGCATCTTCACCGGTATGCA-3’); *nodal6* (T, 5’-CGCATTTCGTTGTGAGGGGT-3’, 5’-TCGTACAGCTTGACCAAACTCT-3’); *odc1* (L, 5’- GCCATTGTGAAGACTCTCTCC-3’, 5’- TTCGGGTGATTCCTTGCCAC-3’); *odc1* (T, 5’-GGGCAAAAGAGCTTAATGTGG-3’, 5’-CATCGTGCATCTGAGACAGC-3’); *otx2* (L, 5’-TTGAACCAGACCTGGACT-3’, 5’-CGGGATGGATTTGTTGCA-3’); *rpl8* (T, 5’- CTGGCTCCAAGAAGGTCATC-3’, 5’-CAGGATGGGTTTGTCAATACG-3’); *sox2* (L, 5’- ACATCACATGGACGGTTGTC-3’, 5’-TCTCTCGGTTTCTGCTCTCG-3’); *sox17b* (T, 5’- GTTTATGGTGTGGGCAAAGG-3’, 5’- TCAGCGATTTCCATGATTTG-3’); *tbx6* (L, 5’-GGGGCACCACGACTCTTAC-3’, 5’-CCCAGAATCCTCTGTGAAGG-3’); and *tbxt* (T, 5’-CCTGTGGATGAGGTTCAAGG-3’, 5’-CACGCTCACCTTTAGAACTGG-3’).

Primers (forward, reverse) for *X. tropicalis* genomic DNA: *β-actin* −0.5kb (5’-TTGTACGCAACACAATGCAG-3’, 5’-AAATGTCAGGACCAAATGCAG-3’); *dlc* −3.2kb (5’-CATCCTGCAATTAACTGTCTGC-3’, 5’-AGGTGTGACAAATGGCACTG-3’); *gdf3* −0.1kb (5’-TGGATGAGCACAGAGAGGTG-3’, 5’-GTCAGGAGGGAGGTGTCAAC-3’); *hoxd1* +3.2kb (5’-TGTTGTAGATGCTGATGCTTATCG-3’, 5’-AACAGAAAATCAAAGGCTTGCA-3’); *mespa* −1.3kb (5’-CTCCTTTGCCCTCTGAAATG-3’, 5’-CTTCAGGGAATTGGCTTGTG-3’); *msgn1* −0.1kb (5’-CCAGTCCATTTTCCATGTTG-3’, 5’-GGGGAGGCCCTTTTATACAG-3’); *myf5* −1.1kb (5’-AAAGAGTTCCTGCACCTTGG-3’, 5’-TCCCAAATCATCAGCACAAC-3’); *nodal6* −0.1kb (5’-ACATGAGTTTGGCCTTGGTC-3’, 5’-AGGTAACCCCTGAGCTGTCA-3’); *pax3* 5’UTR (5’-CAGATGCCAGAGGAGAGAGC-3’, 5’-GGAACTCTCTGGGTCCCAAC-3’); *pax3* +2.7kb (5’-ATGAGAATGTGAGCGACACG-3’, 5’-AAACCCAAGGTGAGCAAGTG-3’); *sia1* −0.2kb (5’-TGGCTTTTAAGTTCGTCTTGG-3’, 5’-CAAAGGGCTTCTCTTTGTGC-3’); *sox3* TSS (5’-GATGGGGCATTAATGGACAG-3’, 5’-CACGGGAGAAAATGAAGCTG-3’); *sox17b* −0.1kb (5’-GTTAGACTCCGCTCCAGACG-3’, 5’-TCTCCACCTAAGGGGAAACC-3’); *wnt8a* −5.7kb (5’-GGAAACTGCCTGTGGAAATG-3’, 5’-ACAAAAGCCCACAGGAGATG-3’); *tbxt* −4.0kb (5’-CAATGGATGCCTCTCTGGAC-3’, 5’-TGGGTTTAGAATGGCAAAGG-3’); and *tbxt* TSS (5’-CACACCTTGGGTTTTGTTCC-3’, 5’-TTCCTCAAGAGAGGGGCTTC-3’).

### Whole-mount *in situ* hybridization (WMISH)

WMISH was conducted using digoxigenin-labeled RNA probes. It was based on previously published protocols^56, 63^. *X. tropicalis* embryos were fixed in MEMFA (1x MEM and 3.7% formaldehyde) at room temperature for 1 h. The embryos were then washed once in 1x PBS and two to three times in ethanol. Fixed and dehydrated embryos were kept at −20°C for ≥24 h to ensure proper dehydration before starting hybridization. Dehydrated embryos were washed once more in ethanol before rehydrating them in two steps to PBT (1x PBS and 0.1% Tween-20). Embryos were treated with 5 µg/ml proteinase K (Thermo Fisher Scientific, #AM2548) in PBT for 6-8 min, washed briefly in PBT, fixed again in MEMFA for 20 min and washed three times in PBT. Embryos were transferred into baskets, which were kept in an 8×8 microcentrifuge tube holder sitting inside a 10×10 slot plastic box filled with PBT. Baskets were built by replacing the round bottom of 2-ml microcentrifuge tubes with a Sefar Nitex mesh. This container system was used to readily process several batches of embryos at once. These baskets were maximally loaded with 40 to 50 *X. tropicalis* embryos. The microcentrifuge tube holder was used to transfer all baskets at once and to submerge embryos into subsequent buffers of the WMISH protocol. Next, the embryos were incubated in 500 µl hybridization buffer (50% formamide, 5x SSC, 1x Denhardt’s, 10 mM EDTA, 1 mg/ml torula RNA, 100 µg/ml heparin, 0.1% Tween-20 and 0.1% CHAPS) for 2 h in a hybridization oven set to 60°C. After this pre-hybridization step, the embryos were transferred into 500 µl digoxigenin-labeled probe (1 ng/µl) preheated to 60°C and further incubated overnight at 60°C. The pre-hybridization buffer was kept at 60°C. The next day embryos were transferred back into the pre-hybridization buffer and incubated at 60°C for 10 min. Subsequently, they were washed three times in 2x SSC/0.1% Tween −20 at 60°C for 10 min, twice in 0.2x SSC/0.1% Tween-20 at 60°C for 20 min and twice in 1x maleic acid buffer (MAB) at room temperature for 5 min. Next, the embryos were treated with blocking solution (2% Boehringer Mannheim blocking reagent in 1x MAB) at room temperature for 30 min, and incubated in antibody solution (10% lamb or goat serum, 2% Boehringer Mannheim blocking reagent, 1x MAB and 1:2,000 Fab fragments from polyclonal anti-digoxigenin antibodies conjugated to alkaline phosphatase) at room temperature for 4 h. The embryos were then washed four times in 1x MAB for 10 min before leaving them in 1x MAB overnight at 4°C. On the final day of the WMISH protocol, the embryos were washed another three times in 1x MAB for 20 min and equilibrated to working conditions of alkaline phosphatase (AP) for a total of 10 min by submerging embryos twice into AP buffer (50 mM MgCl_2_, 100 mM NaCl, 100 mM Tris-HCl pH 9.5 and 1% Tween-20). At this stage, the embryos were transferred to 5-ml glass vials for monitoring the progression of the AP-catalyzed colorimetric reaction. Any residual AP buffer was discarded before adding 700 µl staining solution (AP buffer, 340 µg/ml nitro-blue tetrazolium chloride, 175 µg/ml 5-bromo-4-chloro-3’-indolyphosphate). The colorimetric reaction was developed at room temperature in the dark. Once the staining was clear and intense enough, the color reaction was stopped by two washes in 1x MAB. To stabilize and preserve morphological features, the embryos were fixed with Bouin’s fixative without picric acid (9% formaldehyde and 5% glacial acetic acid) at room temperature for 30 min. Next, the embryos were washed twice in 70% ethanol/PBT to remove the fixative and residual chromogens. After rehydration to PBT in two steps, the embryos were treated with weak Curis solution (1% H_2_O_2_, 0.5x SSC and 5% formamide) at 4°C in the dark overnight. Finally, the embryos were washed twice in PBS before imaging them in PBS on a thick agarose dish by light microscopy.

### Antibody generation

Affinity-purified rabbit polyclonal antibodies were generated by sdix (Delaware, USA) against different amino acid (aa) sequences of *X. tropicalis* Tbx6 avoiding its highly conserved DNA binding domain (aa 95-280) among T-box proteins: aa 1-94 (#4596), aa 304-403 (#5061) and aa 418-517 (#5107). Antibody #5107 did not work (data not shown). Antibodies #4596 and #5061 were suitable for immunoprecipitation and Western blotting to detect endogenous Tbx6 in both *X. tropicalis* and *X. laevis* embryos (Supplementary Fig. 2k-m). Antibody #4596 complied with ENCODE guidelines^64^ and thus was suitable for ChIP (**Supplementary Fig. 2n**).

### Immunoprecipitation (IP)

For each IP, 40 de-jellied embryos were collected at the developmental stage of interest. They were homogenized in 400 µl ice-cold IP buffer (50 mM Tris-HCl pH 7.5, 150 mM NaCl, 5 mM MgCl_2_, 1 mM EDTA and 0.25 % Igepal CA-630) supplemented with an EDTA-free protease inhibitor cocktail (Roche, #11873580001). The extract was kept on ice for 5 min. Subsequently, the extract was mixed with 400 µl 1,1,2-trichloro-1,2,2-trifluoroethane (Sigma) by vigorously inverting the tube and cleared by centrifugation at 16,000 g (4°C) for 5 min. An aliquot of 20 µl supernatant was saved as input. The remaining supernatant was incubated with 0.5 µl antibody (anti-Sox3^48^, anti-VegT^15^ or anti-Tbx6; all rabbit polyclonal antibodies) at 4°C for 1 h. Subsequently, each embryonic extract was supplemented with 20 µl washed magnetic beads conjugated to sheep anti-rabbit IgG (Dynabeads M280, Invitrogen) and kept on a tube rotator at 4°C for another 2 h. Next, the beads were washed three times with IP buffer. The immunoprecipitate was eluted off the beads with 20 µl 1x SDS loading buffer (50 mM Tris-HCl pH 6.8, 2% SDS, 6% glycerol and 5% β-mercaptoethanol). Western blotting was carried out with TrueBlot (Rockland Immunochemicals) and normal horseradish peroxidase (HRP)-conjugated secondary antibodies for IP and input samples, respectively.

### Western blotting

Protein samples were denatured in 1x (final concentration) SDS loading buffer at 70°C for 5 min. Denatured samples were run alongside a standard protein ladder into pre-cast gradient gels (Any kD Mini-PROTEAN TGX, BioRad) at constant 200 V for 1 h. Size-separated proteins were immediately transferred onto hydrophobic Immobilon-P PVDF transfer membranes (Millipore) at constant 100 V for 30 min using standard protein electrophoresis equipment. The membranes were blocked with 5% milk powder in PBS/0.1% Tween-20 (MPBTw) at room temperature for 30 min. Next, the membranes were incubated at room temperature for 1 h with the primary antibodies diluted in MPBTw: 1:5,000 of mouse monoclonal anti-α-tubulin (Sigma, #T5168), 1:2,000 rabbit polyclonal anti-Sox3^48^, 1:2,000 rabbit polyclonal anti-VegT^15^ or 1:2,000 rabbit polyclonal anti-Tbx6 (#5061). The membranes were washed three times with PBST for 10 min before applying 1:1,000 TrueBlot HRP-conjugated anti-rabbit IgG (Rockland Immunochemicals, #18-8816-31) or 1:2,000 normal goat HRP-conjugated anti-mouse IgG (Thermo Fisher Scientific, #31430) in MPBTw at room temperature. After 1 h, the membranes were washed three times in PBTw for 10 min and once in PBS for 5 min. Finally, the membranes were treated with UptiLight US HRP WB reagent (interchim, #58372) for 1 min in the dark. HRP-catalyzed chemiluminescence was detected on the ChemiDoc XRS+ system (BioRad).

### Whole-mount immunohistochemistry (WMIHC)

Embryos were fixed with MEMFA (0.1 M MOPS, 2 mM EGTA, 1 mM MgSO_4_ and 3.7% formaldehyde) in capped 5-ml glass vials at room temperature for 2 h, dehydrated and stored in absolute ethanol at −20°C for ≥24 h. For sagittal sections, fixation was stopped after 30 min. Embryos were placed on a piece of flattened Blu-Tack (Bostik) in a large Petri dish filled with PBS to bisect them with a scalpel. Following a three-step rehydration (50%, 75% and 100% PBS) the embryos were bleached with weak Curis solution (1% H_2_O_2_, 0.5x SSC and 5% formamide) on a light box at room temperature for 2 h or in the dark at 4°C overnight. Bleached embryos were washed three times with PBS/0.3% Triton X-100 (PBT) and pre-incubated in blocking solution (PBS, 20% goat or donkey serum, 2% BSA and 0.1% Triton X-100) at 4°C for 6 h. The embryos were then transferred to 2-ml round-bottom microcentrifuge tubes before discarding all remaining blocking solution. The embryos were incubated at 4°C for 1-3 days in 50 µl blocking solution containing the primary antibody at the following dilutions: 1:1,000 rabbit polyclonal anti-Sox3^48^, 1:500 rabbit polyclonal anti phospho-Smad1/5/8 (Cell Signaling, #9511) or 1:500 goat polyclonal Smad2/3 (R&D Systems, #AF3797). Afterwards, the embryos were transferred back to capped 5-ml glass vials, washed three times with RIPA buffer (50 mM HEPES pH 7.5, 0.5 M LiCl, 1 mM EDTA, 1% Igepal CA-630 and 0.7% sodium deoxycholate) for 1 h and once with PBT for 5 min. Next, the embryos were incubated with the secondary HRP-conjugated antibody diluted in blocking solution at 4°C overnight: 1:400 goat anti-rabbit IgG-HRP (Thermo Fisher Scientific, #G-21234) or 1:200 donkey anti-goat IgG-HRP (Santa Cruz Biotechnology, #sc-2020). The washes of the previous day were repeated before pre-incubating the embryos in 400 µl inactive 3,3’-diaminobenzidine tetrahydrochloride with cobalt (Sigma, #D0426). After 1-2 min, the inactive solution was replaced with the H_2_O_2_-activated DAB solution. The HRP reaction was stopped after 40 sec by washing the embryos several times with PBT. In Figs. 2c and 5a,d, embryos were dehydrated in three steps to absolute methanol and cleared with Murray’s clear (2:1 benzyl benzoate/benzyl alcohol) on a glass depression slide.

### Deep sequencing and quality filter

All deep sequencing libraries were quality controlled: The DNA yield and fragment size distribution were determined by fluorometry and chip-based capillary or polyacrylamide gel electrophoresis, respectively. Libraries were sequenced on the Illumina platforms GAIIx or HiSeq by the Advanced Sequencing Facility of the Francis Crick Institute to produce single or paired-end reads of at least 40 bases. Next-generation capture-C libraries were sequenced on MiSeq with a read length of 150 bases. Sequencing samples and read alignment results are summarized in **Supplementary Table 1**. The metrics of paired-end alignments such as insert size mean and standard deviation were determined by Picard (CollectInsertSizeMetrics) from the Broad Institute (USA).

### Chromatin immunoprecipitation (ChIP)

ChIP was carried out as detailed previously^65^. Briefly, de-jellied *X. tropicalis* embryos were treated with 1% formaldehyde (Sigma, #F8775) in 1% MMR at room temperature for 15-45 min to cross-link chromatin proteins to nearby genomic DNA. Duration of fixation was determined empirically and depended mainly on the developmental stage and antibody epitopes^65^. Fixation was terminated by rinsing embryos three times with ice-cold 1% MMR. Where required, post-fixation embryos were dissected to select specific anatomical regions in ice-cold 1% MMR. Fixed embryos were homogenized in CEWB1 (10 mM Tris-HCl pH 8.0, 150 mM NaCl, 1 mM EDTA, 1% Igepal CA-630, 0.25% sodium deoxycholate and 0.1% SDS) supplemented with 0.5 mM DTT, protease inhibitors and, if using phospho-specific antibodies, phosphatase blockers (0.5 mM orthovanadate and 2.5 mM NaF). To solubilize yolk platelets and separate them from the nuclei, the homogenate was left on ice for 5 min and then centrifuged at 1,000 g (4°C) for 5 min. Homogenization and centrifugation was repeated once before resuspending the nuclei containing pellet in 1-3 ml CEWB1. Nuclear chromatin was solubilized and fragmented by isothermal focused or microtip-mediated sonication. The solution of fragmented chromatin was cleared by centrifuging at 16,000 g (4°C) for 5 min. Where required, ∼1% of the cleared chromatin extract was set aside for the input sample (negative control). ChIP-grade antibodies were used to recognize specific chromatin features and to enrich these by coupling the antibody-chromatin complex to protein G magnetic beads (Thermo Fisher Scientific, #10003D) and extensive washing. These steps were carried out at 4°C. The beads were washed twice in CEWB1, twice in WB2 (10 mM Tris-HCl pH 8.0, 500 mM NaCl, 1 mM EDTA, 1% Igepal CA-630, 0.25% sodium deoxycholate and 0.1% SDS), twice in WB3 (10 mM Tris-HCl pH 8.0, 250 mM LiCl, 1 mM EDTA, 1% Igepal CA-630 and 1% sodium deoxycholate) and once in TEN (10 mM Tris-HCl pH 8.0, 150 mM NaCl and 1 mM EDTA). ChIP was eluted off the beads twice with 100 µl SDS elution buffer (50 mM Tris-HCl pH 8.0, 1 mM EDTA and 1% SDS) at 65°C. ChIP eluates were pooled before reversing DNA-protein cross-links. Input (filled up to 200 µl with SDS elution buffer) and ChIP samples were supplemented with 10 µl 5 M NaCl and incubated at 65°C for 6-16 h. Samples were treated with proteinase K (Thermo Fisher Scientific, #AM2548) and RNase A (Thermo Fisher Scientific, #12091021) to remove any proteins and RNA from the co-immunoprecipitated DNA fragments. The DNA was purified with phenol:chloroform:isoamyl alcohol (25:24:1, pH 7.9) using 1.5-ml Phase Lock Gel Heavy microcentrifuge tubes (VWR) for phase separation and precipitated with 1/70 volume of 5 M NaCl, 2 volumes of absolute ethanol and 15 µg GlycoBlue (Thermo Fisher Scientific, #AM9516). After centrifugation, the DNA pellet was air-dried and dissolved in 11 µl elution buffer (10 mM Tris-HCl pH 8.5). The DNA concentration was determined on a fluorometer using high-sensitivity reagents for double-stranded DNA (10 pg/µl to 100 ng/µl).

The following antibodies were used for ChIP: rabbit polyclonal anti-β-catenin (Santa Cruz Biotechnology, #H-102), rabbit polyclonal anti-H3K4me1 (Abcam, #ab8895), rabbit polyclonal anti-HA (Abcam, #ab9110), mouse monoclonal anti-RNAPII (Covance, #MMS-126R), rabbit polyclonal anti-phospho-Smad1/5/8 (Cell Signaling, #9511), goat polyclonal anti-Smad2/3 (R&D Systems, #AF3797), rabbit polyclonal anti-Sox3^48^, rabbit polyclonal anti-Tbx6 (#4596) and rabbit polyclonal anti-VegT^15^.

### Quantitative PCR (qPCR)

DNA was amplified in technical duplicates with SYBR Green I master mix (Roche, #04707516001) on a Light Cycler 480 II (Roche) cycling 55-times between 94°C, 60°C and 72°C with each temperature step running for 10 sec and switching at +4.8°C/sec and −2.5°C/sec. At the end qPCR reactions were heated from 65°C to 97°C with a gradual increase of 0.11°C/sec (melting curve) to ensure only fluorescence was collected from one specific amplicon. ChIP-qPCR results were based on absolute quantification using an eight-point standard curve of three-fold dilutions of ∼1% input DNA (**Supplementary Fig. 2d,g,i,j,n**). RT-qPCR results were normalized to the housekeeping gene *odc1* (Figs. 4h and 7d and **Supplementary Figs. 10d**) or *rpl8* (**Supplementary Fig. 8b**), and scaled relative to control embryos using the 2^−ΔΔCt^ method^66^. The threshold cycle (C_t_) was derived from the maximum acceleration of SYBR fluorescence (second derivative maximum method).

### ChIP-Seq library preparation

0.5 to 10 ng ChIP DNA or 5 ng input DNA were used to prepare single (only Tbx6 ChIP-Seq) or indexed paired-end libraries as previously described^65, 67–69^.

### Post-sequencing analysis of ChIP-Seq

Single reads of maximal 50 bases were processed using trim_galore v0.4.2 (Babraham Institute, UK) to trim off low-quality bases (default Phred score of 20, i.e. error probability was 0.01) and adapter contamination from the 3’ end. Processed reads were aligned to the *X. tropicalis* genome assembly v7.1 (and v9.1 for some ChIP-Seq data) running Bowtie2 v2.2.9^70^ with default settings (**Supplementary Table 1**). Alignments were converted to the HOMER’s tag density format^71^ with redundant reads being removed (makeTagDirectory -single -tbp 1 -unique -mapq 10 -fragLength 175 -totalReads all). Only uniquely aligned reads (i.e. MAPQ ≥10) were processed. We pooled all input alignments from various developmental stages. This created a comprehensive mappability profile that covered ∼400 million unique base pair positions. All chromatin profiles were position-adjusted and normalized to the effective total of 1 million aligned reads including multimappers (counts per million aligned reads, CPM). Agreeing biological replicates according to ENCODE guidelines^64^ were subsequently merged. For stage 10^+^ (early gastrula stage) β-catenin and Smad2 ChIP-Seq, external datasets^25, 72, 73^ were used as biological replicates. HOMER’s peak caller was used to identify transcription factor binding sites by virtue of ChIP-enriched read alignments (hereafter called peaks): findpeaks-style factor-minDist 175 -fragLength 175 -inputFragLength 175 -fdr 0.001 -gsize 1.435e9 -F 3 -L 1 -C 0.97. This means that both ChIP and input alignments were extended 3’ to 175 bp for the detection of significant (0.1% FDR) peaks being separated by ≥175 bp. The effective size of the *X. tropicalis* genome assembly v7.1 was set to 1.435 billion bp, an estimate obtained from the mappability profile. These peaks showed equal or higher tag density than the surrounding 10 kb, ≥3-fold more tags than the input and ≥0.97 unique tag positions relative to the expected number of tags. To detect focal RNAPII recruitment to pCRMs and avoid calling peaks within broad regions of RNAPII reflecting transcript elongation, the threshold of focal ratio and local enrichment within 10 kb was elevated to 0.6 and 3 (-L 3), respectively. To further eliminate any false positive peaks, we removed any peaks with <0.5 (TFs including signal mediators) or <1 (RNAPII) CPM and those falling into blacklisted regions showing equivocal mappability due to genome assembly errors, gaps or simple/tandem repeats. Regions of equivocal mappability were identified by a two-fold lower (poor) or three-fold higher (excessive) read coverage than the average detected in 400-bp windows sliding at 200-bp intervals through normalized ChIP input and DNase-digested naked genomic DNA. All identified regions ≤800 bp apart were subsequently merged. Gap coordinates were obtained from the Francis Crick mirror site of the UCSC genome browser (http://genomes.crick.ac.uk). Simple repeats were masked with RepeatMasker v4.0.6^74^ using the crossmatch search engine v1.090518 and the following settings: RepeatMasker-species “xenopus silurana tropicalis” -s -xsmall. Tandem repeats were masked with Jim Kent’s trfBig wrapper script of the Tandem Repeat Finder v4.09^75^ using the following settings: weight for match, 2; weight for mismatch, 7; delta, 7; matching probability, 80; indel probability, 10; minimal alignment score, 50; maximum period size, 2,000; and longest tandem repeat array (-l), 2 [million bp]. The multi-genome sequence conservation track (phastCons) for *X. tropicalis* genome assembly v9.1 was obtained from Xenbase^76^. The following eleven vertebrate species were used to evaluate sequence similarity: *X. tropicalis*, *X. laevis*, *Nanorana parkeri* (High Himalaya Frog), *Fugu rubripes* (Japanese Pufferfish), *Chrysemys picta* (Painted Turtle), *Gallus gallus* (Chicken), *Anolis carolinensis* (Green Anole lizard), *Monodelphis domestica* (Gray Short-tailed Opossum), *Canis lupus familiaris* (dog), mouse and human. pCRMs with a phastCons average ≥0.4 were considered ‘conserved’.

### Poly(A) RNA-Seq

Libraries were made from ∼1 µg total RNA by following the low-sample protocol of the TruSeq RNA sample preparation guide version 2 with a few modifications. First, 1 µl cDNA purified after second strand synthesis was quantified on a fluorometer using high-sensitivity reagents for double-stranded DNA (10 pg/µl to 100 ng/µl). By this stage, the yield was ∼10 ng. Second, the numbers of PCR cycles were adjusted to the detected yield of cDNA to avoid products of over-amplification such as chimera fragments: 7 (∼20 ng), 8 (∼10 ng), 9 (∼5 ng) and 12 (∼1 ng).

### RNA-Seq read alignment

Paired-end reads were aligned to the *X. tropicalis* genome assembly v7.1 using STAR v2.5.3a^77^ with default settings (**Supplementary Table 1**) and a revised version of gene models v7.2^50^ to improve mapping accuracy across splice junctions. The alignments were sorted by read name using the sort function of samtools v1.3.1^78^. Exon and intron counts (-t ‘exon;intron’) were extracted from unstranded (-s 0) alignment files using VERSE v0.1.5^79^ in featureCounts (default) mode (-z 0). Intron coordinates were adjusted to exclude any overlap with exon annotation. For visualization, genomic BAM files of biological replicates were merged using samtools v1.3.1 and converted to the bigWig format. These genome tracks were normalized to the wigsum of 1 billion excluding any reads with mapping quality <10 using the python script bam2wig.py from RSeQC v2.6.4^80^.

### Differential gene expression analysis

Differential expression analysis was performed with both raw exon and intron counts excluding those belonging to ribosomal and mitochondrial RNA using the Bioconductor/R package DESeq2 v1.16.1^81^. In an effort to find genes with consistent fold changes over time, p-values were generated according to a likelihood ratio test reflecting the probability of rejecting the reduced (∼ developmental stage) over the full (∼ developmental stage + condition) model. Resulting p-values were adjusted to obtain false discovery rates (FDR) according to the Benjamini-Hochburg procedure with thresholds on Cook’s distances and independent filtering being switched off. Equally, regional expression datasets^51^ without time series were subjected to likelihood ratio tests with reduced (∼ 1) and full (∼ condition) models for statistical analysis. Fold changes of intronic and exonic transcript levels were calculated for each developmental stage and condition using the mean of DESeq2-normalized read counts from biological replicates. Both intronic and exonic datasets were filtered for ≥10 DESeq2-normalized read counts detected at least at one developmental stage in all uninjected or DMSO-treated samples. Gene-specific fold changes were removed at developmental stages that yielded <10 normalized read counts in corresponding control samples. Next, the means of intronic and exonic fold changes were calculated across developmental stages. The whole dataset was confined to 3,687 genes for which at least 50% reductions (FDR ≤10%) in exonic (default) or intronic counts could be detected in α-amanitin-injected embryos. Regional expression was based on exonic read counts by default unless the intronic fold changes were significantly (FDR ≤10%) larger than the exonic fold changes (**Supplementary Table 9**).

### Analyzing ribosome footprinting and mass-spectrometry data

The ribosome footprinting reads were trimmed 5’ by 8 bases and 3’ by as many bases overlapping with the adapter sequence 5’-TCGTATGCCGTCTTCTGCTTG-3’ from the 5’ end. All trimmed reads between 27 to 32 bases in length were aligned first to ribosomal RNA as listed in the SILVA rRNA database^82^ using Bowtie v1.0.1^83^ with the following parameters: --seedlen 25 (seed length) --seedmms 1 (number of mismatches allowed in the seed) --un (unaligned reads were reported). All non-aligned (rRNA-depleted) reads were mapped to the gene model 6.0 of *X. laevis* using Tophat v2.0.10^84^ --no-novel-juncs (spliced reads must match splice junctions of gene model 6.0) --no-novel-indels (indel detetion disabled) --segment-length 15 (minimal length of read fragment to be aligned) –GTF v6.0.gff3 --prefilter-multihits (reads aligned first to whole genome to exclude reads aligning >10 times) --max-multihits 10. Alignment files were converted to the HOMER’s tag density format^71^ before retrieving reads for each CDS per kilobase and one million mapped reads (rpkm) using HOMER’s perl script analyzeRNA.pl. This read count table was merged with a published list of estimated protein concentrations (nM) in the *X. laevis* egg using mass-spectrometry^12^ (**Supplementary Table 4**). Chromatin-associated proteins regulating RNAPII-mediated transcription were filtered based on human (version 2.0, 09/2014) and Xenbase-released gene ontology (GO) associations. The genes associated with any of the following GO terms were verified with UniProt (UniProt Consortium, 2015) whether chromatin binding is supported by functional evidence: chromatin (GO:0000785), DNA binding (GO:0003677), DNA-templated regulation of transcription (GO:0006355), sequence-specific DNA binding and TF activity (GO:0003700) and DNA replication (GO:0006260). We separated specific TFs from all other chromatin binding proteins to form three categories: (1) sequence-specific DNA binding factors, (2) other RNAPII-regulating factors and (3) remaining genes, whereby genes associated with DNA repair (GO:0006281) and RNAPIII (GO:0006383) were moved to the third category.

### SPI-Based enrichment of small DNase-digested fragments (DNase-Seq)

In our hands, the high yolk content in vegetal blastomeres made it impossible to employ ATAC-Seq^11^ on whole *Xenopus* embryos before the onset of gastrulation. Therefore, DNase-Seq was adapted to early *Xenopus* embryos with a novel approach to select small DNase-digested DNA fragments. Ultracentrifugation- or gel electrophoresis-mediated size selection^85, 86^ was replaced by two rounds of solid phase immobilization (SPI) to remove long inaccessible DNA from short accessible DNA. Wide-bore pipette tips were used for the resuspensions and the transfers of samples from the second homogenization step until after SPI to minimize fragmentation of high-molecular DNA. Approximately 250 de-jellied mid-blastula embryos were collected in 2-ml round-bottom microcentrifuge tubes and homogenized in 2 ml ice-cold LB-DNase buffer (15 mM Tris-HCl pH 8.0, 15 mM NaCl, 60 mM KCl, 1 mM EDTA, 0.5 mM EGTA and 0.5 mM spermidine) supplemented with 0.05% Igepal CA-630. This lysate was left on ice for 3 min before centrifuging at 1,000 g (4°C) for 2 min. The supernatant was discarded without disrupting the pellet. The pellet was gently resuspended in 2 ml ice-cold LB-DNase buffer (without Igepal CA-630) before centrifuging again at 1,000 g (4°C) for 2 min. After discarding the supernatant, the pellet was resuspended at room temperature in 600 µl LB-DNase buffer (without Igepal CA-630) with 6 mM CaCl_2_. The sample was distributed equally to two 1.5-ml microcentrifuge tubes, one for probing chromatin accessibility with DNase and the other one for creating a reference profile of DNase-digested naked genomic DNA. Approximately 0.1 U recombinant DNase I (Roche, #04716728001) was added to one aliquot. Both samples were incubated at 37°C for 8 min before adding 300 µl STOP buffer (0.1% SDS, 100 mM NaCl, 100 mM EDTA and 50 mM Tris-HCl pH 8.0) including 80 µg RNase A (Thermo Fisher Scientific, #12091021), 333 nM spermine and 1 µM spermidine. The tubes were inverted gently to mix samples before incubating them at 55°C for 15 min. Next, 200 µg proteinase K (Thermo Fisher Scientific, #AM2548) were added and the tubes were inverted gently to mix the samples again. After 2 h at 55°C, the digests were transferred to pre-spun 1.5-ml Phase Lock Gel Heavy microcentrifuge tubes (VWR). 600 µl phenol:chloroform:isoamylalcohol (25:24:1, pH 7.9) were added to the digests. The tubes were shaken gently and then centrifuged at 1,500 g (room temperature) for 4 min. The top phase was transferred to a fresh 2-ml microcentrifuge tube and mixed with 60 µl 3 M sodium acetate (pH 5.2) and 1.2 ml absolute ethanol to precipitate the genomic DNA. The precipitation was stored at −20°C overnight and centrifuged at 16,000 g (4°C) for 30 min. The supernatant was discarded and the DNA pellet washed by adding 500 µl ice-cold 80% ethanol and centrifuging at 16,000 g (4°C) for 3 min. The ethanol was discarded and the DNA pellet was air-dried at room temperature for 10 min. After that, the DNA pellet was left on ice to dissolve in 27 µl elution buffer (10 mM Tris-HCl pH 8.5) for 20 min. To remove any residual RNA, 10 µg RNase A were added to the DNA. 1 µl was used to determine the DNA fragment size distribution by gel electrophoresis. On a 0.6% TAE agarose gel, a smear of very high molecular DNA was visible as expected from previous DNase experiments^85, 87^. Genomic DNA to the amount of 20 untreated mid-blastula embryos was digested with 0.3 U DNase I at 37°C for 5 min following the same steps as described above, which generated a low-molecular smear of DNA fragments. Next, 70 µl AMPure XP beads (Beckman Coulter, #A63880) per DNase sample were equilibrated to room temperature for 10 min. 22.5 µl AMPure XP beads (0.9x of the volume) were added to the DNA sample without pipetting up and down achieving a final polyethylene glycol concentration of ∼9.5%. After 3 min, by which time high-molecular DNA causes beads to coalesce, the tubes were clipped into a magnetic stand for microcentrifuge tubes. After 3 min or until the beads were separated from the supernatant, the latter was transferred to a 96-well microplate (350-µl round wells with V-shaped bases). 47.5 µl elution buffer and 43 µl AMPure XP beads were added sequentially and mixed gently by slowly pipetting up and down. After 3 min, the plate was transferred to a magnetic stand for 96-well plates. Once the beads have completely separated from the suspension, the supernatant was transferred to a pre-spun 1.5-ml Phase Lock Gel Heavy tube. 162 µl elution buffer and 300 µl phenol:chloroform:isoamylalcohol (25:24:1, pH 7.9) were added. The tubes were shaken gently and then centrifuged at 1,500 g (room temperature) for 4 min. The top phase was transferred to a 1.5-ml low-retention microcentrifuge tube and mixed with 30 µl 3 M sodium acetate (pH 5.2) and 900 µl absolute ethanol to precipitate the DNA as outlined above. The DNA pellet was dissolved in 12 µl elution buffer. The concentrations of the DNA samples were determined on a fluorometer using high sensitivity reagents for dsDNA (10 pg/µl to 100 ng/µl). Libraries were generated as previously outlined^65^ except that all cleaning steps were executed with 0.2x more AMPure XP beads.

### Post-sequencing analysis of DNase-Seq

Single and paired-end reads of maximal 50 bases were processed using trim_galore v0.4.2 from the Babraham Institute (UK) to trim off low-quality bases (default Phred score of 20, i.e. error probability was 0.01) and adapter contamination from the 3’ end. Processed reads were aligned to the *X. tropicalis* genome v7.1 and v9.1 using Bowtie2 v2.2.9^70^ with default settings apart from -X (fragment length), which was reduced to 250 bp for paired-end reads. Alignments were sorted by genomic coordinates and only those with a quality score of ≥10 were retained using samtools v1.3.1^78^. Duplicates were removed using Picard (MarkDuplicates) from the Broad Institute (USA). Paired-end alignments were dissociated using hex flags (-f 0×40 or 0×80) of samtools view. Single alignments were converted to HOMER’s tag density format^71^ (makeTagDirectory - single -unique -fragLength 100 -totalReads all). DNase hypersensitive sites were identified using the following or otherwise default parameters of HOMER’s peak calling: findpeaks -style factor -minDist 100 -fragLength 100 -inputFragLength 100 -fdr 0.001 - gsize 1.435e9 -F 3 -L 1 -C 0.97. This means that alignments of DNase-digested chromatin fragments (or naked genomic DNA fragments considered here as ‘input’) were extended 3’ to 100 bp from the DNase cleavage site to detect significant (0.1% FDR) DNase-enriched read alignments (hereafter called peaks) being separated by ≥100 bp. The effective size of the *X. tropicalis* genome assembly v7.1 was set to 1.435 billion bp, an estimate obtained from the mappability of ChIP input reads. These peaks showed equal or higher tag density than the surrounding 10 kb, at least three-fold more tags than the input and ≥0.97 unique tag positions relative to the expected number of tags. Peaks falling into blacklisted regions (see Post-sequencing analysis of ChIP-Seq) were removed. The profiles of DNase-probed chromatin accessibility were position-adjusted and normalized to the effective total of 1 million aligned reads including multimappers. Genomic coordinates of accessible pCRMs are listed in **Supplementary Tables 3** and **10** (mPouV/Sox3 LOF experiment).

### Next-generation capture-C

About 500 mid-blastula embryos per condition were collected in 9-ml capped glass vials and briefly washed once with 1% MMR. The embryos were then fixed with 1% formaldehyde (Sigma, #F8775) in 9 ml 1% MMR at room temperature for 40 min. The fixation reaction was terminated by briefly rinsing the embryos three times with ice-cold 1% MMR. The embryos were aliquoted into 2-ml round-bottom microcentrifuge tubes in batches of ∼250 embryos (filling the volume of ∼250 µl water). The supernatant was removed before equilibrating embryos in 250 µl ice-cold HEG buffer (50 mM HEPES pH 7.5, 1 mM EDTA and 20% glycerol). Once the embryos settled to the bottom of the tube as much liquid as possible was discarded. The aliquots were snap-frozen in liquid nitrogen and stored at −80°C.

For each chromatin conformation capture (3C) experiment, 10 ml chromatin extraction buffer CEB-3C (10 mM Tris-HCl pH 8.0, 150 mM NaCl, 1 mM EDTA, 1% Igepal CA-630, 0.25% sodium deoxycholate and 0.2% SDS) were supplemented with a mini protease inhibitor tablet (Roche, #11873580001) and 0.05 mM DTT (CEB-3C*). Both aliquots of ∼250 embryos of each condition were homogenized in 2 ml ice-cold CEB-3C*. The tubes were kept on ice for 5 min before centrifuging at 1,000 g (4°C) for 2 min. The supernatants were discarded and the pellet resuspended in 0.5 ml ice-cold CEB-3C*. The resuspensions of each condition were pooled and then equally divided to two 50-ml conical tubes and each filled up with CEB-3C (without protease inhibitors and DTT) to 50 ml. The embryonic extracts were incubated at 37°C for 1 h in a hybridization oven with the tubes rotating inside a hybridization bottle. The tubes were then centrifuged at 1,000 g (room temperature) for 5 min. The supernatants were discarded and the pellets resuspended in 50 ml double-distilled water. The tubes were centrifuged again at 1,000 g (room temperature) for 5 min. The supernatants were discarded and each pellet was resuspended in 300 µl double-distilled water. The resuspensions containing cross-linked nuclei of each condition were pooled in a 1.5-ml microcentrifuge tube. The samples were then digested with 400 U *Dpn*II (NEB, #R0543) in a total volume of 800 µl containing 1x *Dpn*II reaction buffer. The digest was incubated overnight in a thermomixer set to 37°C and 1,400 rpm. Of note, any residual yolk platelets in the solution did not interfere with the restriction digest. The next day, the digest was supplemented with 200 U *Dpn*II and incubated at 37°C and 1,400 rpm for another 6-8 h. After that, *Dpn*II was heat-inactivated at 65°C for 20 min.

To verify the degree of chromatin digestion, an aliquot of 50 µl from each condition was transferred into separate 1.5-ml microcentrifuge tubes and processed as follows: First, any remaining RNA was degraded by incubating the aliquot with 20 µg RNase A (Thermo Fisher Scientific, #12091021) at 37°C for 30 min. Second, proteins were digested by incubating the aliquot mixed with 50 µl SDS elution buffer (50 mM Tris pH 8.0, 1 mM EDTA and 1% SDS) and 20 µg proteinase K (Thermo Fisher Scientific, #AM2548) at 65°C in a hybridization oven overnight. Third, the DNA was purified and ethanol precipitated: 300 µl TE pH 8.0 were added to the digest before transferring it to pre-spun 1.5-ml Phase Lock Gel Heavy microcentrifuge tubes (VWR). The diluted digests were mixed vigorously with 400 µl (1 volume) phenol:chloroform:isoamylalcohol (25:24:1, pH 7.9) before centrifuging at 16,000 g (room temperature) for 1 min. The top (aqueous) phases were transferred into 1.5-ml microcentrifuge tubes and mixed with 40 µl sodium acetate (pH 5.2) and 800 µl absolute ethanol and 15 µg GlycoBlue (Thermo Fisher Scientific, #AM9516). The DNA precipitations were kept at −80°C for 30 min before centrifuging at 16,000 g (4°C) for 30 min. The supernatants were discarded without disturbing the DNA pellets which were subsequently washed once with 400 µl ice-cold 80% ethanol. The tubes were centrifuged at 16,000 g (4°C) for 30 min before discarding the supernatants. The DNA pellets were air-dried for 10 min and resuspended in 10 µl elution buffer (10 mM Tris-HCl pH 8.5). The DNA concentrations were determined with a fluorometer and the DNA fragment distributions were visualized on a 0.6% TAE agarose gel. The digestion of cross-linked chromatin with *Dpn*II caused extensive fragmentation of the genomic DNA such that no high-molecular DNA fragment bands (>10 kb) were visible.

The chromatin digests were processed for proximity ligation by adding 240 U T4 DNA ligase (Thermo Fisher Scientific, #EL0013) in a total volume of 1.2 ml containing 1x T4 DNA ligase buffer including 5 mM ATP. The ligation reactions were incubated at 16°C and 1,400 rpm for ≥16 h before centrifuging at 16,000 g (4°C) for 1 min. The supernatants were discarded. 200 µl elution buffer and 40 µg RNase A were added to the pellets to degrade any residual RNA at 37°C and 1,400 rpm for 30 min. Proteins were digested by adding 200 µl SDS elution buffer and 160 µg proteinase K and incubating at 65°C in a hybridization oven overnight. The digest was transferred to pre-spun 1.5-ml Phase Lock Gel Heavy tubes and processed as described in the previous paragraph to purify and precipitate ligated genomic DNA. The air-dried DNA pellet was dissolved in 20 µl elution buffer and quality controlled as above. Proximity ligation of *Dpn*II-digested chromatin fragments massively increased the size of DNA fragments to the range of high molecular weight (>10 kb) representing the 3C library. The concentration was measured on a fluorometer with broad range concentration (5-500 ng/µl) reagents.

About 10-15 µg of the 3C libraries were diluted with elution buffer to the total volume of 130 µl and transferred to a designated glass vial (microTUBE) for isothermal focused sonication (Covaris). The following settings of the focused ultrasonicator Covaris S220 were used to shear the libraries to an average DNA fragment length of ∼200 bp: duty cycle, 10%; intensity, 5; cycles per burst, 200; duration, 60 sec in frequency sweeping mode. Sonication was run with 6 cycles in batch mode. The degree of DNA fragmentation was verified by gel electrophoresis and the DNA concentration was measured on a fluorometer with broad range concentration reagents.

Approximately 2-4 µg of the DNA fragments were converted into next-generation paired-end libraries using the enzymes of the KAPA Hyper Prep Kit (Kapa Biosystems, #KR0961) and TruSeq adaptors (Illumina). The end-repair and A-tailing reactions were set up according to the manufacturer’s instructions and incubated at 20°C for 1 h followed by 30 min at 65°C. End-repaired and A-tailed DNA fragments were ligated to 150 pmol TruSeq adaptors of index 6 and 12, respectively, at 20°C for 1 h. The DNA was purified using 0.8x volume of AMPure XP beads (Beckman Coulter, #A63880) and amplified in a total volume of 100 µl using 50 pmol of each TruSeq PCR primer, KAPA HiFi HotStart ReadyMix (Kapa Biosystems, #KK2602) and the following PCR conditions: 98°C for 45 sec followed by 3 cycles (98°C for 15 sec, 98°C for 30 sec and 72°C for 30 sec) and a final elongation step of 1 min at 72°C. The lid temperature was set to 105°C. The PCR reactions were cleaned up with 100 µl AMPure XP beads and eluted in 14 µl elution buffer. The integrity of the library DNA was verified on a chip-based capillary electrophoresis system, an 8% TBE polyacrylamide gel or a 2% TAE agarose gel. DNA concentration was measured on a fluorometer with broad range concentration reagents. The DNA yield was 2-4 µg.

To capture the genomic regions of interest (viewpoints), 5’-biotinylated oligonucleotides of 120 bases (hereafter called probes) (**Supplementary Table 10**) were designed for each viewpoint as follows: The viewpoints were *Dpn*II fragments of 300-3,300 bp which were overlapping gene promoters or located <1 kb from them. Each probe matched the terminal sequence of the same DNA fragment strand including the *Dpn*II restriction site GATC. The probe sequence was examined using BLAT to check whether it is unique and did not partially match any other genomic regions. Furthermore, CENSOR v4.2.29^88^ and RepeatMasker v4.0.6^74^ programs were run to discard any probes that contained repeats. Because of these design restrictions, six of the selected 30 gene promoters were captured with only one probe. All the probes were purchased as desalted oligomers (4 nmol) from IDT (ultramer technology) and reconstituted in molecular-grade water to 10 µM. The probes were mixed in equimolar quantities and diluted to 2.9 nM such that 4.5 µl contained 13 fmol of oligomers.

The capture was performed with the SeqCap Hybridization and Wash Kit (Roche, #05634253001) and SeqCap HE-Oligo Kit A (Roche, #06777287001) as follows. Exactly 1 µg of each TruSeq library with index 6 and 12 was mixed with 1 nmol TruSeq universal blocking oligonucleotide and 0.5 nmol blocking oligonucleotides specific to TruSeq index 6 and 12 in a 1.5-ml low-retention microcentrifuge tube. Of note, Cot-1 DNA commonly used to mask repetitive DNA proved to be unnecessary here in reducing non-specific hybridization. The mixture of library and blocking oligonucleotides was dried in a vacuum centrifuge before adding 7.5 µl 2x hybridization buffer (vial 5) and 3 µl hybridization component A (vial 6). This mixture was vortexed for 10 sec, centrifuged at 16,000 g for 10 sec and incubated at 95°C for 10 min. In the meantime, 4.5 µl (13 fmol) of the equimolar probe mixture were transferred to a PCR tube and incubated at 47°C in a PCR machine with the volume and the lid temperature set to 15 µl and 57°C, respectively. Upon denaturation, the libraries and blocking oligonucleotides were added to the probes without removing either tubes from the heating block or the PCR machine. The hybridization mixture was incubated at 47°C for 64-72 h. The wash buffers were prepared according to the manufacturer’s instructions to make 1x working solutions for one single capture experiment. The stringent wash buffer and wash buffer I were pre-warmed to 47°C and 50 µl M270 streptavidin-conjugated magnetic beads (Thermo Fisher Scientific, #65305) were transferred into a 1.5-ml low-retention microcentrifuge tube to let them equilibrate to room temperature for 10 min. The beads were washed twice with 200 µl bead wash buffer. Immediately after the final wash, the hybridization mixture was directly added to the beads and vortexed for 10 sec. The sample was incubated for 45 min in a thermomixer set to 47°C and 1,100 rpm. The beads were washed by adding 100 µl pre-warmed wash buffer I and vortexing for 10 sec. The tube was placed into a magnetic rack. Once the liquid was clear, the supernatant was discarded, and 200 µl pre-warmed stringent wash buffer were added to the beads. In an effort to avoid any temperature drop, it is important to work quickly according to the manufacturer’s instructions. The beads in stringent wash buffer were incubated in a thermomixer set to 47°C and 1,100 rpm for 5 min. The wash with pre-warmed stringent wash buffer was repeated once. Next, 200 µl pre-warmed wash buffer I was added to the beads and vortexed at 1,400 rpm (room temperature) for 2 min. After removing the respective wash buffer, beads were vortexed at 1,400 rpm in 200 µl wash buffer II for 1 min and 200 µl wash buffer III for 30 sec. After the final the final wash, as much liquid as possible was removed and the beads were resuspended in 40 µl molecular-grade water.

The captured DNA was directly amplified from the beads using KAPA HiFi HotStart master mix and 25 pmol of each TruSeq PCR primer in two separate 50-µl PCR reactions. The PCR conditions were the same as outlined above except that 14 cycles were used for amplification. The PCR reactions were pooled and the DNA was purified using 100 µl AMPure XP beads. The DNA was eluted with 11 µl elution buffer. The eluted DNA was subjected to another round of probe-mediated capture with the hybridization timespan reduced to 24 h. Furthermore, after washing the beads, the captured fragments were amplified using only 10 cycles of PCR.

### Post-sequencing analysis of next-generation capture-C data

The analysis was carried out in accordance with the original description of the method^89^ with some modifications. Paired-end reads were processed using trim_galore v0.4.2 from the Babraham Institute (UK) to trim off low-quality bases (default Phred score of 20, i.e. error probability was 0.01) and adapter contamination from the 3’ end. Only complete mate pairs were processed further to reconstruct single reads from overlapping paired-end sequences using FLASH v1.2.11^53^ with interleaved output settings for non-extended reads (flash --interleaved-output --max-overlap 150). FLASH reads were split in *silico* at *Dpn*II restriction sites using a designated perl script (https://github.com/telenius/captureC/releases) before aligning them to the *X. tropicalis* genome assembly 7.1 using Bowtie2 v2.2.9^70^. The alignment was run with default settings and one thread only to keep the order of the reads. The view function of samtools v1.3.1^78^ was used to retain alignments with a quality score of ≥10. The alignments were analyzed further using a suite of perl scripts (https://github.com/Hughes-Genome-Group/captureC/releases) modified to process both chromosome and scaffold coordinates. Viewpoint coordinates included a 1-kb proximity exclusion range. Restriction fragments were classified as capture, proximity-exclusion or reporter. PCR duplicates were removed. The interaction map was based on the number of unique paired-end reads per restriction fragment. Windows of 2 kb incrementing by 200 bp were used to consolidate interactions, which were normalized to 10,000 interactions per viewpoint. Promoter contacts are listed in **Supplementary Table 12**.

### Analysis of genomic profiles, DNA motifs and super-enhancers

Unless otherwise stated, Spearman correlation (R_s_), relative tag density (meta-plots), principal component analysis (PCAs) and enrichment of DNA motifs were determined using a limited number of detected peaks per chromatin profile: top 2,000 peaks (Figs. 3c-g and 4c and **Supplementary Figs. 5, 6** and **7d**), top 5,000 peaks (**Supplementary Fig. 1b**) and top 10,000 peaks (**Figs. 4b,e** and **5c** and **Supplementary Fig. 7a-c**). R_s_ and PCA were calculated using DiffBind v2.4.8^90^ in R (**Figs. 1d**, **3d,e** and **4b** and **Supplementary Figs. 5a**, **7a,b** and **12a**). Z-scores of motif enrichments at RNAPII peaks were calculated using ISMARA v1.2.2^91^ (https://ismara.unibas.ch/fcgi/mara) in expert mode (**Fig. 1d**). Heat maps of relative tag densities were hierarchically clustered (Ward’s method) using the R package seriation^92^ except for those in **Figs. 4c** and **8d**, where genomic coordinates were sorted upon significant changes in DNA occupancy or chromatin accessibility (see Figure legends). In **Supplementary Fig. 6**, cluster leaves were additionally ordered with the Gruvaeus-Wainer’s algorithm. The matrices of normalized tag densities, p-values and motif instances were generated with HOMER as follows: annotatePeaks.pl -size 2000 (or 200 for **Supplementary Fig. 6**) -hist 25 (bin size) -ghist. In heat maps, all genomic profiles were scaled to the means of the highest bin values of the top 100 (or 1,000 in **Figs. 4c** and **8d**) peaks. In addition, to visualize low RNAPII binding and RNA expression, their top means were divided by 5 or 500, respectively (**Fig. 8d**). Super-enhancers were assembled by stitching together peaks that are ≤25 kb apart: findPeaks -style super -fragLength 175 -inputFragLength 175 –fdr 0.001 -gsize 1.435e9 -F 3 -L 1 -localSize 10000 -C 0.97 -superSlope 1 -minDist 25000. Subsequently, we combined these super-enhancers according to the developmental stage of profiling (see **Supplementary Table 6** for genomic coordinates). Super-enhancers ≤5 kb from TSS apart were associated with zygotic genes defined by RNAPII-mediated transcription elongation^52^.

### Analysis of enriched gene ontology (GO) terms

Over-represented GO terms were found by applying hypergeometric tests of the Bioconductor/R package GOstats v2.42.0^93^ on gene lists. The process was also supported by the Bioconductor/R packages GSEABase v1.38.1^94^ and GO.db v3.4.1^95^. The gene universe was associated with GO terms by means of BLAST2GO^96^ as previously outlined^67, 97^.

### Differential analysis of chromatin features

The significance of differential chromatin accessibility between control and mPouV/Sox3 LOF embryos was assessed at pCRMs using the Bioconductor/R package DiffBind v2.4.8^90^ and the DESeq2^81^ algorithm. The size of all pCRMs was fixed to 500 bp. Only the very 5’ end of each aligned DNase-Seq read, which represents the DNase cleavage site, was considered for this analysis. Read counts from mapping DNase-digested naked genomic DNA were subtracted from chromatin accessibility readings at each pCRMs. Any changes to DNase-mediated cleavages at pCRMs with FDR ≤10% were determined as statistically significant (**Fig. 8b**). For visualization, the orientations of pCRMs were aligned to the direction of transcription of the closest zygotic gene. Matrices of normalized and raw tag counts were generated at pCRMs for various chromatin features and RNA (**Fig. 8b**). Their rows were sorted according to the statistical significance of differential chromatin accessibility. Raw tag counts in sliding (+200 bp) windows of 400 bp (accessibility, H3K4me1, RNAPII, Smad2 and β-catenin) or 2 kb (RNA and promoter contacts) across the genome were processed to estimate p-values of chromatin changes (pΔ) using DESeq2 v1.16.1^81^. Read count dispersions were locally fitted. The shrinkage of log_2_ fold changes, threshold on Cook’s distances and independent filtering were all switched off. For the comparison of local interactions with promoters at affected and unaffected pCRMs, the size of the pCRMs was widened to 1.2 kb and filtered for ≥5 reporter fragments (**Fig. 9c**). Normalized line and bar graph tracks of control and mPouV/Sox3 LOF embryos were superimposed in the IGV genome browser v2.3.92^98^ (**Figs. 9f** and **10a,b** and **Supplementary Figs. 13** and **14**).

### Software for computational analysis and graphical illustration

Post-alignment analysis and graphical illustration were performed using samtools v1.3.1^78^, bedtools v2.25.0^99^, HOMER v4.8.3^71^ and tools in R v3.4.1 / Bioconductor v3.5, Perl v5.18.2 or Python v2.7.12.

### Quantification and statistical analysis

No statistical method was used for determining sample size. We followed the literature to select the appropriate sample size. The experiments were not randomized. Due to the nature of experiments, the authors were not blinded to group allocation during data collection and analysis. Only viable embryos and embryonic tissues were included in the analysis. Due to space constraints, data from the trials of establishing and optimizing protocols (ChIP-Seq, DNase-Seq and next-generation capture-C) were excluded. Frequencies of shown morphological phenotypes and WMISH patterns are included in every image. Significances of normally or non-normally distributed data points were calculated using a paired Student’s t-test (one-tailed or two-tailed) or paired Wilcoxon test (two-tailed), respectively. The effect size r was estimated from the standard normal deviate of the Wilcoxon p-value (p) as previously described^100^, r_effect_=Z/sqrt(N), where Z=qnorm(1-p/2) is the standardized Z-score and N is the number of observations. Significance of motif enrichments or GO terms are based on hypergeometric tests. For RNA-Seq, biological triplicates were used to account for transcriptional variability between clutches. Each LOF experiment has its own control embryos collected in parallel from the same mothers: exp. #1 (α-amanitin), uninjected embryos; exp. #2 (BMP or Nodal LOF), DMSO-treated embryos; exp. #3 (Wnt LOF), uninjected embryos; exp. #4 (mPou5f3/Sox3 LOF), uninjected embryos; exp. #5 (mVegT LOF), uninjected embryos. The gene expressions of control MO-injected embryos of exp. #2 and #5 were normalized to their corresponding uninjected embryos. The mean of these normalizations and conservative FDR estimations (i.e., higher FDR of the two likelihood ratio tests) were used for the comparison with LOF conditions. RNA-Seq libraries from each experiment were generated simultaneously to mitigate any batch effects. The significance of transcriptome-wide differential expression was adjusted for multiple comparisons using the Benjamini-Hochburg procedure.

### External datasets

The following external datasets were used for this study: 5mC at stage 9, and ChIP for H3K4me3 and H3K27me3 at stage 9 (GSE67974), β-catenin ChIP at stage 10^+^ (GSE72657), Smad2 ChIP at stage 10^+^ (GSE30146 and GSE53654), Foxh1 ChIP at stage 8 and 10^+^ (GSE85273 and GSE53654), total RNA recorded at 30-min intervals from unfertilized eggs to 23.5 hpf (GSE65785), ribosome footprinting from stage 3, 9 and 12^+^ (GSE52809), and regional expression along the animal-vegetal and dorso-ventral axis at stage 10^+^ (GSE81458).

### Data and code availability

Programming code and some intermediate data underlying Figs. 1d-f, 2b, 3c-g, 4b,c,e, 5c, 6e,f, 7a,b,e, 8b-g and 9c, Supplementary Figs. 1a,b,d, 5a,b, 7a-d, 9, 10a,b, 11a,b, 12a and 15-17 and Supplementary Tables 9, 11 and 12 are available on GitHub at https://github.com/gegentsch/SignalCompetence.

## Notes

#### Summary of Updates

We have responded to the comments of the reviewers and revised our manuscript accordingly. The revised manuscript contains 10 main figures, 17 supplementary figures, 12 supplementary tables and 1 supplementary movie. Programming code and intermediate files have been updated and are now available on GitHub at https://github.com/gegentsch/SignalCompetence. The raw sequencing and processed data files have also been made publicly available in the GEO database (www.ncbi.nlm.nih.gov/geo) under accession number GSE113186.

